# Unraveling Phylogenetic Relationships, Reticulate Evolution, and Genome Composition of Polyploid Plant Complexes by Rad-Seq and Hyb-Seq

**DOI:** 10.1101/2021.08.30.458250

**Authors:** Kevin Karbstein, Salvatore Tomasello, Ladislav Hodač, Natascha Wagner, Pia Marinček, Birthe Hilkka Barke, Claudia Pätzold, Elvira Hörandl

## Abstract

Complex genome evolution of young polyploid complexes is poorly understood. Besides challenges caused by hybridization, polyploidization, and incomplete lineage sorting, bioinformatic analyses are often exacerbated by missing information on progenitors, ploidy, and reproduction modes. By using a comprehensive, self-developed bioinformatic pipeline integrating phylogenetic, structure, network, and SNP-origin analyses, we for the first time unraveled polyploid phylogenetic relationships and genome evolution within the large Eurasian *Ranunculus auricomus* species complex comprising more than 840 taxa. Our results rely on 97,312 genomic RAD-Seq loci, target enrichment of 576 nuclear genes (48 phased), and 71 plastid regions (Hyb-Seq; OMICS-data) derived from the 75 most widespread polyploid apomictic taxa and four di- and one tetraploid potential sexual progenitor species. Phylogenetic tree and structure analyses consistently showed 3–5 supported polyploid groups, each containing sexual progenitor species. In total, analyses revealed four diploid sexual progenitors and a one unknown, probably extinct progenitor, contributing to the genome composition of *R. auricomus* polyploids. Phylogenetic network, structure, and SNP-origin analyses based on RAD-Seq loci and phased nuclear genes completed by plastid data demonstrated predominantly allopolyploid origins, each involving 2–3 different diploid sexual subgenomes. Allotetraploid genomes were characterized by subgenome dominance and large proportions of interspecific, non-hybrid SNPs, indicating an enormous degree of post-origin evolution (i.e., Mendelian segregation of the diploid hybrid generations, back-crossings, and gene flow due to facultative sexuality of apomicts), but only low proportions of lineage-specific SNPs. The *R. auricomus* model system is the first large European polyploid species complex studied with reduced representation OMICS data. Our bioinformatic pipeline underlines the importance of combining different approaches and datasets to successfully unveil how reticulate evolution and post-origin processes shape the diversity of polyploid plant complexes.

## Introduction

Polyploidy, the presence of two or more full genomic complements (whole genome duplication), occurs across the tree of life (Otto and Whitton 2000; Van De Peer et al. 2017; Rothfels 2021). Whole-genome duplications have been observed in seed plants, and in several lineages of animals (mainly fish and amphibians), fungi, and protists (Mable et al. 2011; Van De Peer et al. 2017; Blischak et al. 2018). Polyploid cells and tissues occur throughout nature (also in humans) and are regarded as a cellular strategy for higher stress tolerance (Schoenfelder and Fox 2015; Fox et al. 2020).

All flowering plants are ancient polyploids, as at least one polyploidization event occurred in their common ancestor, and several additional ones in various lineages (Soltis and Soltis 2016; Van de Peer et al. 2017; Leebens-Mack et al. 2019). Neopolyploid formation for flowering plants is estimated to range between 30–70% of species and to cause upshifts of diversification rates in young polyploid complexes (Wood et al. 2009; Soltis et al. 2015; Landis et al. 2018). Key innovations in flowering plants have been hypothesized to be connected to polyploidy, for example, the carpel, double fertilization, and vessel elements (Soltis et al. 2015; Soltis and Soltis 2016; Leebens-Mack et al. 2019). In addition to its evolutionary significance, important crop plants are natural polyploids (e.g., wheat, potato, strawberry, coffee, cotton), and their evolution has often been exploited for agricultural purposes (Gordon et al. 2020).

The presence of multiple gene copies in polyploids allows for gene neo- and subfunctionalizations, epigenetic changes, and consequently a differential expression of homeologous genes (Comai 2005; Blischak et al. 2018). Polyploidy provides larger physiological and phenotypic flexibility to respond to different environmental conditions (Hörandl 2006; Marchant et al. 2016; Van De Peer et al. 2017; Karbstein et al. 2021), which facilitates colonization of various ecosystems (Te Beest et al. 2011; Rice et al. 2019; Fox et al. 2020; Meudt et al. 2021).

Genome evolution of polyploid lineages is complex and not only shaped by evolutionary origin and the genomic contributions of progenitors, but also by post-origin processes, resulting in a mosaic-like genome structure with parental, additive, and novel features (Soltis et al. 2015). Different polyploid formation types influence genome evolution: Autopolyploids arise within a species (tree-like evolution), whereas allopolyploids are formed by hybridization between different species/lineages followed by polyploidization (network-like evolution; Comai 2005; Wendel 2015; Blischak et al. 2018). Consequently, autopolyploids contain genetically similar subgenomes whereas genomes of allopolyploids are composed of previously diverged subgenomes. Allopolyploidization is considered particularly likely to create biotypes with novel genomic features (Abbott et al. 2013; Van de Peer et al. 2020; Rothfels 2021). For instance, allopolyploids showed higher degrees of genomic, transcriptomic, and epigenetic changes than autopolyploids (Comai 2005; Chen et al. 2007; Wendel 2015; Soltis et al. 2015; Spoelhof et al. 2017).

After evolutionary origin, the genome structure of neopolyploids is fluid over evolutionary time scales, and genomes revert to a functionally diploid state (Soltis et al. 2015; Van De Peer et al. 2017). At the beginning of this process, various mechanisms influence polyploid genomes. Expression bias due to epigenetic changes and homeologous gene loss (biased fractionation) after polyploidization can cause subgenome dominance (Soltis et al. 2015; Wendel 2015; Blischak et al. 2018; Alger and Edger 2020). Moreover, Mendelian segregation in the first diploid hybrid generations before polyploidization, and/or backcrossing of polyploids to their sympatric progenitors might distort the original subgenome contributions (Barke et al. 2018; Hodač et al. 2018; Wagner et al. 2020). Gene flow between polyploid lineages further influences genome structure (Melichárková et al. 2020).

In plants, polyploidization and/or hybridization are frequently connected to apomixis, i.e., reproduction via asexually-formed seeds (Asker and Jerling 1992; Brukhin et al. 2018; Hojsgaard and Hörandl 2019). Noteworthy, not all neopolyploids are apomicts (e.g., Masci et al. 1994), and not all apomicts are polyploids (e.g., Brukhin et al. 2018). Apomixis is usually facultative, and residual sexuality allows for backcrossing to progenitors and intercrossing of polyploids, resulting in huge networks of hundreds to thousands hybridogenetic lineages (Fig. 1). Such complexes occur in many abundant plant genera, e.g., dandelions (*Taraxacum*), hawkweeds (*Hieracium* s.l.), brambles (*Rubus*), and *Citrus*. With higher ploidy levels and/or time, these lineages are expected to become fixed, and mutations remain as the only source of genetic variation (Grant 1981; Coyne and Orr 2004; Fig. 1). With reduced recombination, heterozygosity in allopolyploids is additionally increased by allelic sequence divergence in asexual lineages (Meselson effect; Welch and Meselson 2000; Pellino et al. 2013). Studies using genome-wide data showed that heterozygosity significantly increased with higher ploidy levels (Mohammadin et al. 2018; Karbstein et al. 2021). In general, heterozygosity has several benefits, such as novel genetic combinations, heterosis, buffering effects of deleterious mutations, or changes in secondary metabolites (Comai 2005; Qiu et al. 2020). Increased heterozygosity is considered an important factor for the spreading of polyploids towards more variable climatic conditions (Hörandl 2006; Rice et al. 2019; Karbstein et al. 2021).

**Fig. 1.**
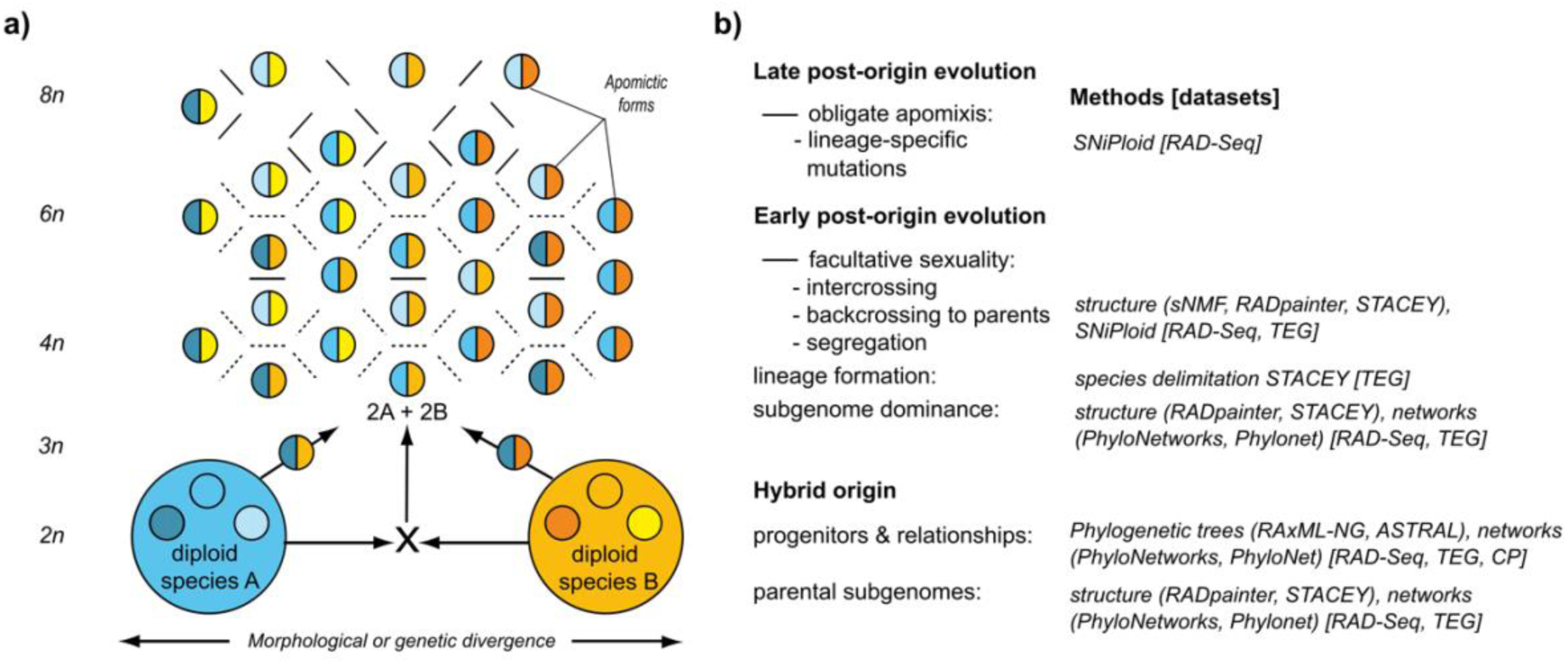
Evolutionary processes in young polyploid species complexes and methods to address these processes (a) Evolution of an apomictic polyploid complex from two sexual progenitor species and evolution of lineages after origin (redrawn and modified after Babcock and Stebbins 1938; see also Grant 1981 and Coyne and Orr 2004 for modern interpretations). (b) description of respective evolutionary processes and the corresponding analytical methods and pipelines applied here; CP = chloroplast (plastid) regions, RAD-Seq = RAD-Seq loci, TEG = target enriched nuclear genes. For a detailed scheme of bioinformatic pipelines see Fig. 3.

Despite the evolutionary, ecological, and economical importance of polyploidy, the understanding of phylogenetic relationships, genome diversity and evolution of fast-evolving, young polyploid species complexes remains limited. Traditional sequencing markers from organellar DNA were insufficient for reconstructing reticulate relationships in polyploid complexes because of uniparental inheritance (Rothfels 2021). Nuclear markers from single regions (e.g., ribosomal DNA) are biased by a strong marker-specific evolution (e.g., Zarrei et al. 2014; Fehrer et al. 2021). Historically, polyploids were thus often avoided or dropped in phylogenetic studies (Freyman et al. 2020; Rothfels 2021). Already the sexual progenitors of polyploid complexes are often characterized by low genetic divergence, incomplete lineage sorting (ILS), gene flow, and partial hybridogenic origins (Hörandl 2018; Pease et al. 2018; Wagner et al. 2019; Karbstein et al. 2020a, 2020b). The use of ‘OMICS’-data provides orders of magnitude more information compared with traditional genetic markers (Harrison and Kidner 2011; Soltis et al. 2013). OMICS-data have proven effective at resolving diploid (and few polyploid) phylogenetic relationships of species that diversified more than 30 or even less than 0.1 Ma (Pellino et al. 2013; Hipp et al. 2014; Carter et al. 2019; Gordon et al. 2020; Karbstein et al. 2020b; Tomasello et al. 2020; Wagner et al. 2020).

Currently, two main reduced representation approaches of OMICS-data are commonly used: restriction site-associated DNA sequencing (RAD-Seq; and similar methods) and target enrichment (hybrid capture). RAD-Seq covers a subset of anonymous, non-coding and coding regions across of the entire genome and is mostly used for population genomics and phylogenomics of closely related species within genera, up to tens of million years of divergence (Davey et al. 2011, Ree and Hipp 2015, McKain et al. 2018). RADseq is less costly and work-intense compared to target enrichment (McKain et al. 2018), and hence allows to process larger sample sets. Target enrichment usually addresses a subset of several hundreds of low-copy nuclear genes, and thus provides more conservative markers for resolving relationships within and among genera (Schmickl et al. 2016; McKain et al. 2018; Carter et al. 2019; Tomasello et al. 2020; Melichárková et al. 2020).

Although RAD-Seq yields much more information (number of loci and SNPs) than target enrichment, locus dropout caused by mutation accumulation in cutting sites can become more and more problematic with increasing species divergence (but see in Eaton et al. 2017 for the influence of sequencing coverage). Moreover, the correct definition of loci and filtering of paralogs based on anonymous short sequence reads is a bioinformatic challenge (Ree and Hipp 2015; O’Leary et al. 2018; McKain et al. 2018). Target enrichment loci are predefined from probe design and assembled loci are usually longer (McKain et al. 2018), allowing for gene tree estimation and allele phasing, and thus coalescent-based methods. Allelic information (segregating markers at a single locus) is particularly important for correct phylogenetic inferences in highly reticulate, young evolutionary relationships (Eriksson et al. 2018). Coalescent-based models can reconstruct correct species trees and estimate species boundaries while accounting for stochastic processes like ILS (Rannala and Yang 2003; Jones et al. 2015; Rannala 2015). In addition, plastid data can be easily gained from off-target reads of target enrichment (Hyb-Seq; Weitemier et al. 2014; Folk et al. 2015; McKain et al. 2018). The incorporation of plastid data into network reconstruction to gain information on the maternal progenitor has been largely overlooked in the last few years. Nuclear-plastid discordances have been assessed on shallow to deep phylogenetic scales, and elucidated group-specific evolutionary processes (Huang et al. 2014; Stull et al. 2020).

Elucidating the evolution of allopolyploids is even more challenging due to reticulate evolution. Tree methods can give a first phylogenetic framework for polyploid reconstructions when no previous phylogenetic study exists. Particularly in evolutionary young species complexes containing genetically close taxa (see e.g. McDade 1992 for tree stability in presence of hybrids from closely related taxa) and without any previous knowledge about auto- vs. allopolyploid origins, trees and (quartet) support values can give valuable information on conflicting signals. For example, a non-conflicted, tree-like pattern rather hints at autopolyploids whereas trees with low (quartet) support values can indicate the presence of reticulations and/or ILS (Lo et al. 2010; Brandrud et al. 2020; Karbstein et al. 2020b, Tomasello et al. 2020). However, hybridogenic, network-like origins cannot be inferred by both standard and coalescent methods based on bifurcating models, leading to incongruences in tree reconstructions (McBreen and Lockhart 2006; Rothfels 2021). Consequently, phylogenetic relationships should not (only) be presented by bifurcating trees (McDade 1992, 1995; Huson and Bryant, 2006; Rothfels 2021).

Distance-based network methods like for example the popular NeighborNet algorithm can visualize reticulate relationships better than trees, but detailed information on ancestry or parentage in hybrid scenarios requires phylogenetic networks (Huson and Bryant 2006; Oxelman et al. 2017). Recently developed software can model network-like evolution with maximum pseudolikelihood from gene trees or SNP-based multilocus sequence data under the coalescent model accommodating ILS (e.g., PhyloNet or PhyloNetworks; Than et al. 2008; Solís-Lemus et al. 2017; Wen et al. 2018; Olave and Meyer 2020; Flouri et al. 2020).

Phylogenetic network inference requires information on ploidy level and diploid progenitors, allowing correct heterozygosity estimations and allele phasing in polyploids. Recently, an increasing number of studies focused on network estimations and technical improvements in polyploid reconstructions (e.g., phasing, allele sorting, subgenome assignment, or modeling the (allo)polyloidization process; Bertrand et al. 2015; Jones 2017a; Oberprieler et al. 2017; Dauphin et al. 2018; Cao et al. 2019; Lautenschlager et al. 2020; Freyman et al. 2020, Yan et al. 2020; Šlenker et al. 2021; Tiley et al. 2021). Nevertheless, knowledge on putative progenitor species, number of contributing progenitors, ploidy levels, and formation types of the polyploids within large species complex are frequently missing. Moreover, only one resource-intensive program is currently capable to model the polyploidization process itself, i.e., that homeologues of an allotetraploid share demographic parameters or divergence times from their progenitors (Jones 2017a). In addition, for current allele assignment methods (e.g., Lautenschlager et al. 2020; Šlenker et al. 2021), subgenomes should be well genetically differentiated, sequences of the diploid parents available/not extinct, and locus/gene datasets not too big. Therefore, more sophisticated methods (e.g., polyploid networks, multi-labeled subgenome trees) are often not applicable at that stage of research and/or currently still inappropriate for young polyploid species complexes.

Even with this information, reconstruction of relationships and discrimination between auto- and allopolyploid scenarios might be difficult. For example, when using maximum likelihood (or pseudo-likelihood) approaches, likelihood scores of networks are usually not directly comparable to those of trees. A-posteriori model comparisons need to be applied to discriminate among scenarios with different numbers of reticulations (e.g., Kamneva et al. 2017; Cai and Ané 2020). Moreover, a correct and unequivocal network inference is hard to reconstruct in young species complexes where progenitors exhibit high levels of genetic admixture and polyploids possess high levels of genome-wide heterozygosity. Therefore, relationships, reticulate evolutionary processes, genome composition, structure, and evolution within large polyploid species complexes remain largely uninvestigated.

In this study, we unravel for the first time the evolutionary processes shaping apomictic polyploid complexes on the model system *Ranunculus auricomus* by using reduced-representation genomic data. The complex ranges from Greenland, Europe to Western Siberia, and spans arctic, boreal, temperate, and Mediterranean climates (Jalas and Suominen 1989). Linnaeus (1753) already described a species with dissected basal leaves, *R. auricomus*, and one with undivided basal leaves, *R. cassubicus*. Since then, more than 840 taxa (morphospecies) have been described, inhabiting stream- and riversides, and semi-dry to marshy meadows and forests (Karbstein et al. 2020b, 2021). Most of these taxa are tetra- to hexaploid and apomictic (Jalas and Suominen 1989; Karbstein et al. 2021). Only four di- and one tetraploid, genetically and geographically distinct, sexual species were detected so far and originated 0.83–0.58 Mya (Karbstein et al. 2020a, 2020b; Tomasello et al. 2020): *R. cassubicifolius* s.l. (di- and autotetraploid) and *R. notabilis* s.l. (diploid) are most distantly related whereas *R. flabellifolius*, *R. envalirensis* s.l. (both diploid), and *R. marsicus* (tetraploid), are grouped in intermediate positions (Karbstein et al. 2020b). *R. cassubicifolius* s.l. and *R. flabellifolius* are characterized by non-dissected basal leaves whereas the other species show a strongly heterophyllous cycle with dissected basal leaves during anthesis (Karbstein et al. 2020b).

Vicariance processes probably triggered allopatric speciation during climatic deteriorations in the late Pleistocene from a widespread European ancestor (Tomasello et al. 2020). It has been hypothesized that the large number of asexual, mainly tetra- to hexaploid polyploids arose from hybridization of sexual progenitors (Hörandl et al. 2009; Hodač et al. 2014, 2018; Hojsgaard et al. 2014; Barke et al. 2018). Some polyploid apomicts are probably less than 0.1 Mya (Paun et al. 2006; Pellino et al. 2013). They occupy larger, more northern areas, possess higher levels of genome-wide heterozygosity, and are obligate apomictic or with low levels of facultative sexuality (Karbstein et al. 2021). Nevertheless, origin, relationships, and genomic composition of the polyploid complex have never been analyzed due to genetic and bioinformatic limitations.

In this study, we compare a comprehensive taxon sampling based on genomic RAD-Seq data (280 individuals, 80 taxa), (phased) nuclear genes (113 individuals, 50 taxa), and plastid regions (87 individuals, 45 taxa), to unravel phylogenetic relationships and genome composition of the large, evolutionary young, *R. auricomus* polyploid complex. We use a comprehensive, self-developed bioinformatic pipeline combining previous knowledge about sexual progenitors, ploidy and reproductive data with tree, structure, network, and SNP-origin methods across different datasets (Fig. 1) to answer the following questions: (i) Are the applied tree analyses able to give a first phylogenetic framework, and do well-supported (main) clades exist? (ii) Do genomic, nuclear-gene, and plastome data reveal congruent tree topologies or rather conflicting signals due to reticulate evolution? (iii) Do RAD-Seq or phased nuclear gene data reflect any clear genetic and/or geographical structure? (iv) Are polyploid lineages of auto- or allopolyploid origin? (v) If the latter, how many progenitors contributed to their genomes? (vi) To which extent are polyploid genomes influenced by post-origin evolution? (vii) How can analyses of RAD-Seq and Hybseq data be integrated for unraveling evolutionary processes in polyploid complexes?

## Materials & Methods

### Population Sampling

In the present study, we included four di- and one tetraploid sexual species, and 75 of the most widespread, tri- to hexaploid apomictic *R. auricomus* taxa. The new classification of sexual species is described in Karbstein et al. (2020b). Ploidy and reproduction mode measurements of *R. auricomus* individuals and populations (sexual, and facultative and obligate apomictic) needed for the performed analyses herein are published in Karbstein et al. (2021) (see also Supplementary Table S1, Figs. S4, S5 in Karbstein et al. 2021, and data on FigShare). The diploids *R. sceleratus* and *R. pygmaeus* were used as outgroups. We collected silica-gel dried leaf material from living plants for all genetic analyses, and additionally, leaf material from herbarium specimens for target enrichment analyses. Finally, we used 280 samples originating from 235 collection sites (populations) across Europe for further analyses (Fig. 2, Supplementary Table S1). Concerning genomic analyses, the sexual progenitor species were treated as parental subgenomes and abbreviated according to the legend of Fig. 2.

**Fig. 2.**
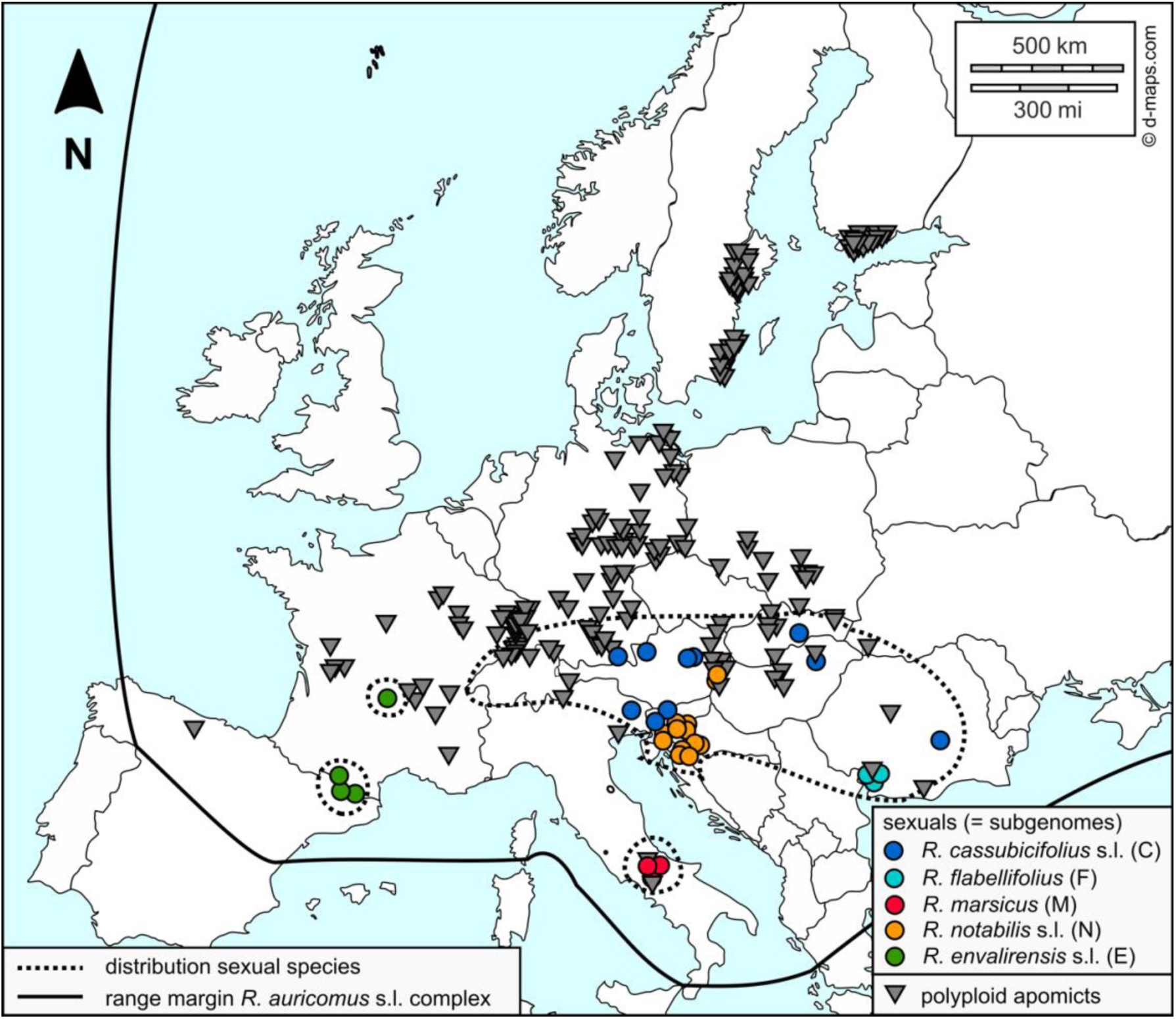
Locations of studied *R. auricomus* populations across Europe. We investigated 235 sexual and apomictic populations (see Supplementary Table S1 for details). Symbols represent reproduction modes of populations (colored circles = sexuals, defined as subgenomes here for further data analyses), dark grey triangles = obligate or facultative apomictic, also in Karbstein et al. (2020b, 2021). Circles of sexual species were highlighted according to the color scheme of Fig. 4. The solid line shows the range margin of the *R. auricomus* complex, and the pointed lines highlight the distribution of sexual species. The original map was downloaded from https://d-maps.com/, created by Karbstein et al. (2021), and modified for this study.

### Laboratory Work, Locus Assembly, and Parameter Optimization

DNA extraction of 280 *R. auricomus* and outgroup samples, adjustment of DNA concentration, DNA quality check, RAD lab workflow and sequencing with the cutting enzyme *PSTI* and single-end RAD sequencing of 100bp reads (Baird et al. 2008), raw read quality check, raw read demultiplexing, removal of adapter sequences and restriction overhang, and further quality filtering in IPYRAD (Eaton and Overcast 2020) followed Karbstein et al. (2020b, 2021). Here, we used the already sequenced samples of Karbstein et al. (2020b, 2021).

*PSTI* is a methylation-sensitive enzyme and hence can considerably reduce the fraction of repetitive elements that is otherwise very high in plants. Therefore, the enzyme targets mostly nuclear genes and a few organelle sites (Fellers 2008). This dataset of coding- and non-coding regions complements the markers derived from expressed genes (transcriptomes of flowering buds) and selected for target enrichment (see Tomasello et al. 2020 for baits design in *R. auricomus*) to get a comprehensive representation of the nuclear genome. Target enrichment further provides markers of the plastid genome. The self-developed, bioinformatic pipeline combining different datasets and analyses, and previously published *R. auricomus* studies (sexual progenitor species, reproduction modes) is illustrated in Fig. 3.

**Fig. 3.**
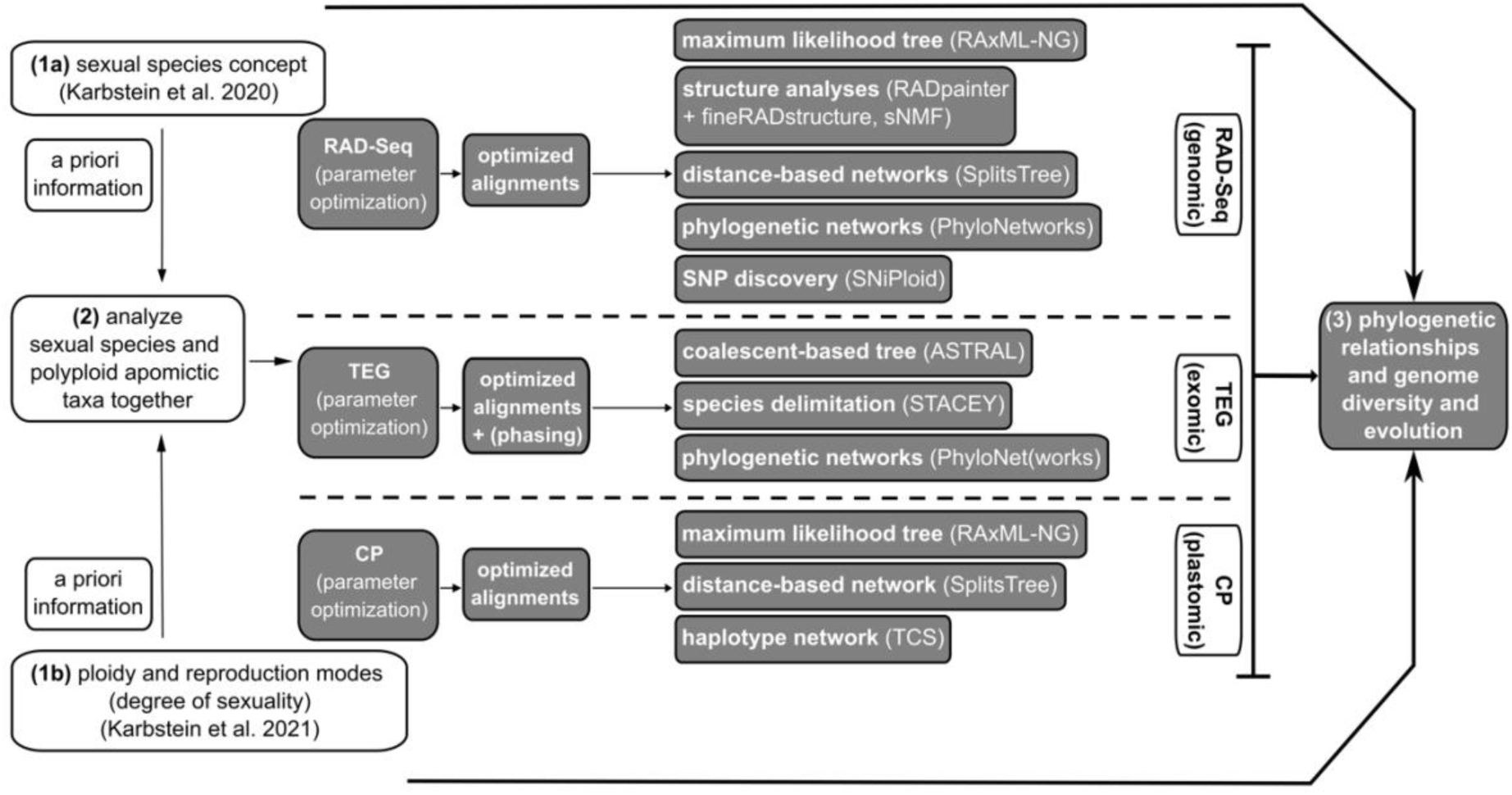
Bioinformatic pipeline to resolve polyploid species complexes. Here, we used *R. auricomus* as a model system and basically followed the concept of Hörandl (2018) to disentangle complicated species complexes. We analyzed (2) sexual species and apomictic taxa together, using a priori information about (1a) sexual species (Karbstein et al. 2020b) and (1b) ploidy levels and reproduction modes (Karbstein et al. 2020b, 2021). Analyses are based on the optimized alignments of three different datasets covering genomic parts (RAD-Seq), exomic nuclear regions (target enrichment, TEG), and plastome regions (chloroplast, CP). We used RAD-Seq datasets to calculate maximum likelihood (ML) trees, genetic structure analyses, distance-based networks, maximum pseudolikelihood networks, and SNP discovery analyses. To study the robustness of RAD-Seq results, we computed coalescent-based trees, species delimitation analysis, and maximum pseudolikelihood networks based on target enrichment datasets. A ML tree, and distance-based and haplotype networks of the CP dataset were also included to get further details about hybridogenic origins of polyploids.

For target enrichment, we added 85 newly sequenced polyploid apomictic samples to the already existing 28 samples sequenced by Tomasello et al. (2020) (113 samples in total; Supplementary Table S1). All plastome data (CP) from off-target reads is published here. We included almost the same samples as in the RAD-Seq analyses and added as described above herbarium specimens (types or collections from type locations; Supplementary Table S1). We used the bait set as described previously in Tomasello et al. (2020), consisting of 17,988 probes and capturing 736 target genomic regions. Library preparation and hybrid capture protocols are available as Supplementary Text S1 in Tomasello et al. (2020). Libraries were sequenced in five different paired-end runs (24 samples each) with 2×250-bp on an Illumina MiSeq system at NGS Integrative Genomics Core Unit of the University of Göttingen (Germany).

For *de novo* assembly of RAD-Seq loci and parameter optimization, we used IPYRAD vers. 0.9.14 (and vers. 0.9.52) on the local HPC-Cluster (GWDG, Göttingen, Germany). For parameter optimization, we applied an already established workflow accounting for different ploidy levels of *R. auricomus* individuals: The within-sample clustering similarity threshold was optimized for each ploidy level (*2n*–*6n*) balancing number of RAD-Seq clusters, cluster depth, and clusters rejected due to high heterozygosity. Then, the among-sample clustering threshold was optimized for the merged assembly optimizing number of polymorphic loci, SNPs, loci filtered by maximum number of SNPs, removed duplicates, shared loci, and new polymorphic loci. Maximum number of SNPs per locus and of indels per locus were increased to 30% and 12, respectively, to account for greater genetic variation in polyploids as described in Karbstein et al. (2021).

For subsequent analyses, we created a ‘without-outgroup’ and a ‘total’ dataset. To assess effects of number of loci and missing data on phylogenetic analyses (Eaton et al. 2017; O’Leary et al. 2019; Karbstein et al. 2020b), we selected different minimum amounts of samples per locus and created ‘min10’ (10%), ‘min30’ (30%), and ‘min50’ (50%) alignments balancing the specific program requirements and informativeness of datasets (see below). The final sample size totals 282 individuals (incl. outgroup). For both datasets, sample filtering led to ca. 74% (min10), 55% (min30), and 44% (min50) missing data in the final sequence matrices.

For target enrichment data analysis, reads were processed with HybPhyloMaker vers. 1.6.4 (Fér and Schmickl 2018) (Supplementary Text S1), using the target regions (exons) selected for the bait design from transcriptomes as ‘pseudoreference’ for read mapping (Supplementary Table S2 in Tomasello et al. 2020). Samples with more than 40% missing data were filtered out from each exon region. In addition, only loci including more than 90% of samples were further processed (579 genes). From those 579 genes, 50 loci were selected for the species delimitation and phylogenetic network analyses, to be informative, non-homoplasious, and free from paralogue sequences. To select these loci, we assessed four different parameters across the 579 alignments, scoring the respectively best performing 25% of loci with 1 and the remainder with 0. The parameters were the following (Herrando-Moraira et al. 2018): (i) the R^2^ of mutational saturation regression curves (Philippe and Forterre 1999), (ii) the standard deviation of the sample-specific long-branch scores (LB scores; Struck et al. 2014), (iii) the clocklikeness, and (iv) average bootstrap (BT) support. This was done (i-iii) following the idea that in such a young species group like *R. auricomus* evolutionary rates will not change considerably among branches in orthologous regions. Finally, we selected the 50 loci with the highest overall score (Supplementary Table S3).

For inference of polyploid origins, retrieving allelic information is crucial. We phased these 50 most informative loci using a similar approach as described in Eriksson et al. (2018). We processed the mapped BAM files of all samples with SAMtools vers. 0.1.19 (Li et al. 2009) (‘sort’ and ‘phase’ commands). The polyploid samples (tri- to hexaploids) were phased further, looking at the phased BAM files in IGV vers. 2.8.9 (Robinson et al. 2011) (usually one of the BAM files was a consensus of alleles with some vowels corresponding to the heterozygous sites) and manually adding alleles in relation to the known ploidy level to the alignments using AliView vers. 1.26 (Larsson, 2014). For two of the 50 loci, it was not possible to unequivocally detect alleles in at least one of the polyploid samples. Therefore, we excluded these two loci from further analyses (Supplementary Table S4).

Off-target reads were used to gain information on the plastid genome again performing HybPhyloMaker. We used the *Ranunculus repens* plastid genome as reference (Supplementary Table S5). Considering the low number of mapped reads and the resulting highly fragmented alignments, we excluded regions and samples with the highest amounts of missing data, to minimize phylogenetic inaccuracy in the subsequent analyses. First, we excluded samples with more than 50% missing data (from each plastome region separately), and regions containing sequence information for fewer than 50% of the samples. Second, we excluded from all regions all samples missing from more than 50% of the alignments. After filtering, we retrieved a subset of 71 regions including genes and intergenic spacers, for 87 samples from the original 113 (Supplementary Table S6).

### Maximum Likelihood Tree and Quartet Sampling (RAD-Seq)

To infer phylogenetic relationships among sexuals and polyploid apomicts (Figs. 1, 3), maximum likelihood (ML) analyses were performed with RAxML-NG vers. 0.9.0 (Kozlov et al. 2019) on the final RAD-Seq min10, min30, and min50 datasets. As input, we used the *.phy IPYRAD output files (each individual characterized by one sequence [majority-rule base calling], all loci concatenated into a supermatrix). Alignment patterns were compressed and stored in binary formats (RBA). Then, we inferred the respective tree under the GTR+GAMMA model with 10 random and 10 parsimony starting trees. Standard non-parametric BT were performed, and the MRE-based bootstopping test (cutoff: 0.05) applied. Felsenstein bootstrap proportion (FBP) and transfer bootstrap expectation (TBE) values were calculated by RAxML-NG. TBE is more appropriate for large phylogenies (>300 samples) and for phylogenies with conflicted (deep) branches compared to FBP (e.g., hybridization events; Lemoine et al. 2018). FBP and TBE values were mapped by RAxML-NG onto the best-scoring ML trees. The min10 compared with min30 and min50 alignments showed the highest mean BT support (FBP: 70/66/47, TBE: 85/82/64) and the largest number of monophyletic taxa (13/10/9) (Supplementary Figs. S1–S3a–d). Moreover, the min10 tree topology was similar to min30 and min50 tree topologies (Fig. 4a, Supplementary Figs. S1, S2). Therefore, we selected the min10 tree for final interpretations.

**Fig. 4.**
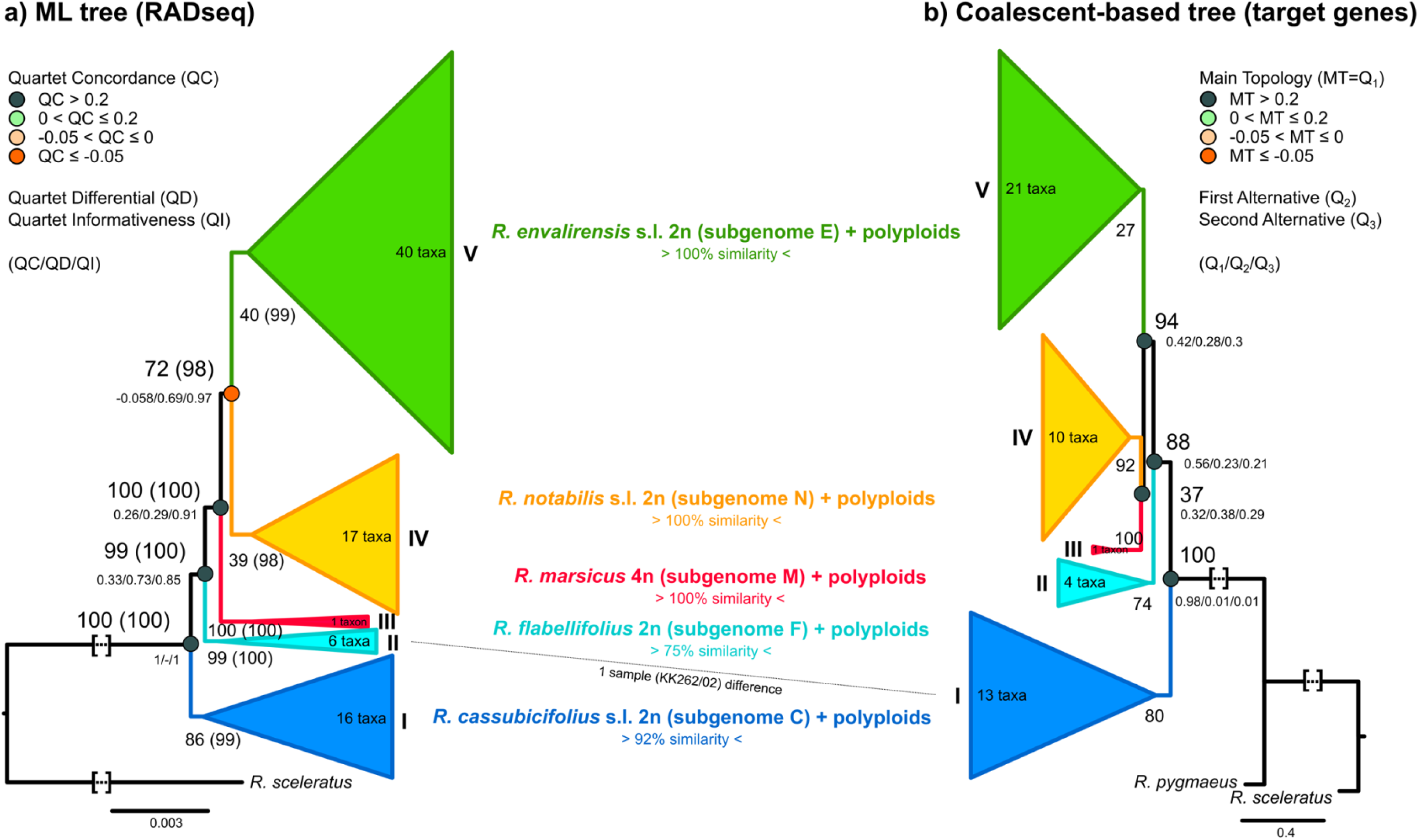
Phylogenetic trees based on RAD-Seq and target enrichment data. (a) a ML tree based on RAxML-NG results and a min10 RAD-Seq alignment (280 samples, 97,312 loci, 438,775 SNPs) and (b) a coalescent-based tree based on ASTRAL results and a target enrichment alignment (113 samples, 576 non-phased genes). For Fig. 4a, FBP, TBE, and quartet sampling scores (QC/QD/QI; see Supplementary Fig. S3a-d and legend for explanations) are displayed per branch. Nodes are colored according to QC values (legend on the left). For Fig. 4b, FBP and quartet support scores (MT=Q_1_/Q_2_/Q_3_) values are shown per branch. In general, only supported main clades (I-V, FBP/TBE>70; except Fig. 4b split between *R. notabilis* s.l. and *R. marsicus*) are illustrated because of mostly low/no BT support (BT<70) within clades. Each clade contains a sexual species and several polyploids. Number of polyploid taxa is given per main clade. We calculated sample composition between RAD-Seq and target enrichment clades (only shared samples were evaluated due to different sample sizes), and illustrated values in the central part of the figure (0%=main clades are composed of completely different samples, 100%=main clades are composed of completely equal samples). Dotted lines show differences between clades of both datasets. See Supplementary Figs. S1, S2, S4a-c and Figshare data repository for more details. Squared brackets: A part of the branch was cut for illustrative purposes.

We addressed potentially inflated BT values in concatenated analyses with ‘Quartet Sampling’ (QS) vers. 1.3.1 (Weisrock et al. 2012; Shen et al. 2017; Pease et al. 2018; and see also Karbstein et al. 2020b). The quartet concordance score (QC) is defined as the ratio of concordant to both discordant quartets (1: all concordant, > 0: more concordant patterns, < 0: more discordant patterns), the quartet differential score (QD) indicates the skewness of both discordant patterns (1: equal, 0.3: skewed, 0: all topologies 1 or 2), and the quartet informativeness score (QI) describes the proportion of informative replicates (1: all informative, 0: none informative; see Pease et al. 2018). QD values around 1 indicate ILS (presence of both discordant topologies) whereas QD values towards 0 hint at directional introgression (presence of one alternative topology; Pease et al. 2018; see also Karbstein et al. 2020b). We set 100 replicates per branch and log-likelihood threshold cutoff to 2. The quartet concordance factor (QC), quartet differential (QD), and the quartet informativeness (QI) scores together with BT values were illustrated in Fig. 4a (detailed QS values in Fig. S3a–d).

### Coalescent-based Species Tree and Quartet Support (Targeted Genes)

As an equivalent to ML trees for RAD-Seq alignments, we estimated a coalescent-based tree based on 576 target enriched genes. First, gene trees were inferred in RAxML vers. 8.2.12 (Stamatakis 2014). Analyses were run with 100 standard BT replicates, setting the GTR+GAMMA model and partitioning by exons. Second, gene trees were rooted and combined into a single newick file in HybPhyloMaker (Fér and Schmickl 2018). Third, the species tree was inferred by applying the coalescent-based algorithm implemented in ASTRAL III vers. 5.6.3 (Zhang et al. 2018) with 100 multilocus BT replicates. To assess the amount of gene tree conflict on branches, we measured quartet support on the ASTRAL tree (Sayyari and Mirarab 2016; Fig. 4b, Supplementary Fig. S4a–c).

### Plastome Phylogeny and Network Analysis (CP)

We used the 71 selected plastid regions to infer a ML tree with 100 BT replicates by RAxML-NG (Kozlov et al. 2019). Models of sequence evolution were assessed for each region separately using ModelTest-NG vers. 0.1.6 (Darriba et al. 2020). Alignments were concatenated (80,461 base-pairs in total) and different regions were treated as different partitions, each with its respective sequence evolution model.

To gain additional information about haplotype evolution, the concatenated matrix was used to infer a haplotype network with TCS vers. 1.13 (Clement et al. 2000). We used the web-based software tcsBU (Múrias Dos Santos et al. 2016) to produce a graphical representation of TCS haplotype network. In addition to the TCS networks, we calculated neighbor-net networks as described in Karbstein et al. (2021).

### Genetic Structure (RAD-Seq)

To investigate genetic structure of the polyploid species complex (Figs. 1,3), we first conducted analyses with RADpainter+fineRADstructure vers. 0.3.2 (Malinsky et al. 2018) using the *.alleles.loci IPYRAD output files (each locus with a maximum of four allowed alleles (only two phases due to diploid SNP calls), with individual sequences). RADpainter is based on a coancestry matrix, uses all SNPs, allows a varying allele number, and tolerates moderate amounts of missing data (Malinsky et al. 2018). We ran RADpainter to calculate the coancestry matrix, and used fineRADstructure to assign individuals to groups (1,000,000 burn-in and 1,000,000 sample iterations) including a simple tree building (MCMC; 100,000 burn-in). Finally, we plotted results using a modified R script of ‘fineRADstructurePlot.R’ (R vers. 4.0.3 (R Core Team 2020) for all R analyses). We then compared results of min10, min30, and min50 alignments. With increasing number of loci and missing data, an increased number of groups and genetic dissimilarity among groups was detected (Supplementary Fig. S5). Thus, we selected min10 for further interpretations.

Second, we carried out structure analyses applying sNMF within the R package ‘LEA’ vers. 3.0.0 (Frichot et al. 2014; Frichot and François 2015). sNMF provides a fast and efficient estimation of individual coancestry, is robust to deviations from Hardy-Weinberg equilibrium, and can deal with moderate levels of missing data (Frichot and François 2015). We used *.ugeno (each individual characterized by numbers indicating one randomly chosen per SNP locus) IPYRAD files, and set number of genetic clusters (K) from 1 to 80 (maximum number of included taxa), ploidy to 4 (as maximum), and repetitions to 7. To choose the number of ancestral Ks, we used the implemented cross-entropy criterion. Cross-entropies were plotted for all Ks. Across datasets, we found the optimal Ks between 3 and 5 (Supplementary Fig. S6a–f). We plotted results of optimal Ks as bar graphs and across Europe (Supplementary Figs. S7a–c, S8a–c). Additionally, we displayed ancestry coefficients (method ‘max’, i.e., at each point the cluster for which the ancestry coefficient is maximal; Supplementary Figs. S9–11 without method ‘max’) of the respective best run of each K on geographical maps of Europe, using location coordinates and the R script POPSutilities.R (source: http://membres-timc.imag.fr/Olivier.Francois/POPSutilities.R; Supplementary Fig. S8a–c). The min30 datasets balanced number of loci and amounts of missing data and revealed the most reasonable results (see explanation in legend of Supplementary Fig. S8, and was therefore selected for further interpretations.

### Genetic Structure (Targeted Genes)

To unravel the genetic structure of the polyploid complex based on nuclear genes (Figs. 1, 3), we utilized the coalescent-based species delimitation approach of STACEY vers. 1.2.1 (Jones 2017b). Input files were prepared in BEAUTI vers. 2.6.1 (Bouckaert et al. 2014) using the 48 phased loci. For the analyses, each sample was treated as ‘minimal cluster’ (i.e., alleles of the same individuals were represented by a single tip in the species tree/species delimitation results). Sequence substitution models were selected for each locus separately using the Bayesian Information Criterion (BIC) in ModelTest-NG. Substitution models, clock models, and gene trees were treated as unlinked for all loci. To reduce the search space, parameters of the substitution models were fixed to those found in ModelTest-NG. The strict clock was enforced for all loci fixed at an average rate of 1.0 in one random locus while estimating all other clock rates in relation to this locus. We set the ‘collapse height’ to 1×10^-5^, which was estimated using a Beta prior with parameters α=1.0 and β=1.0, and which represent a flat distribution between 0 and 1 (i.e., all possible species delimitation scenarios have an equal prior probability). Finally, we gave to the bdcGrowthRate prior a log-normal distribution (M=4.6 and S=1.5), a gamma shape (α=0.1 and β=3.0) to the popPriorScale prior, and for the relativeDeathRate, we set a beta prior (α=1.0 and β=1.0; optimized in Karbstein et al. 2020b).

The analyses were run for 2×10^9^ iterations sampling every 200,000th generation in BEAST vers. 2.6.1 (Bouckaert et al. 2014). Two independent runs were performed and, after checking convergence between independent analyses and Effective Samples Size values (ESS > 100) in TRACER vers. 1.6 (Rambaut et al. 2018), we combined trees output files using LogCombiner vers. 2.6.1 (Bouckaert et al. 2014) and discarding 10% of the analyses as burn-in (as described in Karbstein et al. 2020b). The obtained file was processed with the ‘species delimitation analyser’ (Jones et al. 2015). The similarity matrix was produced using a modified version of the R script (Jones et al. 2015).

### Detecting Subgenome Contribution for selected Polyploids (RAD-Seq, Target Genes)

To investigate genome diversity, composition, and evolution of polyploids in more in detail (Figs. 1,3), we selected 2–4 polyploid individuals with obvious reticulation (coancestry) signals per main genetic cluster and tested ten polyploids for subgenome contributions (H_1_-H_10_; Table 1, Supplementary Table S1). We used hybrid binomials to clearly distinguish these taxa from sexual species (Hörandl et al. 2009). RADpainter+fineRADstructure analyses include all SNPs per locus and varying ploidy levels (Malinsky et al. 2018), and is therefore here considered as superior compared with sNMF and SplitsTree (1 SNP/locus). Therefore, we evaluated the RADpainter coancestry matrix, and calculated a median coancestry value of 343 (mean right-skewed). We took the median as the critical threshold to assess the potential subgenome contributions of polyploids. The same procedure was also applied to the STACEY posterior probability matrix (median=0.000555). To ensure comparability among datasets, we aimed at selecting the same individuals. Only for ‘*R.* × *elatior*’ (H_2_), we selected another individual in STACEY analysis (*R.* × *elatior* is monophyletic, Supplementary Figs. S1–S4a–c).

**Table 1.**
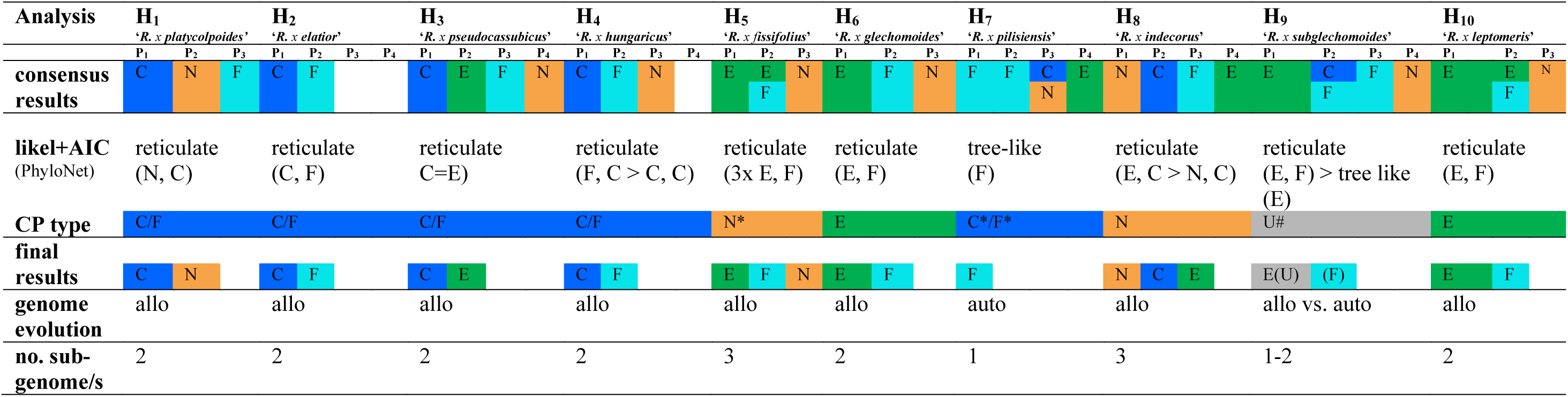
Genetic Structure and Phylogenetic Network results of tested tetraploid *R. auricomus* accessions (H_1_-H_10_). Each row (H_1_-H_10_) represents a separately analyzed individual. Results are based on RAD-Seq (RADpainter+fineRADstructure, PhyloNetworks) and phased nuclear target enrichment gene (STACEY, PhyloNetworks, Phylonet) datasets. Consensus results summarize all previously gained information. Final results indicate final subgenome contribution(s), i.e., consensus results corrected by the full likelihood approach+AIC calculations in Phylonet (likel+AIC, AIC = Akaike Information Criterion; more than one result if AIC network difference was less than 10 units) and plastome analysis results (CP type; C/F= plastid type shared by the diploid sexual species *R. cassubicifolius* and *R. flabellifolius*, *=not the same sample between CP and network analyses, #=haplotype from an unknown/extinct sexual progenitor species of Central Europe). Concerning the final results of H_9_, we classified P_1_ as “E(U)” because of the *R. envalirensis*-like U plastid type (see above). According to final results, we classified genome evolution of investigated polyploids (allo=allopolyploid, auto=autopolyploid), and the number of involved subgenomes in polyploid formation. See also Figs. 5, 6, 8, Supplementary Tables S1, S7, and data on Figshare for sample IDs, genetic structure, and network results.

### Phylogenetic Network Analyses (RAD-Seq)

To corroborate the already gained information by appropriate network methods (Figs. 1,3), we carried out analyses with PhyloNetworks vers. 0.12.0 (Solís-Lemus et al. 2017). PhyloNetworks allows network inference with maximum pseudolikelihood from multilocus sequences (SNaQ). SNaQ uses a multi-species network coalescent model (MSNC) that is capable of handling ILS (Solís-Lemus et al. 2017). Since SNPs were not intended as input for SNaQ, we used the recently published function SNPs2CF.R vers. 1.2 (Olave and Meyer 2020) to transform SNP-based RAD-Seq alignments into quartet concordance factors (CF). We selected the min30 dataset to avoid bias of network analyses by excessively high amounts of missing data. We converted the *.ustr (each individual characterized by numbers indicating one randomly chosen SNP per locus, two phases (maximum of four allowed alleles) per individual) IPYRAD file with a custom R script to an adequate input format for SNPs2CF. Network analyses are computationally intensive. Thus, we created ten subsets each containing one tetraploid accession (individual) and all available accessions (individuals) of diploid sexual progenitor species. The above-mentioned, preselected tetraploids were used for subset building (see Detecting Subgenome Contribution for selected Polyploids). We excluded the sexual tetraploid *R. marsicus* from network analyses because no significant subgenome contribution of this species was observed in previous genetic structure analyses. We used the converted *.ustr files and imap files containing individual-species associations as input for SNPs2CF. We specified ‘between species only’ comparisons, no maximum number of SNPs, maximum number of quartets of 1000, and 100 BTs.

We used the received quartet CF matrices and quartet-CF-based starting trees to run maximum pseudolikelihood (SNaQ) analyses with default settings. We initially allowed no hybridization event. Afterward, the output was used as a start network (net0) for the next analysis allowing one hybridization event (net1). Per polyploid, the likeliest network was commonly the one with the polyploid as hybrid (seven out of ten). The polyploids H_4_, H_5_, and H_9_, were not inferred as the likeliest hybrids. An explanation might be the low genetic divergence among polyploids and diploid progenitors causing problems in ILS and hybridization modeling. However, since SNaQ (PhyloNetworks) takes no hybrid constraint and polyploids cannot be the progenitor of diploid sexuals here, we had to select the less likely hybrid network in these cases for further polyploid analyses.

### Phylogenetic Network Analyses (Targeted Genes)

To assess the validity of previous structure and RAD-Seq-network results (Figs. 1, 3), we additionally performed phylogenetic network analyses using the 48 phased target genes. We also investigated H_1–10_, taking gene trees as input and two different, separately performed, coalescent-based approaches: SNaQ implemented in PhyloNetworks (Solís-Lemus et al. 2017) as for RAD-Seq data and the maximum pseudolikelihood (InferNetwork_MPL) approach implemented in Phylonet vers. 3.8.2 (Than et al. 2008; Wen et al. 2018). Phylonet is one of the most widely used and established programs for species tree/network reconstructions based on multilocus datasets. We told both programs that all alleles (phases in RAD-Seq-based networks) of diploid species are from their respective species, and alleles (pseudophases in RAD-Seq-based networks) of the polyploid are only from the single polyploid accession. Thus, we used network results based on phased nuclear genes for further validation of previous results.

For each polyploid testing, alignments were modified to include all diploid accessions (except for *R. cassubicifolius* s.l. LH006 and EH9126, and *R. flabellifolius* LH021; 22 samples in total) and the polyploid individuals. Models of sequence evolution were selected with ModelTest-NG, and 100 BT gene trees were inferred with RAxML-NG for each of the 48 selected loci. Therefore, 100 gene trees per locus (4,800 trees in total) were used as input, to incorporate gene tree uncertainty while inferring species networks (and to ensure dataset comparability for SNAQ and PhyloNet). For the PhyloNetworks analyses, we used the gene trees and a mapping file (mapping alleles to species) to calculate a species-wise CF table. We continued the analyses as for the RAD-Seq dataset, with the only exception that the starting tree was inferred using ASTRAL III. For the Phylonet MPL analyses, the polyploid was always specified as the putative hybrid. We performed 10 runs per search, each returning five optimal networks. After the search, the returned species networks were optimized for their branch lengths and inheritance probabilities under full likelihood (-po option in PhyloNet), using the default settings.

### Subgenome Contributions and Polyploid Origin (RAD-Seq, Target Genes, and CP)

We applied criteria for building consensus results on previously generated genetic structure and phylogenetic network results (details in legend of Supplementary Table S7). We mainly assessed the parental subgenome contributions per polyploid individual as follows: (i) take the most abundant parent within a column; (ii) if there were two equally abundant parents (e.g., two-times sexual progenitor subgenomes C and F) within a column, both parental subgenome contributions were taken for the consensus result (e.g., C/F); (iii) if two parental subgenome contributions within a column existed, we included them with a value of ‘0.5’ (instead of ‘1.0’) in consensus calculations.

To validate the obtained consensus results and to infer genome evolution (tree-like, autopolyploid vs. network-like, allopolyploid), we submitted all previously generated results (before consensus results building) to the full likelihood approach implemented in PhyloNet. The CalcProb function calculates the likelihood of gene trees under a given species network and thus the total likelihood of the same network. Thus, we employed the gene trees used in the network analyses based on the target enrichment dataset mentioned above.

To include RADpainter+fineRADstructure and STACEY results, networks were manually constructed using the tree backbone topology in Karbstein et al. (2020b) and the first two putative progenitors identified by these methods (Supplementary Table S7). The autopolyploid scenario was tested utilizing the ASTRAL III trees already used as starting tree for the PhyloNetworks analyses. We rooted all networks with *R. cassubicifolius* s.l. to make scenarios more comparable. To compare tree-like (autopolyploid) scenarios with network-like (allopolyploid) ones, we scored results using the Akaike Information Criterion (AIC), taking into account that the number of parameters in a tree/network is equal to the number of branch lengths plus (for the networks) the parental contributions (i.e., k=8 and k=13 for the tree and the networks, respectively).

We determined the final subgenome contribution(s) by correcting the consensus results by the previously generated full likelihood approach results of Phylonet and plastome (CP) results (Supplementary Table S7). According to final results, we classified the origin of polyploids, and the number of subgenomes involved in polyploid formation.

### SNP Discovery (RAD-Seq)

To investigate post-origin evolution of allopolyploids in more detail (H_1_–H_6_, H_8_, and H_10_; Figs. 1, 3), we carried out SNiPloid (vers. 17^th^ March 2016; Peralta et al. 2013) analyses mainly following the workflow of Wagner et al. (2020) (see bash-script on Github). SNiPloid compares the genome of an allotetraploid and a diploid putative parental species (DIPLOID2) with a diploid parental reference (DIPLOID1). The resulting SNPs were categorized: cat1&2 result from post-origin interspecific hybridization, e.g., backcrossing to the parental species; cat3&4 represent post-origin lineage-specific SNPs (not present in the parents); cat5 represents the homeo-SNPs from the hybrid origin from the two parents (Peralta et al. 2013; Wagner et al. 2020). For example, a first-generation-hybrid is expected to have only homeo-SNPs inherited from the parental species (cat 5), and no interspecific SNPs (cat 1,2) or derived SNPs (cat 3/4).

We created references of diploids by merging all accessions of a single progenitor species into a single *.fastq file (all possible parental SNPs of genetically close individuals of a species have to be covered) and conducted within-sample clustering in IPYRAD (filtering and clustering settings identical to Karbstein et al. 2020b). Obtained consensus files were used as DIPLOID1 (reference) and merged *.fastq files as DIPLOID2. We specified a minimum read depth per position of 20 (default; majority of positions showed more than 100 reads coverage). First, we excluded the category ‘others’ (heterozygous positions of DIPLOID2) from final results. High percentages of this category (30–64%) are probably due to multi-sample accessions and high individual heterozygosity in natural diploid populations. To address the influence of ‘others’, we evaluated this category by splitting heterozygous positions (REF and ALT), and categorizing the remaining ALT SNPs of DIPLOID2 and REF SNPs of DIPLOID1 according to SNP categories of SNiPloid. Moreover, we always observed a dominance of interspecific SNPs of cat2 compared with cat1 SNPs, independent of parental combination. This was probably due to neglection of natural genetic variation in the majority rule base call references. Therefore, we generally summarized both categories to ‘cat1&2’ to avoid biases within interspecific SNP categorization.

## Results

### Phylogenetic Tree Analyses unraveled Five Main Clades and showed Large Congruence among Datasets

For sexual and asexual *R. auricomus* individuals across Europe, we generated genomic RAD-Seq, nuclear target enrichment gene, and plastome (CP) data based on 97,312 loci (280 individuals), 576 genes (113 individuals), and 71 regions (87 individuals), respectively. Both ML (RAD-Seq) and coalescent-based (nuclear genes) phylogenetic tree analyses revealed five main clades (I–V). BT support of tree ‘backbones’ was generally high (most FBP/TBE values 90–100, Fig. 4a,b). Clades were well supported (FBP/TBE values 70–100), but particularly FBP support of clades IV and V for the ML (39–40 vs. 98–99 TBE support) and of clade V for the coalescent-based tree (27) was very low. Within clades, BT support was very low or absent (most FBP/TBE values 0–80/40–90; Supplementary Fig. S4a–c). Each clade contained one sexual species and polyploid taxa of various geographical origins (ML and ASTRAL taxon names, respectively): (I) *R. cassubicifolius* (subgenome C) with 13 and 16 tetra- and hexaploid samples, (II) *R. flabellifolius* (subgenome F) and with 4 and 6 tri- to hexaploid, (III) *R. marsicus* (subgenome M) and one tetra- to hexaploid, (IV) *R. notabilis* (subgenome N) and with 10 and 17 tetraploid, and (V) *R. envalirensis* (subgenome E) and with 21 and 40 tetraploid taxa. Whereas clades I and II were predominantly characterized by undivided basal leaf types, clades III–IV exhibited only dissected ones.

ML tree nodes (RAD-Seq) were highly informative (QI=0.85–1, Fig. 4a). Quartet concordance metrics (QC) for RAD-Seq and main topology for target genes (MT=Q_1_) showed highly concordant patterns with almost no alternative topologies (QC=1, QD=-/MT=0.98, Q_2_=0.01. Q_3_=0.01, Fig. 4a,b) for the node splitting clade I and all remaining ones. All remaining nodes showed moderately to highly conflicting signals with varying distribution of alternative topologies (QC=-0.06–0.33, QD=0.29–0.73/MT=0.32–0.56, Q_2_=0.23–0.38, Q_3_=0.21–0.30), particularly the nodes splitting main clades IV and V (QC=-0.06)/M+N and E (MT=0.42), supported by lowered BT values (72–98). The sample composition of clades was highly similar between the different approaches.

### Plastome Phylogeny showed Incongruences with Nuclear Data

The ML tree based on plastome data (CP) revealed four well-supported main clades with FBP=100 (i.e., haplotype groups; Fig. 5a, Supplementary Fig. S12a–d). In general, within-clade (within-haplotype-group) relationships were mainly low or not supported (FBP <<70). Clade I consists of haplotypes from *R. cassubicifolius*, *R. flabellifolius*, and various polyploids. Within the first clade, accessions of *R. cassubicifolius* and *R. flabellifolius* were completely intermingled, contrary to nuclear datasets (Fig. 4). Clade II contained only haplotypes from polyploid taxa. The remnant two haplotype clades III and IV consisted of *R. envalirensis* and few polyploid accessions, and *R. notabilis*, *R. marsicus*, and various polyploids, respectively. Interestingly, accessions of the diploid *R. notabilis* and the tetraploid *R. marsicus* are intermingled, indicating that they belong to the same haplotype group (Supplementary Fig. S12b), contrary to nuclear data (Fig. 4). The splitsgraph of the neighbor-net analysis also exhibited four, weakly differentiated, clusters (Fig. 5b, Supplementary Fig. S13). The same is true for the TCS haplotype network (Supplementary Fig. S14).

**Fig. 5.**
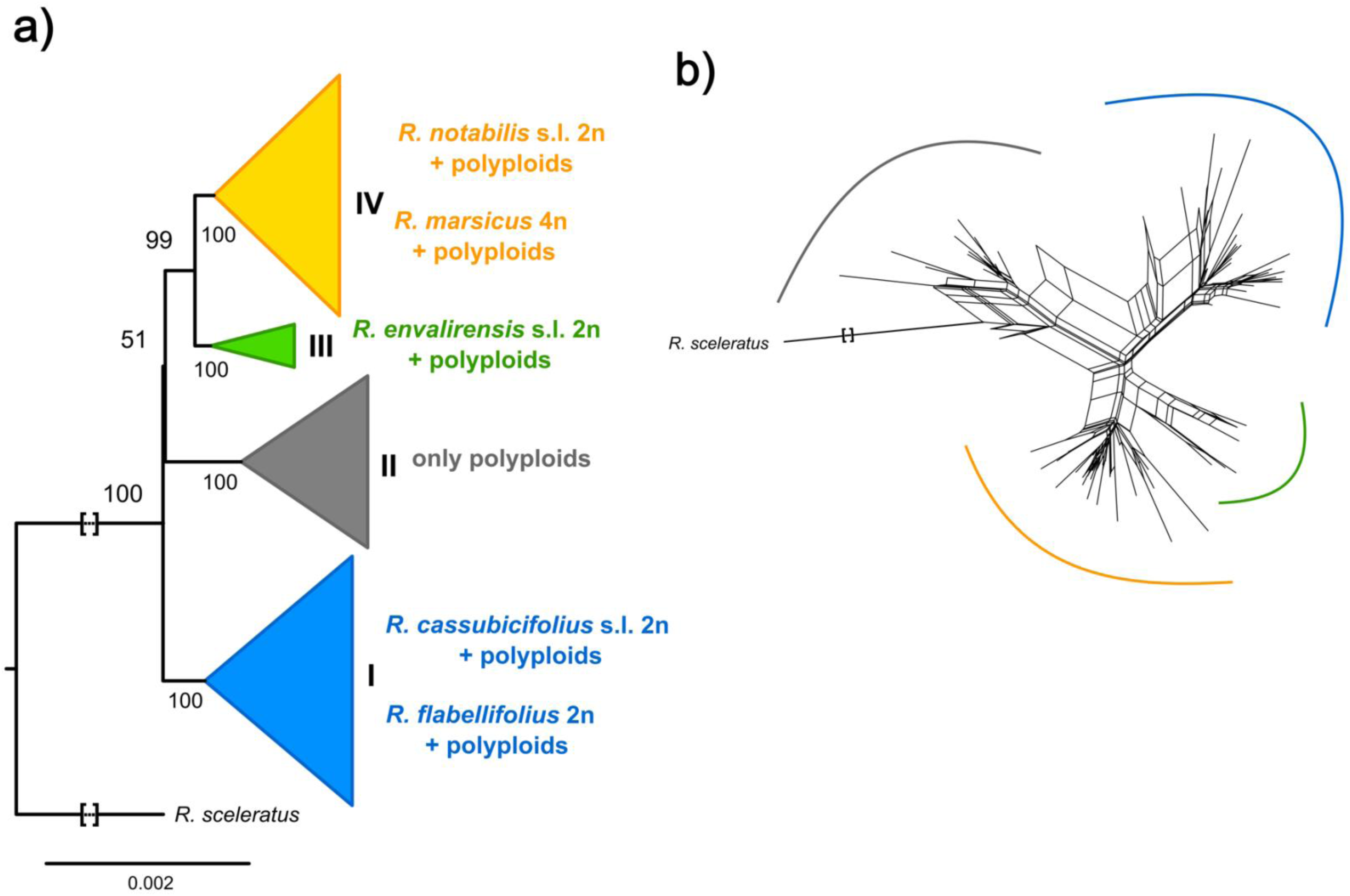
Phylogenetic tree and genetic structure based on plastome (CP) data. (a) ML tree (RAxML-NG) based on 87 samples and 71 plastid regions of the plastome (CP) dataset. Only main clades containing sexual species are shown (coloring according to Fig. 4). Concerning the clade in grey, plastid types of asexual polyploids were not found in any of the sexual species suggesting the former existence of a nowadays extinct sexual progenitor species. FBP values are given for each branch and clade (I-IV). (b) Neighbor-net analysis (SplitsTree) based on genetic distances (general time reversible [GTR] model with estimated site frequencies and ML), 87 samples, and 71 plastid regions of the plastome (CP) dataset. We colored main splits according to Fig. 4a. See Supplementary Fig. S5, S6 for more details. Squared brackets: A part of the branch was cut for illustrative purposes.

### Genetic Structure Analyses indicated 3-5 Clusters, strong Reticulation, and a Geographical Pattern

Structure analyses based on RADpainter+fineRADstructure revealed three supported main clusters (Fig. 6a). Sexual species were also clustered with polyploids: (I) *R. cassubicifolius* (C) and tetra- to hexaploid taxa, (II) *R. flabellifolius*, *R. marsicus*, and *R. notabilis* (F, M, and N) and tri-to hexaploid taxa, and (III) *R. envalirensis* (E) and tetraploid taxa. Commonly, polyploids showed high coancestry values, i.e., orange to red colors, with different clusters indicating reticulation events (see particularly polyploids H_1_–H_10_). In addition, highest values were found in relation to the sexual subgenomes occurring in the same cluster (Supplementary Table S7). Polyploids of cluster I showed highest similarity values with C and lowest ones with N and F. In contrast, cluster II is genetically more heterogeneous (subclusters IIa, IIa). Polyploids shared high similarity values with N and low coancestry values with F, E, and C. In cluster III, polyploids only exhibited high similarity to E.

**Fig. 6.**
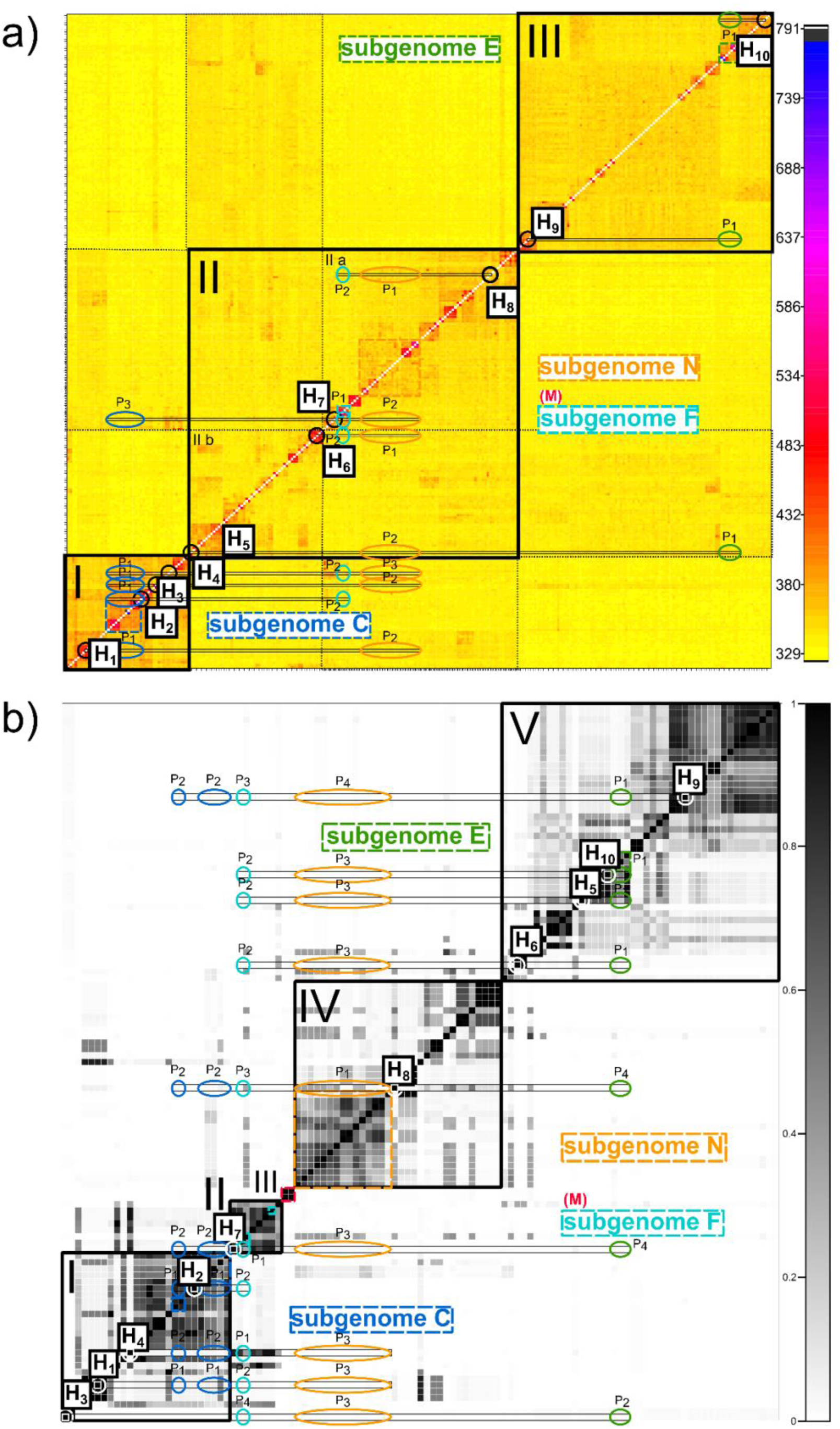
Genetic structure analyses based on RAD-Seq and target enrichment data. (a) Clustered fineRADstructure coancestry matrix of 280 sexual and polyploid apomictic individuals of the *R. auricomus* complex based on the ‘min10’ RAD-Seq alignment (97,312 loci, 438,775 SNPs). The legend on the right shows the color-coding of genetic similarity (coancestry values): the darker the square, the higher the similarity between a pair of individuals. (b) Similarity matrix of STACEY species delimitation analyses of 113 individuals based on a phased target enrichment alignment (48 genes). Posterior probabilities for belonging to the same cluster (species) are shown for pairs of individuals in the legend on the right: black is for 1.0 posterior probability and white for 0.0. See Fig. 4b, Supplementary Figs. S4a-c, S10 and high resolution figures on Figshare for clustering structure (a) tree with posterior probability group assignment probabilities and (b) coalescent-based tree. We indicated supported genetic clusters with solid lines, shared similarity among (sub)clusters with dotted lines, and sexual species with broad dashed colored squares (subgenomes C, F, M, N, and E). Small black squares (‘H_n_’) indicate selected tetraploid apomicts, which were investigated for allo- vs autopolyploid origin (10 polyploids; a small square indicates the analyzed individual; see IDs in Supplementary Table S1). Using lines and colored circles/ellipses, we highlighted the potential parental subgenome contributions for each polyploid (P_1_, P_2_, … and P_n_ with n=the n-th parental subgenome contribution. P_1_ is always the parental subgenome contribution with the highest coancestry score/posterior probability (likeliest) followed by other parental/subgenome contributions with decreasing coancestry scores/posterior probabilities (minor parental/subgenome contributions not drawn, see Supplementary Table S2 for more details).

Structure analysis based on STACEY revealed similar results (Fig. 6b, Supplementary Table S7). Sexuals are also surrounded by polyploids, and polyploids showed several reticulations and highest posterior probabilities with intra-cluster sexual subgenomes. There are few differences here: the former cluster II is divided into three distinct clusters each containing a single sexual species (II–IV), and many polyploids of the former cluster IIa are incorporated into cluster V. In addition, polyploids of cluster I also revealed significant posterior probabilities to E, and polyploids of cluster V to F, C, and N. In general, subgenome M shared no significant coancestry/posterior probability values with polyploids.

Structure analyses based on sNMF also unraveled three to four (up to five) main clusters (Fig. 7a–d, Supplementary Figs. S6–11, S15).

**Fig. 7.**
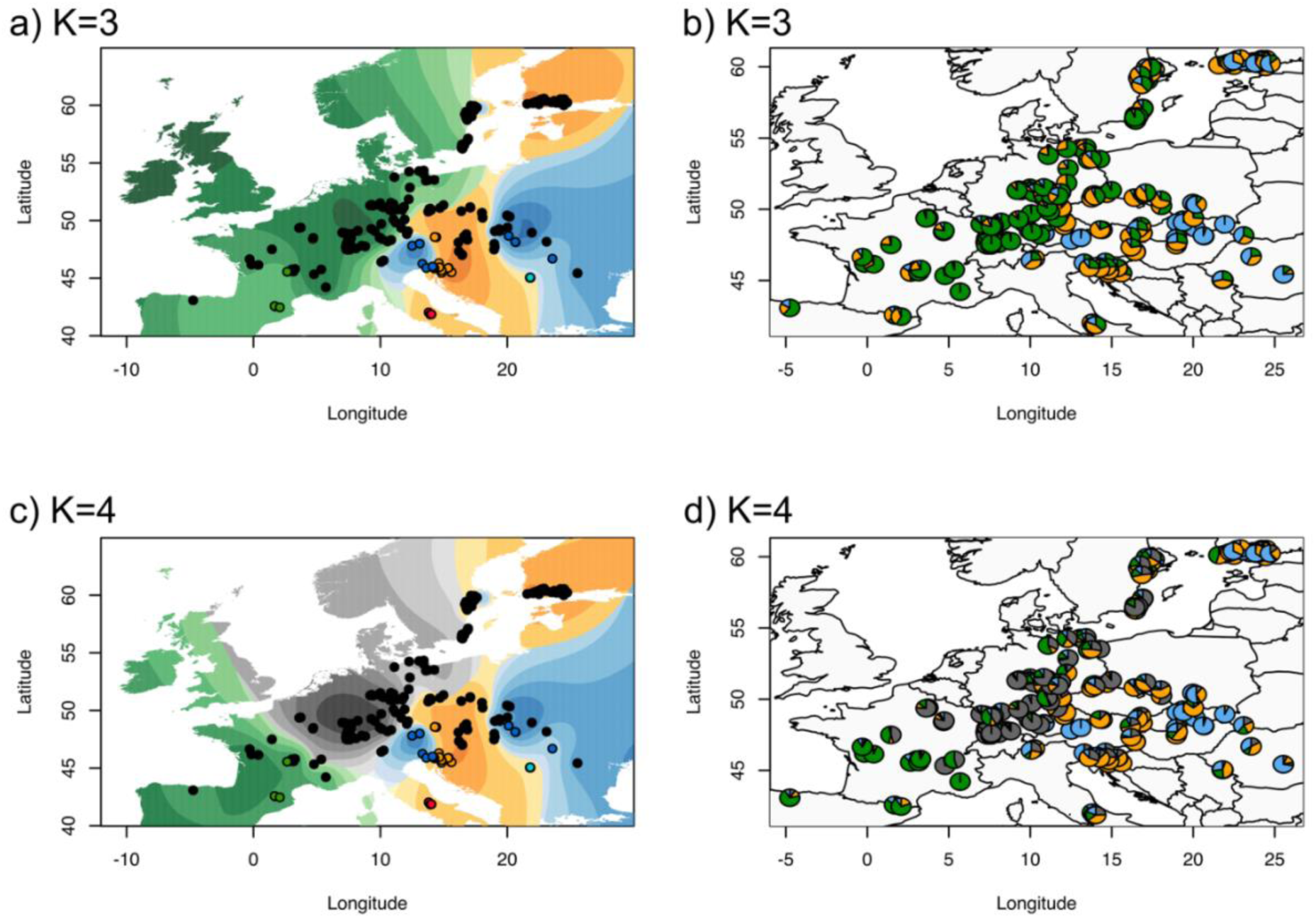
Geographic maps showing genetic clusters and ancestry coefficients across Europe. (a, c) interpolated values of ancestry coefficients (method ‘max’, i.e., at each point the cluster for which the ancestry coefficient is maximal) and (b, d) location-wise admixture estimate pie charts using K=3 and 4 genetic clusters. Results are based on sNMF results of 280 sexual and apomictic *R. auricomus* individuals and the ‘min30’ unlinked-SNP RAD-Seq alignment (33,165 loci). See Supplementary Figs. S6-11, S15, and figures on Figshare. In (a) and (c), colored circles represent sexual species (coloring according to Fig. 4): blue=*R. cassubicifolius* s.l. (C), turquoise=*R. flabellifolius* (F), red=*R. marsicus* (M), green=*R. envalirensis* s.l. (E), and orange=*R. notabilis* s.l (N). We adopted the coloring also to pie charts in (b) and (d). Europe map source: https://maps.ngdc.noaa.gov.

Although polyploids were characterized by a dominant genetic partition, they also showed 1–3 minor genetic partitions (Fig. 7b,d, Supplementary Fig. S7a–c). The likeliest number of K (clusters), K=3, showed a west-east distribution of clusters across Europe (Fig. 7a,c). The clusters themselves are north-south distributed. *Ranunculus envalirensis* and related polyploids (E, green partition) mainly inhabit regions in southwestern, central, and northern Europe. *Ranunculus flabellifolius*, *R. marsicus*, and *R. notabilis* and related polyploids (F, M, and N, orange partition) predominantly occupy southern, central-eastern, and northern Europe. *Ranunculus cassubicifolius* and related polyploids (C, blue partition) range from southeastern to northern Europe, including a disjunct distribution in central Europe. When comparing results of K=3 and K=4, the only remarkable difference is the emergence of a genetic cluster in central and northern Europe without a sexual species (grey partition) out of the former green one (Fig. 7a,c). In general, sNMF results are comparable to all previous analyses (grey partition predominantly found in clade V (Fig. 4a,b)/cluster III (Fig. 6a)/cluster V (Fig. 6b), and the orange partition mostly situated in clade II–IV (Fig. 4a,b)/cluster III (Fig. 6a)/cluster II–IV (Fig. 6b). The splitsgraph of the neighbor-net analysis (RAD-Seq) also exhibited three main genetic clusters weakly differentiated from each other (Supplementary Figs. S16a and Fig. 6a).

### Phylogenetic Networks supported by Genetic Structure revealed Allopolyploidy, 2-3 Contributing Subgenomes, and Subgenome Dominance

For most tested polyploids, phylogenetic networks based on RAD-Seq and target enrichment datasets showed two different subgenome contributions (Figs. 8a–h, 9, Tables 1, Supplementary Table S7). These polyploids were usually characterized by a dominant (P_1_=51–99%, mean 74%) and a minor subgenome contribution (P_2_=1–49%, mean 26%). Concerning PhyloNet likelihood+AIC calculations, reticulate evolution and thus allopolyploid origin was confirmed in most cases (H_1_–H_6_, H_8_, and H_10_). Within clade I and cluster I (Figs. 4, 6), polyploids H_1_–H_4_ possessed the dominant subgenome C whereas minor ones came from F followed by E and N. The blue haplotype C+F of these polyploids matched the dominant subgenome C. Final results indicated that ‘*R.* × *platycolpoides*’ (H_1_) is composed of subgenomes C and N, ‘*R.* × *elatior*’ (H_2_) of C and F, ‘*R.* × *pseudocassubicus*’ (H_3_) of C and E, and ‘*R.* × *hungaricus*’ (H_4_) of C and F.

**Fig. 8.**
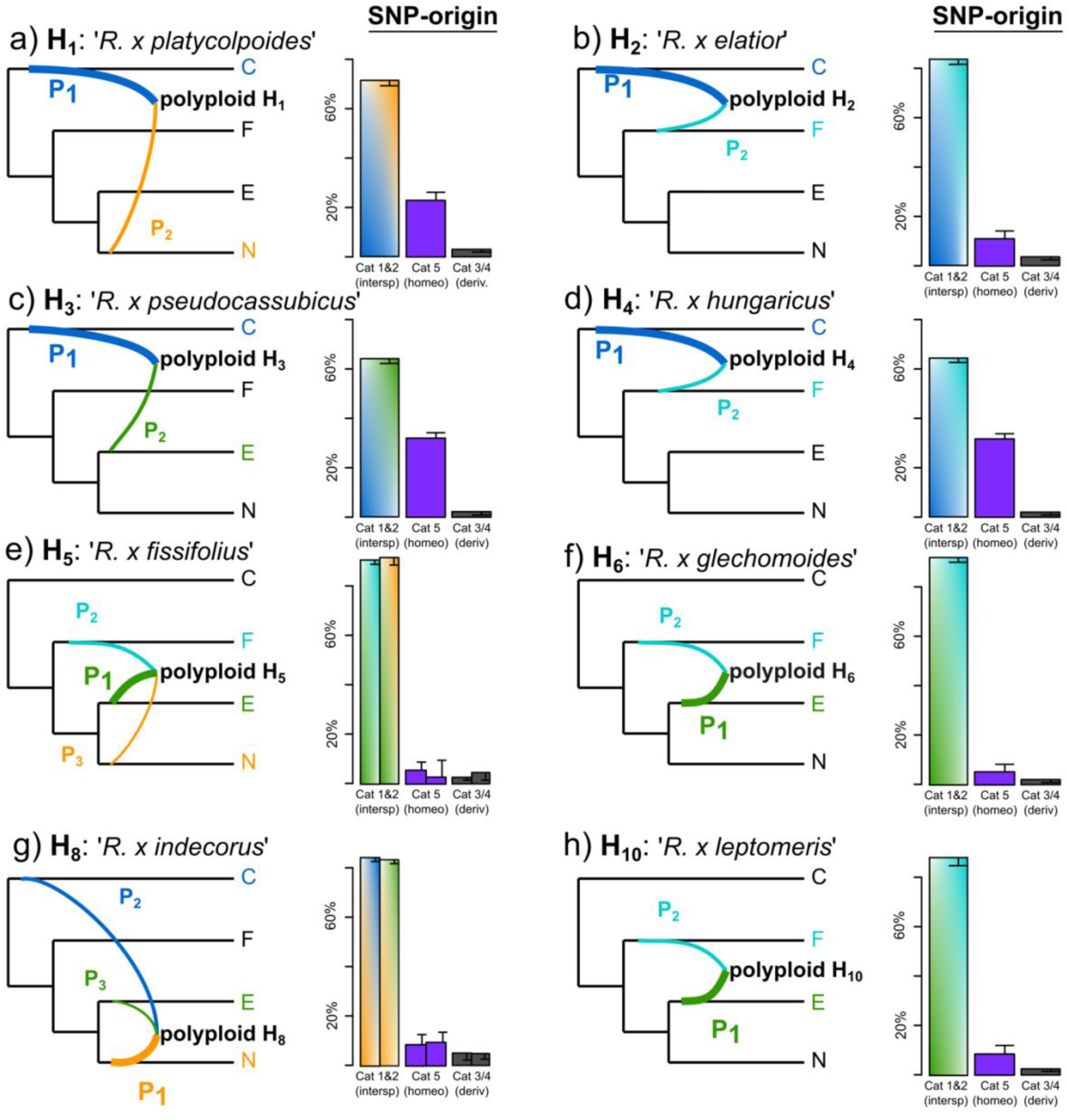
Reconstructed phylogenetic networks based on RAD-Seq, target enrichment, and CP data. (a-h, left) Final networks of allopolyploids are based on genetic structure and phylogenetic network results (consensus results) corrected by the by the full likelihood approach+AIC calculations in Phylonet and CP data. P_1_ defines the largest subgenome contribution, followed by P_2_ and P_3_. The network topology follows the published rooted phylogeny of *R. auricomus* sexuals (without tetraploid *R. marsicus*; Karbstein et al. 2020b). Curves indicate subgenome contributions (P_1_-P_3_). (a-h, right) Bar charts based on SNiPloid results are shown. Bar charts show SNP origins in percents (cat 1=SNPs identical to DIPLOID2, cat 2=SNPs identical to DIPLOID1/reference, cat 3/4=derived SNPs, cat 5=homeo-SNPs. We highlighted SNPs percent concerning all SNPs (see Material and Methods for additional evaluation of SNP category ‘others’) with additional black T-bars. Concerning H_5_ and H_8_, we calculated two SNiPloid analyses because three parents have contributed to its origin. Coloring of sexual progenitor subgenomes is according to Fig. 4. Subgenomes of C=*R. cassubicifolius* s.l., F=*R. flabellifolius*, E=*R. envalirensis* s.l., and N=*R. notabilis* s.l.

**Fig. 9.**
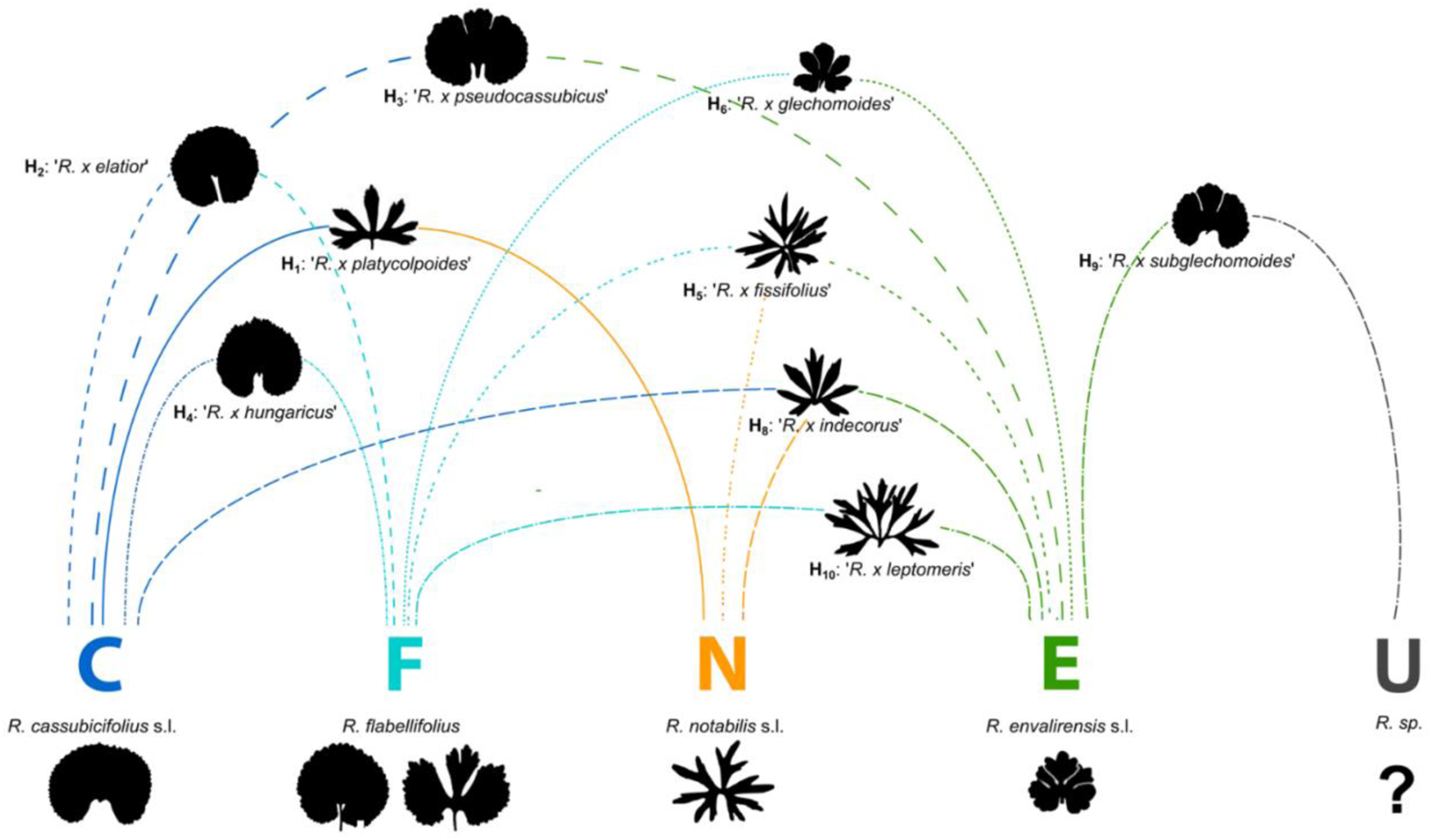
Hybrid scheme of sexual progenitors and selected polyploid *R. auricomus* derivates. The diploid sexual progenitor species *R. cassubicifolius* s.l. (C), *R. flabellifolius* (F), *R. notabilis* s.l. (N), *R. envalirensis* s.l. (E), and a hypothetical unknown one (U) in different combinations gave rise to asexual polyploid derivates (same polyploid individuals as in Fig. 8). Per allopolyploid, curves to the left and right indicate parental subgenome contributions. Subgenome dominance is shown by the relative position of the polyploid to the progenitors, for example ‘*R. x elatior*’ is closer to subgenome C due to C subgenome dominance). We also illustrate characteristic basal leaf types of taxa during anthesis (variation not covered, two types illustrated for *R. flabellifolius* due to the frequent occurrence of undivided and divided types; source: herbarium type specimens). The hybrid scheme is based on phylogenetic network and genetic structure results of RAD-Seq and target enrichment datasets supported by CP data (Table 1).

Moreover, we inferred varying subgenome contributions for the polyploid ‘*R.* × *pilisiensis*’ (H_7_) of clade II (Fig. 4a,b) and cluster II/III (Fig. 6a/b), but consensus results supported by CP results revealed subgenome F as the dominant one. The likeliest scenario is a tree-like evolution (autopolyploid origin). The polyploid ‘*R.* × *indecorus*’ (H_8_), positioned in clade IV (Fig. 4a,b) and cluster II/IV (Fig. 6a/b), showed three subgenomes, whereas N was the slightly dominant one, and C and N the minor ones. ‘*Ranunculus* × *fissifolius*’ (H_5_) was characterized by the orange haplotype N and is also composed of three different subgenomes (E, F, and N).

The polyploids ‘*R.* × *glechomoides*’ (H_6_), ‘*R.* × *subglechomoides*’ (H_9_), and ‘*R.* × *leptomeris*’ (H_10_) exhibited subgenome E as the dominant contribution. In most tree and genetic structure analyses, these polyploids were also situated close to E. CP analyses showed the green haplotype E for H_6_ and H_10_, but not for H_9_. Final results indicated E and F subgenome contributions for H_6_ and H_10_. ‘*Ranunculus* × *subglechomoides*’ (H_9_) exhibited the grey, unknown haplotype U and Phylonet AIC+likelihood calculations detected similarly-likely scenarios of reticulate (E and F) or tree-like evolution (E) (Table 1, Fig. 9).

### Post-origin Genome Evolution of Allopolyploids is shaped by Interspecific Gene Flow

SNP discovery (SNiPloid) based on RAD-Seq data supported allopolyploid hybrid origins with 3–33% (9–36% with evaluated others) homeo-SNPs of cat5. Whereas polyploids H_1_, H_3_, and H_4_ showed relatively high percentages of homeo-SNPs (>20%), the polyploids H_2_, H_4_–H_10_ exhibited low amounts (<15%). The majority of SNPs, however, indicated considerable post-origin evolution of allopolyploids (Fig. 8a–h, Supplementary Table S8). Across datasets, SNiPLoid assessed 64–93% (62–89% with evaluated ‘others’) interspecific SNPs of cat1&2, and 3–5% (2–3%) derived SNPs of cat3/4. Interspecific SNPs of cat1&2 were lowest for polyploids H_1_, H_3_, and H_4_ and highest for H_2_, H_4_–H_10_.

## Discussion

Polyploid phylogenetics is an emerging and bioinformatically challenging field, with important consequences for understanding plant speciation and macroevolution (Soltis et al. 2015; Landis et al. 2018; Rothfels 2021). Here, we used a comprehensive genomic, nuclear gene, and plastome dataset to unravel evolutionary processes of a less than 1 Mya polyploid species complex. Different kinds of evidence included in our self-developed bioinformatic pipeline confirmed that hybridization of sexual progenitors followed by polyploidization (allopolyploidy) is the dominant formation type in our model system (Table 1). Allopolyploidy also shaped the evolution of many other young polyploid complexes (Sochor, 2015; Spoelhof et al. 2017; Dauphin et al. 2018; Rothfels 2021). In addition, we also demonstrated remarkable post-origin genome evolution of allopolyploids, mostly due to interspecific gene flow. The bioinformatic pipeline presented here disentangled the parental contributions and the genomic diversity and composition of different polyploid apomictic lineages that have evolved. The *Ranunculus auricomus* model system is the first well-studied, large polyploid European species complex using OMICS data. The major conceptual breakthrough presented here is the combination of several datasets and up-to-date NGS methods into a new pipeline, starting with the diploid progenitors and ending up with the polyploid derivates, to receive, for the first time, a complete picture of the evolution of a large polyploid complex. The novel aspects particularly comprise network analyses, consensus result making of previous structure and network results, and auto- vs. allopolyploid testing.

### The Phylogenetic Pattern

Phylogenetic trees based on RAD-Seq and nuclear gene data surprisingly revealed only five well-supported main clades (Fig. 4a,b). Congruence between data sets hints at a strong evolutionary signal regardless of analyzing anonymous genomic regions (RAD-Seq) or only coding nuclear genes, and regardless of applying different analytical approaches (concatenation and ML vs. coalescence). Since the two marker sets complete each other to some extent (see Materials & Methods), we infer a robust phylogenetic framework for all further analyses.

Each main clade contained a sexual species surrounded by asexual polyploid lineages, showing that polyploids largely derived from their clade-specific progenitors (Fig. 4). This pattern is rather unique compared with other polyploid complexes, where usually clades with several diploid and (allo)polyploid taxa, but also clades with only polyploids were found (Kirschner et al. 2015; Dauphin et al. 2018; Carter et al. 2019; Wagner et al. 2020). Despite well-supported tree backbones according to BT values and highly informative branches (Fig. 4a), quartet support is partly low and distribution of alternative topologies (QD, Q’s; Fig. 4) hints at both inter-clade reticulation and ILS. Particularly within main clades, BT values are extremely low and quartet support metrics (QD, Q’s) indicate high rates of introgression and ILS signals (Supplementary Figs. S3, S4). Concerning the ML tree, TBE support was, especially at the backbone, higher than BT support probably due to less sensitivity of TBE to hybridization events and ‘jumping taxa’ (Lemoine et al. 2018). Reticulate evolution at the backbone of the tree is further indicated by the incongruent position of F polyploid derivatives in the plastid tree compared with the nuclear trees. The placement of diploid *R. flabellifolius* within the *R. cassubicifolius* plastid clade contrary to the nuclear trees suggests that *R. flabellifolius* could be an ancient homoploid hybrid species.

Within these main clades, the majority of described polyploid morphospecies are non-monophyletic (Supplementary Figs. S1–S4a–c). No clear clade-/group-specific morphological trend is recognizable (Figs. 4–9), except that taxa with undivided basal leaves were mainly found in clades I+II. This incongruence with morphology also rejects the old Linnaean classification of two main morphotypes (previously also rejected for sexual progenitors in Karbstein et al. 2020b). Network-like evolution through hybridization (allopolyploidy) is well-known to cause severe conflicting signals in tree reconstructions (Lemoine et al. 2018; Pease et al. 2018; Rothfels 2021). Moreover, cladogenetic speciation of *R. auricomus* allopolyploids from common polyploid ancestors is unlikely. Cladogenetic speciation would lead to bifurcating, tree-like post-origin evolution and thus only low conflicting signals (Jones 2017a) at middle and terminal branches. Here, these branches are extremely conflicting, suggesting extensive and repeated reticulate polyploid formation events. In contrast, coalescent-based phylogenetic analyses in the 20 Mya genus *Rubus* detected varying main clade positions, but highly resolved relationships within each clade (Carter et al. 2019). In brambles, ancient polyploidization events, strong geographical structure between continents, clade-specific and rather less post-origin evolution might explain this pattern. In *R. auricomus*, repeatedly ongoing hybridization and/or polyploidization of sexual progenitors during Pleistocene climate fluctuations in Europe and partly high facultative sexuality of polyploid apomicts in central and southern regions (Tomasello et al. 2020; Karbstein et al. 2021) potentially led to highly conflicting phylogenetic signals at middle and terminal branches.

### Genetic Structure, Origin, and Parentage of Polyploids

Genetic structure also revealed 3–5 main clusters (Fig. 6), largely fitting the phylogenetic main clades. We detected more reticulation signals in RAD-Seq data, probably favored by the incorporation of all SNPs of 97,312 loci, maximizing the information for analyzing young plant complexes. In contrast, analyses based on nuclear genes better inferred genetic boundaries by splitting each sexual progenitor species and accompanying polyploids into separate clusters. This advantage is probably related to the coalescent-based species delimitation using allelic information of the 48 phased loci (Jones 2017b; Karbstein et al. 2020b; Tomasello et al. 2020). Interestingly, we observed a west-east distribution of clusters within Europe (Fig. 7a, see also Karbstein et al. 2020b, 2021): a western cluster related to *R. envalirensis* (E), a central cluster related to *R. flabellifolius*, *R. marsicus*, and *R. notabilis* (F, M, and N), and an eastern cluster related to *R. cassubicifolius* (C), each ranging from southern to northern Europe, respectively. West-east allopatric speciation of sexual progenitors in combination with both allopolyploidization events and north-south migration of populations due to past climatic changes may explain this pattern (Abbott et al. 2013; Tomasello et al. 2020). Moreover, we detected a subdivision of the western cluster (green, Fig. 7c,d). The grey, central European subcluster widely corresponds to the grey ‘polyploid-only’ haplotype of the ML plastid tree (Fig. 5a). The fact that ploidy levels of *R. auricomus* populations in central Europe are well-studied (Jalas and Sumoninen 1989; Paule et al. 2018; Karbstein et al. 2021) makes it unlikely that an extant diploid was simply overlooked. Moreover, some polyploids possess a plastid type from an unknown diploid, suggesting an already extinct sexual progenitor related to subgenome E. Extinction of sexuals is a commonly considered or observed phenomenon in young polyploid complexes shaped by past climatic deteriorations (Sochor et al. 2015; Rothfels 2021) and is supported here by missing speciation events between 0.6 and 0.3 Ma (Million-years-ago; Tomasello et al. 2020). Alternatively, extinction could be due to past human activity.

Both genetic structure and phylogenetic network results based on RAD-Seq and nuclear gene data revealed that the majority of polyploids were composed of two, surprisingly some of three, subgenome contributions (Figs. 6, 8, 9, Table 1). Plastome data underlined inferred subgenome contributions. We detected only one polyploid, that didn’t show evidence of a reticulate evolutionary history: ‘*R. × pilisiensis*’ (H_7_). Here, varying subgenome contributions were found (Fig. 6, Table 1), but network analyses suggested autopolyploid origin from progenitor subgenome F. This lineage might also represent a segmental allopolyploid, as auto- and allopolyploidy are connected by transitions (Comai 2005; Spoelhof et al. 2017). Autotetraploid cytotypes are also known from the otherwise diploid sexual species *R. cassubicifolius* (Hörandl and Greilhuber 2002). However, autopolyploidy is currently only present in the *R. auricomus* complex in clades/clusters I and II, supporting its probably less frequent occurrence compared with allopolyploidy in nature (Spoelhof et al. 2017; but see many unnamed autopolyploids in Barker et al. 2015).

All diploid sexual progenitors were involved in allopolyploid formation (C, F, N, E, and U; Fig. 9). Polyploids were probably formed multiple times out of different progenitor combinations followed by considerable post-origin evolution. Close crosses result more easily in homoploids, whereas distant ones tend to become rather allopolyploid (Soltis and Soltis 2009). However, we observed allopolyploids out of genetically distant (C+N, N+C+E), moderate (C+E, E+F+N), and close (C+F, E+F) crosses. Extant homoploid hybridization is probably inhibited by the allopatric distribution of sexual species (see Tomasello et al. 2020). Interestingly, allopolyploids showed more meiotic errors than homoploid hybrids in experimental crossings (distant C+N *R. auricomus* crosses; Barke et al. 2020). However, polyploids can escape hybrid sterility via apomixis, vegetative reproduction, and/or selfing (Hörandl 2006; Soltis and Soltis 2009; Barke et al. 2018, 2020).

Extant sexuals have restricted ranges and are separated by thousands of kilometers across Europe (Fig. 2). Nevertheless, all main genetic clusters are present in central Europe (Fig. 7a, c). Sexual progenitors might have repeatedly met in this region during past interglacial times, giving rise to multiple allopolyploidization ‘waves’ with varying subgenome contributions. Interestingly, polyploids composed of three different subgenomes were only found in Central Europe and South Sweden (H_5_, H_8_), underlining the importance of secondary contact zones for the allopolyploid origin of the *R. auricomus* complex.

The genetic and phenotypic diversity of *R. auricomus* biotypes with more than 840 described morphospecies is probably formed by multiple allopolyploidization events from four extant and at least one probably extinct diploid, sexual progenitor species. Other studies already demonstrated that few diploid progenitors were capable of producing a magnitude of allopolyploids, for example in *Botrychium* (Dauphin et al. 2018), *Rubus* (Sochor et al. 2015; Carter et al. 2019), or *Taraxacum* (Kirschner et al. 2015).

### Post-Origin Evolution: Subgenome dominance, Hybridization, and Mutation

Subgenome dominance was inferred in almost all tested allopolyploids and resulted in 74% mean inheritance probabilities from network analyses (Figs. 6, 8, 9, Table 1). Per main genetic cluster, allopolyploids were usually composed of dominant intra-cluster subgenomes and 1(−2) minor, varying inter-cluster subgenome(s), although trigenomic polyploids showed rather similar contributions (supported by SNP-origin analyses; Figs. 6, 8). Plastid data supported the dominant subgenome contribution and, in some cases, unraveled an additional or unknown/extinct subgenome (Table 1). Sequence subgenome dominance probably leads to the grouping of polyploids close to the dominant progenitor (Figs. 4, 6). Hence, polyploids of cluster I, II (II–IV), and III probably received at least one subgenome from C, N, and E(-like) progenitor lineages.

Subgenome dominance is a common feature of allopolyploids (Blischak et al. 2018; Alger and Edger 2020). Here, segregation after hybridization, and after polyploidization, gene flow due to facultative sexuality of apomicts might cause the observed dominance. We consider biased fractionation of minor importance regarding the less than 1 Mya or even younger *R. auricomus* polyploids. For example, similar old *Brachypodium hybridum* showed only gradual gene loss and no subgenome expression dominance 1.4 million years after allopolyploidization (Gordon et al. 2020). The importance of diploid hybrid segregation and post-origin gene flow due to facultative sexuality is highlighted by SNiPloid analyses. We detected only a minority of SNPs from hybridogenic origins (3–36%), but considerable proportions of interspecific, post-hybridization SNPs (62–93%). Since apomixis establishes only stepwise, the first diploid hybrid generations still exhibit predominant sexual reproduction (Barke et al. 2018), allowing for Mendelian segregation. The potential of Mendelian segregation in early hybrid generations for creating the nearly complete extant morphological diversity of the complex was demonstrated experimentally by Barke et al. (2018) and Hodač et al. (2018).

Moreover, facultative sexuality and maintenance of functional pollen probably allowed the newly formed hybrids for backcrossing with their parental species, and among each other. Extant polyploids under natural conditions exhibit usually low, but varying degrees of facultative sexuality (mean 2.15%, range 0–34%; Karbstein et al. 2021). Relatively high sexuality values of central and southern European asexual populations may indicate some ongoing gene flow. For example, in apomictic *Rubus* or *Pilosella* polyploids, which are also characterized by highly variable, facultative sexuality, post-origin scenarios by interlineage gene flow and backcrossing to sexual parents have also been considered (Sochor et al. 2015; Hörandl 2018; Nardi et al. 2018; Carter et al. 2019). However, processes of fast segregation with backcrossing before vs. gene flow after polyploidization cannot be distinguished here, but we recognize here post-origin genome evolution as an important factor shaping genome structure of polyploids.

Only a few morphospecies appeared as monophyletic groups (e.g., H_2_, H_6_, H_7_; see also Supplementary Figs. S1–S3) and are evolving towards more stable lineages. The original hypothesis of Babcock and Stebbins (1938) predicted that such lineages would only form at higher ploidy levels (>*6n*, see Fig. 1). However, already Grant (1981) recognized that formation of stable apomictic lineages in a ‘mature complex’ is less dependent on cytotype but rather correlated to age, loss of sexuality, and extinction of sexual progenitors. According to Grant’s (1981) definition, the *R. auricomus* complex is in an early mature stage of evolution, with extant diploid progenitors and a broad array of apomictic biotypes. This hypothesis is confirmed by the low proportion of lineage-specific SNPs (Fig. 8). SNP origin analyses revealed only 2–5% derived SNPs in allopolyploids attributable to mutations (Welch and Meselson 2000; Pellino et al. 2013). Low degrees of post-origin sequence evolution are not surprising when comparing the evolutionary young *R. auricomus* polyploids to similar-old or older allopolyploids (Pellino et al. 2013; Gordon et al. 2020; Tomasello et al. 2020; Wagner et al. 2020). For example, several million years old *Salix* sexual allopolyploids exhibited 19–47% post-origin, species-specific SNPs (Wagner et al. 2020).

### Integration of Datasets and Analyses for unraveling Evolutionary Processes in Young Polyploid Complexes

Our case study demonstrates that even with OMICS approaches it is useful to rely on different complementary reduced-representation datasets to tackle polyploid complexes: genomic RAD-Seq, nuclear genes, and plastomic regions. On the one hand, RAD-Seq provided the highest number of (allelic) information (loci and SNPs) both from non-coding and coding regions. On the other hand, correct allele phasing and discrimination of homoeologues is desirable for polyploids (Eriksson et al. 2018; Freyman et al. 2020; Lautenschlager et al. 2020; Rothfels 2021) but still a challenge for non-model plants. Both datasets together represent well the nuclear genome and can be much easier collected, cheaper sequenced, and easier analyzed for a large number of samples than entire transcriptomes or genomes (McKain et al. 2018; Johnsen et al. 2019). Moreover, target enrichment even allows the inclusion of young to old herbarium-type material (here, up to 74-years-old; up to 204 years in Brewer et al. 2019). This is particularly important in times of traveling restrictions and crucial for the correct application of taxon names in extremely morphologically diverse species complexes.

Here, we combined the advantages of three datasets to unravel evolutionary processes in polyploid complexes. We confirm previous approaches (e.g., Lo et al. 2010; Brandrud et al. 2020) demonstrating that a combination of tree building, structure, and network analyses is most useful to reconstruct non-hierarchical relationships in these complexes. The sexual progenitor species often diversify in a rather tree-like bifurcating manner that can be recognized with tree-building supported by genetic structure and/or morphometric methods (Burgess et al. 2015; Wagner et al. 2019; Karbstein et al. 2020b).

In our study, we demonstrate that the RAD-Seq ML tree revealed a highly congruent topology compared with the target enrichment nuclear gene coalescent-based tree. These trees gave a first phylogenetic framework for the *R. auricomus* complex, as also shown in evolutionary young polyploid complexes of *Crataegus* (Lo et al. 2010) or *Dactylorhiza* (Brandrud et al. 2020). Here, the low interclade and extremely low intraclade (quartet) support values indicated the presence of reticulations and/or ILS. Nevertheless, allopolyploids originated by diverged progenitor species introduce errors in ordinary tree reconstructions due to network-like evolution and smushing of different evolutionary histories in consensus sequences (McDade 1992, Oxelman et al. 2017, Rothfels 2021). We regard this issue in our study as minor, since progenitors of polyploids are genetically less diverged (Karbstein et al. 2020b), tree and genetic structure analyses show comparable results, and subgenome dominance of allopolyploids was probably also expressed in consensus sequences used for tree analyses.

The applied approaches provide a phylogenetic framework for recognizing sexual progenitor species and thus for unraveling the origins of (allo)polyploids. The detection of incongruences between plastid trees and nuclear datasets is a strong signal for hybridization events already on the diploid level (McKain et al. 2018; Dauphin et al. 2018). In this study, incongruences delivered valuable information for both sexual progenitors and polyploid derivates (see e.g., parental contribution of a probably already extinct species in H_9_). However, for the reconstruction of the reticulate relationships of allopolyploids, only genetic structure and network methods are able to unravel the correct evolutionary relationships.

Since neopolyploid complexes are evolutionary young and are characterized by reticulations and ILS, it is useful to employ genetic structure analyses that incorporate a maximum of allelic sequence diversity information. These analyses were previously often conducted with DNA fingerprinting markers (e.g., microsatellites, AFLPs). AFLP markers do not inform about heterozygosity, and in polyploids also microsatellites are usually scored as presence/absence, because allele dosages can often not be reliably assessed (e.g., Hodac et al. 2018; Karbstein et al. 2019; Melicharkova et al. 2020). RADseq covers a magnitude of markers, providing genome-wide sequence diversity per locus (SNPs) and thus robust results for genetic structure. RADpainter+fineRADstructure incorporates all SNPs, and varying allele numbers and amounts of missing data appropriate for young polyploid analyses (Malinsky et al. 2018; Wagner et al. 2021). In addition, the employed sNMF algorithm based on unlinked SNPs is not only faster, but also less sensitive to deviations from Hardy-Weinberg equilibrium (HWE) than the popular STRUCTURE software; it tolerates missing data, and is also applicable to different ploidy levels (Frichot et al. 2014; Frichot and François 2020; Karbstein et al. 2021).

Whereas these methods impressively showed hybridity and a not yet recognized species gene pool of apomicts, coalescent-based STACEY species delimitation based on phased nuclear genes more clearly delimited the genetic structure of the polyploid complex (Fig. 6a,b). Allele phasing was demonstrated and is particularly considered as crucial for resolving young, reticulate relationships (Andermann et al. 2018; Eriksson et al. 2018; Freyman et al. 2020; Rothfels 2021). Moreover, phylogenetic network analyses and subsequent tests mainly based on phased nuclear genes (see also Tiley et al. 2021) best unraveled subgenome contributions per polyploid and demonstrated predominant allopolyploid origins. Performing several network methods across different datasets informed by plastid information (i.e., consensus making) is the most important part of our study to get a reliable picture about polyploid evolution in such a young complex. Nevertheless, in general, potential limitations are the not realizable correct allele phasing of short RAD-Seq loci and relatively low number of nuclear genes, which were compensated by the combination of both datasets.

Disentangling genetic markers (SNPs) of polyploids for post-origin processes informs about divergence and stability of lineages. This information is crucial for classification and delimitation of species (Grant 1981; Hörandl 2018). Although incorporating ten thousands of RAD-Seq loci, our SNiPloid analyses assigned only a minor fraction of RAD-Seq-SNPs to homoeologous SNPs derived from hybrid origin. Here, a similar approach that incorporates homeologs (from various ploidy levels) derived from phased nuclear genes would be more favorable to assess polyploid (post-)origin evolution. A limitation of the SNiPloid pipeline is that the parental species must be defined for the input, only single samples can be analyzed, and that the algorithm is so far limited to tetraploids (i.e., not applicable to higher or lower ploidy levels; see also Wagner et al. 2020). However, the congruence of our results in eight independently analyzed hybrid lineages indicates two major trends in *R. auricomus*, namely considerable segregation of the diploid hybrid generation combined with gene flow after polyploidization, and so far only a low divergence via mutation in the more or less stable lineages.

Using the gained knowledge of this study, i.e., potential progenitor species of (allo)polyploids, ploidy levels and reproduction modes, and allo- vs. autopolyploid origins, subgenome assignments of allopolyploids and more appropriate phylogenetic allopolyploid networks (e.g., Jones 2017a; Cao et al. 2019; Lautenschlager et al. 2020; Šlenker et al., 2021) are applicable or should be optimized for the polyploid complex. The combination of datasets and analytical pipelines gives a more comprehensive and complete picture of the evolution of young polyploid complexes.

## Data availability

The authors declare that basic data supporting the findings are available within the manuscript and Supporting Information. RAD-Seq, target enrichment, and CP alignments, and tables and figures supporting the results are deposited on FigShare (https://doi.org/10.6084/m9.figshare.14046305). RAD-Seq reads are deposited on the National Center for Biotechnology Information Sequence Read Archive (SRA): BioProject ID PRJNA627796 http://www.ncbi.nlm.nih.gov/bioproject/627796). Flow cytometric (FC) and flow cytometric seed screening (FCSS) data are also stored in Figshare (https://doi.org/10.6084/m9.figshare.13352429).

## Code availability

We deposited custom bash, R, and Julia scripts on Github (https://github.com/KK260/Ranunculus_auricomus_phylogenetic_network_scripts).

## Acknowledgments

We acknowledge Franz G. Dunkel for providing garden plants and herbarium specimens, Ena Lehtsaar and Julius Schmidt for technical help, and John Paul Bradican for suggestions on previous manuscript versions. We thank the herbaria of Jena (JE), Munich (M), Oslo (O), Uppsala (UPS), and the University of Vienna (WU) for loans of *R. auricomus* type species material. We thank three referees and the editors for valuable comments on the manuscript.

## Author contribution

K.K., S.T., and E.H. designed research; K.K., L.H., and E.H. collected plant materials; K.K., S.T., P.M., B.B.H., and C.P. performed lab work; K.K., S.T., and N.W. analyzed data; K.K. wrote the paper with contributions of all authors.

