## Supplementary for "Unraveling Phylogenetic Relationships, Reticulate Evolution, and Genome Composition of Polyploid Plant Complexes by Rad-Seq and Hyb-Seq"

**Supplementary Table S1 (Excel sheet).** Location details of sampled *R. auricomus* populations across Europe. We investigated 235 sexual and apomictic populations (see also Fig. 2). Population ID, taxon name, ploidy, main reproduction mode (see also Karbstein et al. 2021, assumed reproduction modes due to missing data in brackets), locality by country and ISO code 3166-2, collection date, altitude (in meter above sea level, m.a.s.l.), latitude (N, decimal), longitude (E, decimal), habitat, collector, herbarium voucher specimens ([#]=holotype, [+]=private herbarium F.G.Dunkel), and available samples per dataset (RAD-Seq, TE=target enrichment dataset, and CP=plastome dataset) and individuals in network analyses (tested polyploids) are given.

**Supplementary Text S1.** HybPhyloMaker settings (target enrichment analysis).

Sequence adapters were removed, and reads were quality-trimmed using Trimmomatic vers. 0.32 (Bolger et al. 2014), with the default settings implemented in HybPhyloMaker vers. 1.6.4. Duplicated reads were removed with FastUniq vers. 1.1 (Xu et al. 2012). We used the concatenated sequences of target exons separated by stretches of 800 Ns as 'pseudo-reference' for read mapping with BWA vers. 0.7.12 (Li and Durbin 2010). Consensus sequences of mapped reads were produced with ConsensusFixer vers. 0.4 (available at: https://github.com/cbg-ethz/ConsensusFixer), as this is the only approach available in HybPhyloMaker for calling ambiguity DNA codes in case of multiple bases per site in the mapped reads. For ConsensusFixer, we used the following settings: minimum relative abundance of the alternative base ('plurality' in the settings file of HybPhyloMaker) of 0.2 and a minimum read coverage for ambiguity calling ('mincov') of 5.

Consensus sequences were matched to sequences of the target exons to produce *.pslx files using BLAT vers. 35.1 (Kent 2002), and combined across samples to produce exon-wise matrices with 'assembled_exons_to_fastas.py' (Weitemier et al. 2014). Matrices were aligned with MAFFT vers. 7.029 (Katoh and Standley 2013), using the default program settings, and then gene-wise concatenated with AMAS vers. 1.0 (Borowiec 2016). We excluded samples with more than 40% of missing data from each exon region, and subsequently, retained only exons including more than 90% of samples.

**Supplementary Table S2 (Excel sheet).** Quality trimming and read mapping (target enrichment analysis). Detailed results of quality and duplicate trimming (Trimming) and read mapping to the ‘pseudo-reference’ consisting of concatenated target regions (Mapping).

**Supplementary Table S3 (Excel sheet).** Loci selection (target enrichment analysis). We scored exon regions to select the 50 best loci to be phased and used in network analyses (see Materials for details on how the scoring was performed). 'R_squared' defines the $R^2$ of mutational saturation regression curves, and 'LBscoresSD' describes the standard deviation of the sample-specific long-branch scores (LB scores). The last column gives information on the amount of parsimony informative sites in each exon region. The selected loci are highlighted in pink.

**Supplementary Table S4 (Excel sheet).** Results of the phasing procedure (target enrichment analysis). Information is given on the number of alleles found for each sample in the 50 loci selected for the species delimitation and network analyses. The last column gives information on the best-fitting model of sequence evolution found with ModelTest-NG.

**Supplementary Table S5 (Excel sheet).** Read mapping (plastome analysis). Detailed results of the read mapping to the chloroplast reference genome. We used the *Ranunculus repens* plastid genome (GenBank accession number NC_036976; Dann et al. 2017) as assembly reference.

**Supplementary Table S6 (Excel sheet).** List of regions used for the chloroplast phylogeny (plastome analysis). The alignments were selected excluding first samples with more than 50% missing data and second alignments including less than 50% of samples. From those selected alignments (71 in total), we excluded samples missing from more than 50% of the alignments (marked in pink). Numbers indicate percentages of missing data (0=no missing data; empty cell=completely missing).

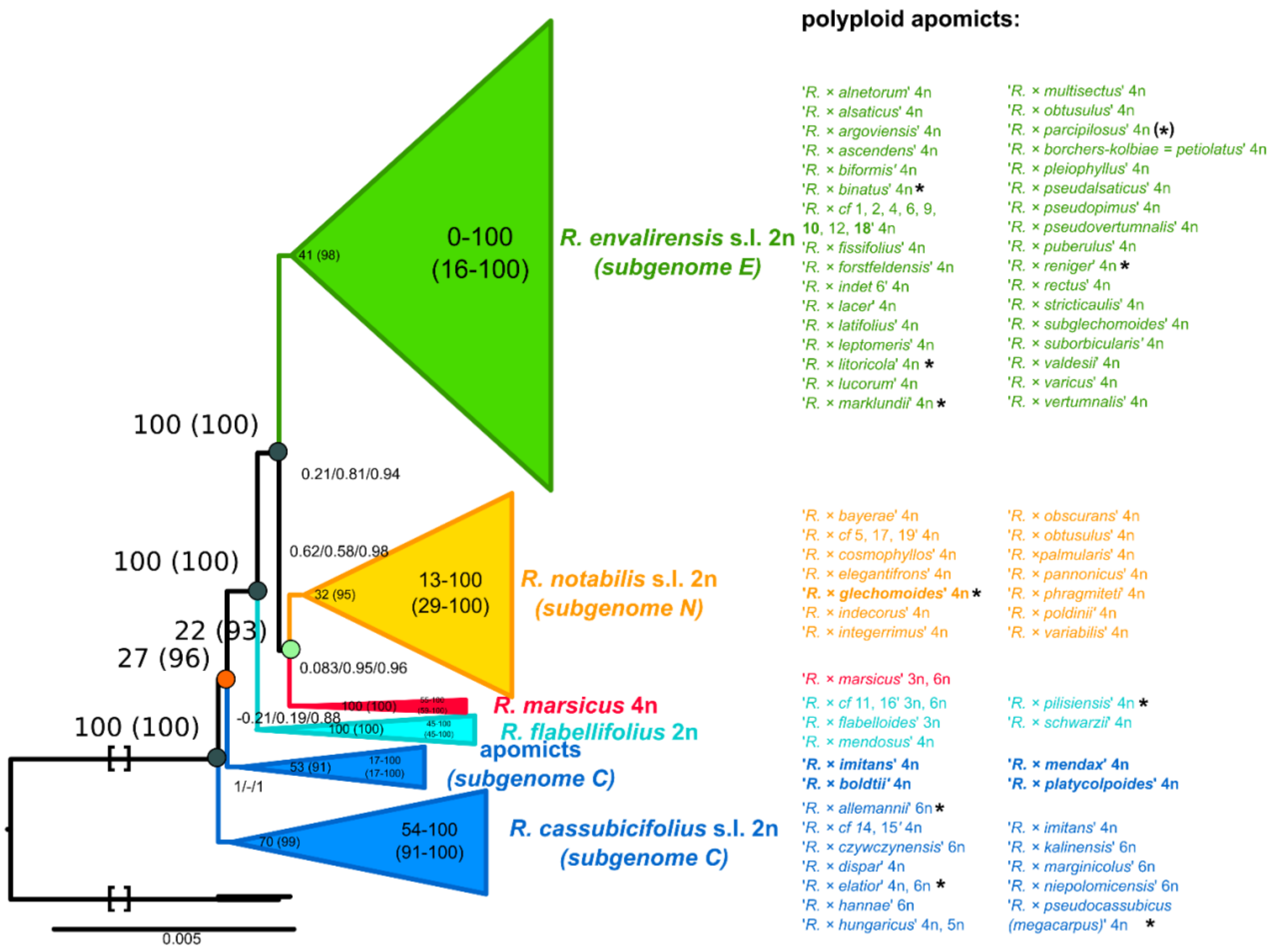


**Supplementary Fig. S1.** Concatenated ML tree with summarized main clades (min30). Results are based on RAxML-NG results and the 'min30' RAD-Seq alignment (280 samples, 33,165 loci, 194,083 SNPs). Felsenstein Bootstrap Proportions (FBP) and Transfer Bootstrap Expectation (TBE, in brackets) values are displayed on branches. Range of BT values (i.e., FBP (TBE) value range of all internal nodes) is shown for each main clade. Names of sexual species and apomictic taxa resolved in the main clades are listed to the right in corresponding colors, with samples resolved differently compared to the ‘min10’ ML tree in bold. Ploidy levels are listed to the right of species/taxon names. See Figshare data repository for the complete ML tree. *=monophyletic polyploid apomictic taxon. Squared brackets: A part of the branch was cut for illustrative purposes.

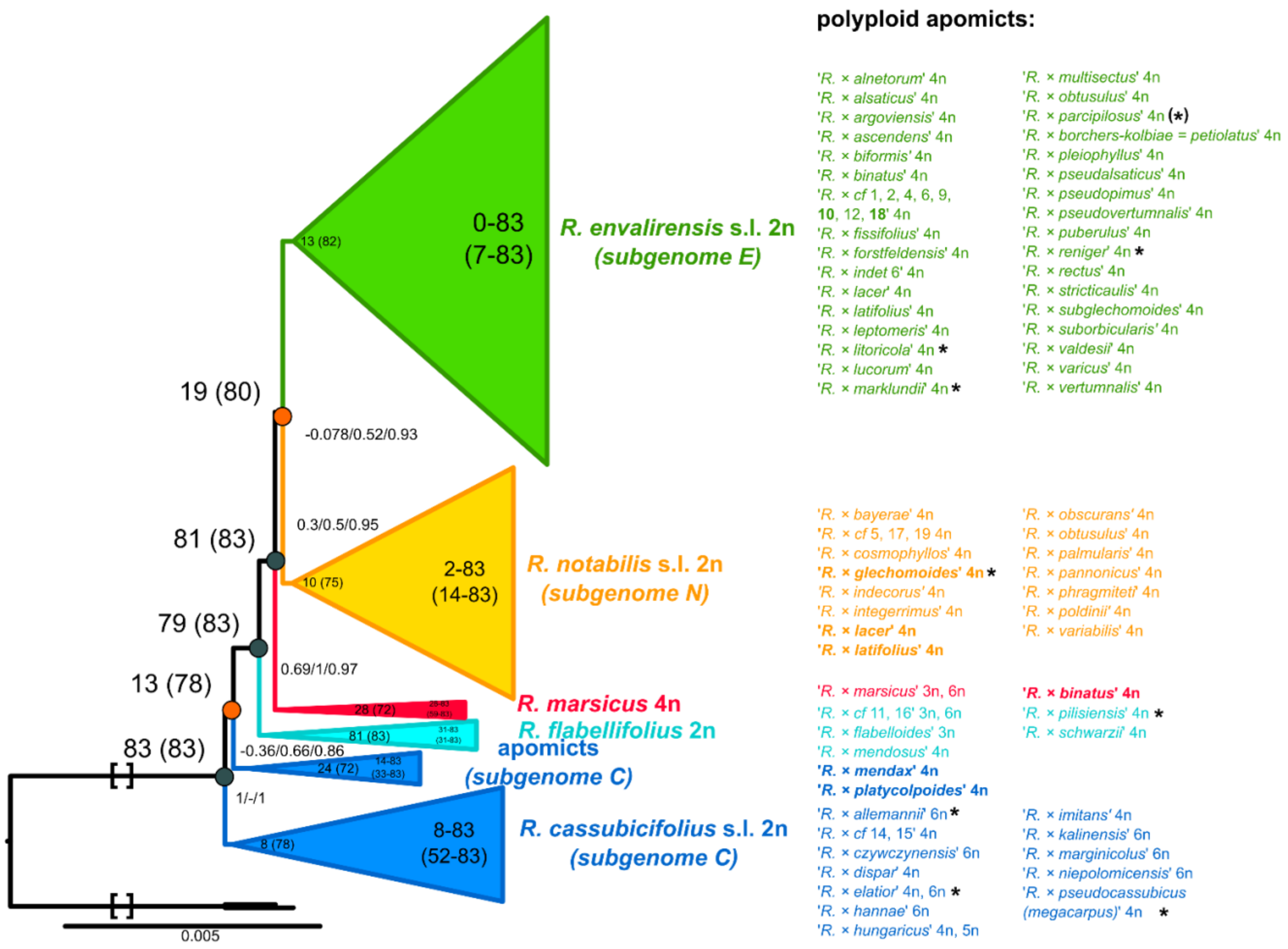


**Supplementary Fig. S2.** Concatenated ML tree with summarized main clades (min50). Results are based on RAxML-NG results and the 'min50' RAD-Seq alignment (280 samples, 11,196 loci, 64,554 SNPs). Felsenstein Bootstrap Proportions (FBP) and Transfer Bootstrap Expectation (TBE, in brackets) values are displayed on branches. Range of BT values (i.e., FBP (TBE) value range of all internal nodes) is shown for each main clade. Names of sexual species and apomictic taxa resolved in the main clades are listed to the right in corresponding colors, with samples resolved differently compared to the ‘min10’ ML tree in bold. Ploidy levels are listed to the right of species/taxon names. See Figshare data repository for the complete ML tree. *=monophyletic polyploid apomictic taxon. Squared brackets: A part of the branch was cut for illustrative purposes.

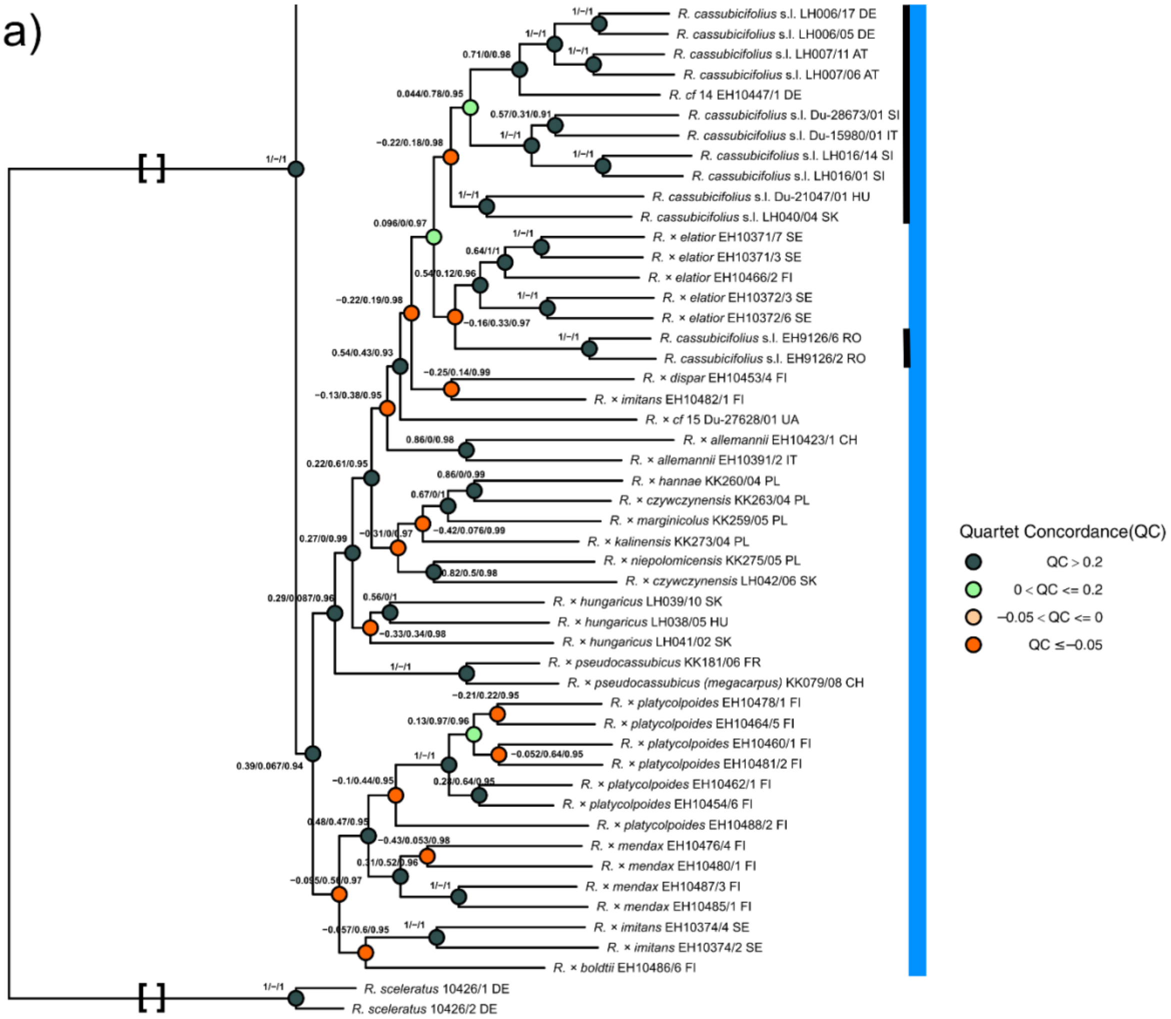

a)
R. cassubicifolius s.l. LH006/17 DE
R. cassubicifolius s.l. LH006/05 DE
R. cassubicifolius s.l. LH007/11 AT
R. cassubicifolius s.l. LH007/06 AT
R. cf 14 EH10447/1 DE
R. cassubicifolius s.l. Du-28673/01 SI
R. cassubicifolius s.l. Du-15980/01 IT
R. cassubicifolius s.l. LH016/14 SI
R. cassubicifolius s.l. LH016/01 SI
R. cassubicifolius s.l. Du-21047/01 HU
R. cassubicifolius s.l. LH040/04 SK
R. × elatior EH10371/7 SE
R. × elatior EH10371/3 SE
R. × elatior EH10466/2 FI
R. × elatior EH10372/3 SE
R. × elatior EH10372/6 SE
R. cassubicifolius s.l. EH9126/6 RO
R. cassubicifolius s.l. EH9126/2 RO
R. × dispar EH10453/4 FI
R. × imitans EH10482/1 FI
R. × cf 15 Du-27628/01 UA
R. × allemannii EH10423/1 CH
R. × allemannii EH10391/2 IT
R. × hannae KK260/04 PL
R. × czywczynensis KK263/04 PL
R. × marginicolus KK259/05 PL
R. × kalinensis KK273/04 PL
R. × niepolomicensis KK275/05 PL
R. × czywczynensis LH042/06 SK
R. × hungaricus LH039/10 SK
R. × hungaricus LH038/05 HU
R. × hungaricus LH041/02 SK
R. × pseudocassubicus KK181/06 FR
R. × pseudocassubicus (megacarpus) KK079/08 CH
R. × platycolpoides EH10478/1 FI
R. × platycolpoides EH10464/5 FI
R. × platycolpoides EH10460/1 FI
R. × platycolpoides EH10481/2 FI
R. × platycolpoides EH10462/1 FI
R. × platycolpoides EH10454/6 FI
R. × platycolpoides EH10488/2 FI
R. × mendax EH10476/4 FI
R. × mendax EH10480/1 FI
R. × mendax EH10487/3 FI
R. × mendax EH10485/1 FI
R. × imitans EH10374/4 SE
R. × imitans EH10374/2 SE
R. × boldtii EH10486/6 FI
R. sceleratus 10426/1 DE
R. sceleratus 10426/2 DE
Quartet Concordance(QC)
QC > 0.2
0 < QC <= 0.2
−0.05 < QC <= 0
QC ≤ −0.05

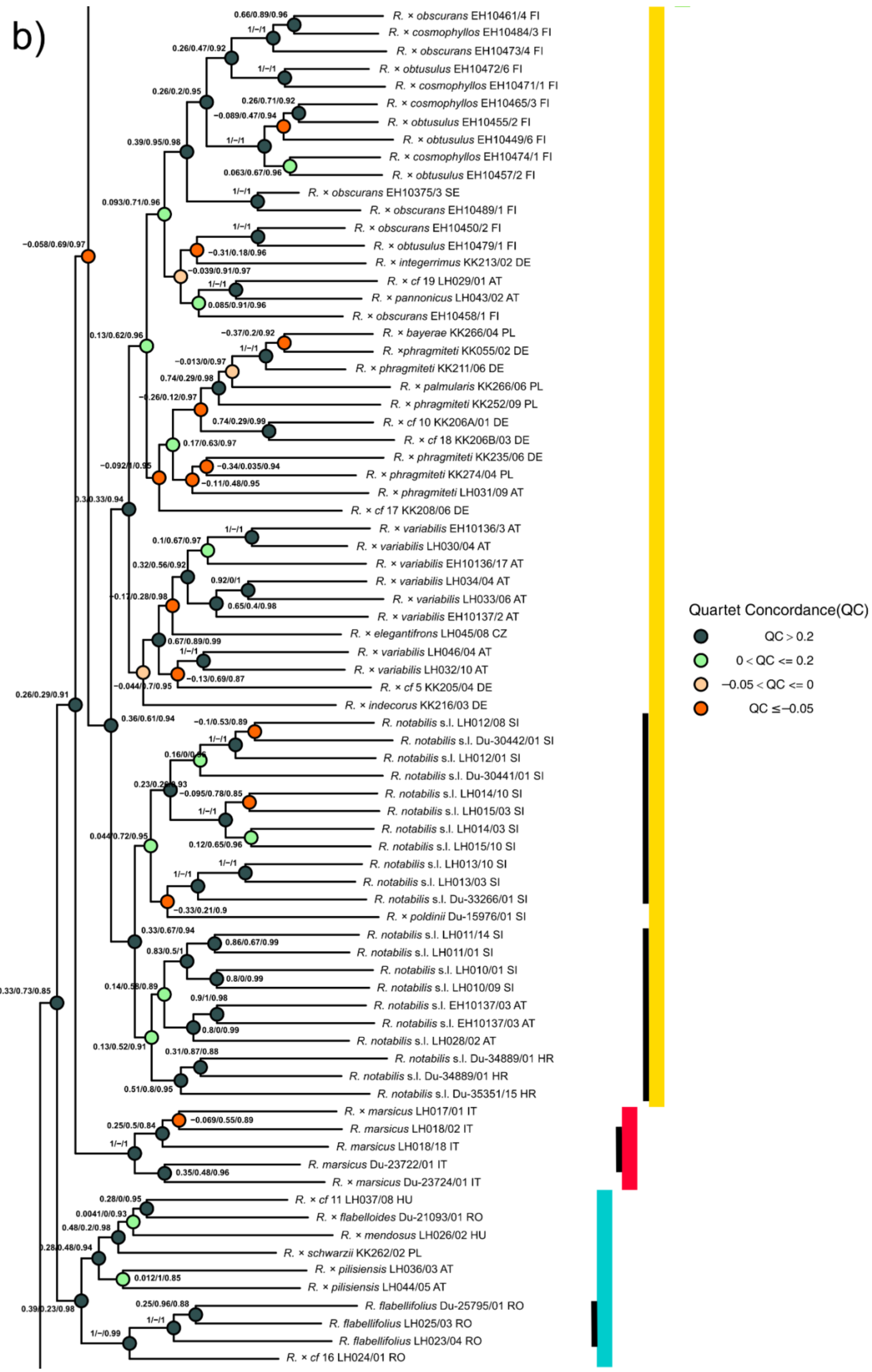
b)
R. × obscurans EH10461/4 FI
R. × cosmophyllos EH10484/3 FI
R. × obscurans EH10473/4 FI
R. × obtusulus EH10472/6 FI
R. × cosmophyllos EH10471/1 FI
R. × cosmophyllos EH10465/3 FI
R. × obtusulus EH10455/2 FI
R. × obtusulus EH10449/6 FI
R. × cosmophyllos EH10474/1 FI
R. × obtusulus EH10457/2 FI
R. × obscurans EH10375/3 SE
R. × obscurans EH10489/1 FI
R. × obscurans EH10450/2 FI
R. × obtusulus EH10479/1 FI
R. × integerrimus KK213/02 DE
R. × cf 19 LH029/01 AT
R. × pannonicus LH043/02 AT
R. × obscurans EH10458/1 FI
R. × bayerae KK266/04 PL
R. ×phragmiteti KK055/02 DE
R. × phragmiteti KK211/06 DE
R. × palmularis KK266/06 PL
R. × phragmiteti KK252/09 PL
R. × cf 10 KK206A/01 DE
R. × cf 18 KK206B/03 DE
R. × phragmiteti KK235/06 DE
R. × phragmiteti KK274/04 PL
R. × phragmiteti LH031/09 AT
R. × cf 17 KK208/06 DE
R. × variabilis EH10136/3 AT
R. × variabilis LH030/04 AT
R. × variabilis EH10136/17 AT
R. × variabilis LH034/04 AT
R. × variabilis LH033/06 AT
R. × variabilis EH10137/2 AT
R. × elegantifrons LH045/08 CZ
R. × variabilis LH046/04 AT
R. × variabilis LH032/10 AT
R. × cf 5 KK205/04 DE
R. × indecorus KK216/03 DE
R. notabilis s.l. LH012/08 SI
R. notabilis s.l. Du-30442/01 SI
R. notabilis s.l. LH012/01 SI
R. notabilis s.l. Du-30441/01 SI
R. notabilis s.l. LH014/10 SI
R. notabilis s.l. LH015/03 SI
R. notabilis s.l. LH014/03 SI
R. notabilis s.l. LH015/10 SI
R. notabilis s.l. LH013/10 SI
R. notabilis s.l. LH013/03 SI
R. notabilis s.l. Du-33266/01 SI
R. × poldinii Du-15976/01 SI
R. notabilis s.l. LH011/14 SI
R. notabilis s.l. LH011/01 SI
R. notabilis s.l. LH010/01 SI
R. notabilis s.l. LH010/09 SI
R. notabilis s.l. EH10137/03 AT
R. notabilis s.l. EH10137/03 AT
R. notabilis s.l. LH028/02 AT
R. notabilis s.l. Du-34889/01 HR
R. notabilis s.l. Du-34889/01 HR
R. notabilis s.l. Du-35351/15 HR
R. × marsicus LH017/01 IT
R. marsicus LH018/02 IT
R. marsicus LH018/18 IT
R. marsicus Du-23722/01 IT
R. × marsicus Du-23724/01 IT
R. × cf 11 LH037/08 HU
R. × flabelloides Du-21093/01 RO
R. × mendosus LH026/02 HU
R. × schwarzii KK262/02 PL
R. × pilisiensis LH036/03 AT
R. × pilisiensis LH044/05 AT
R. flabellifolius Du-25795/01 RO
R. flabellifolius LH025/03 RO
R. flabellifolius LH023/04 RO
R. × cf 16 LH024/01 RO
Quartet Concordance(QC)
QC > 0.2
0 < QC <= 0.2
−0.05 < QC <= 0
QC ≤ −0.05

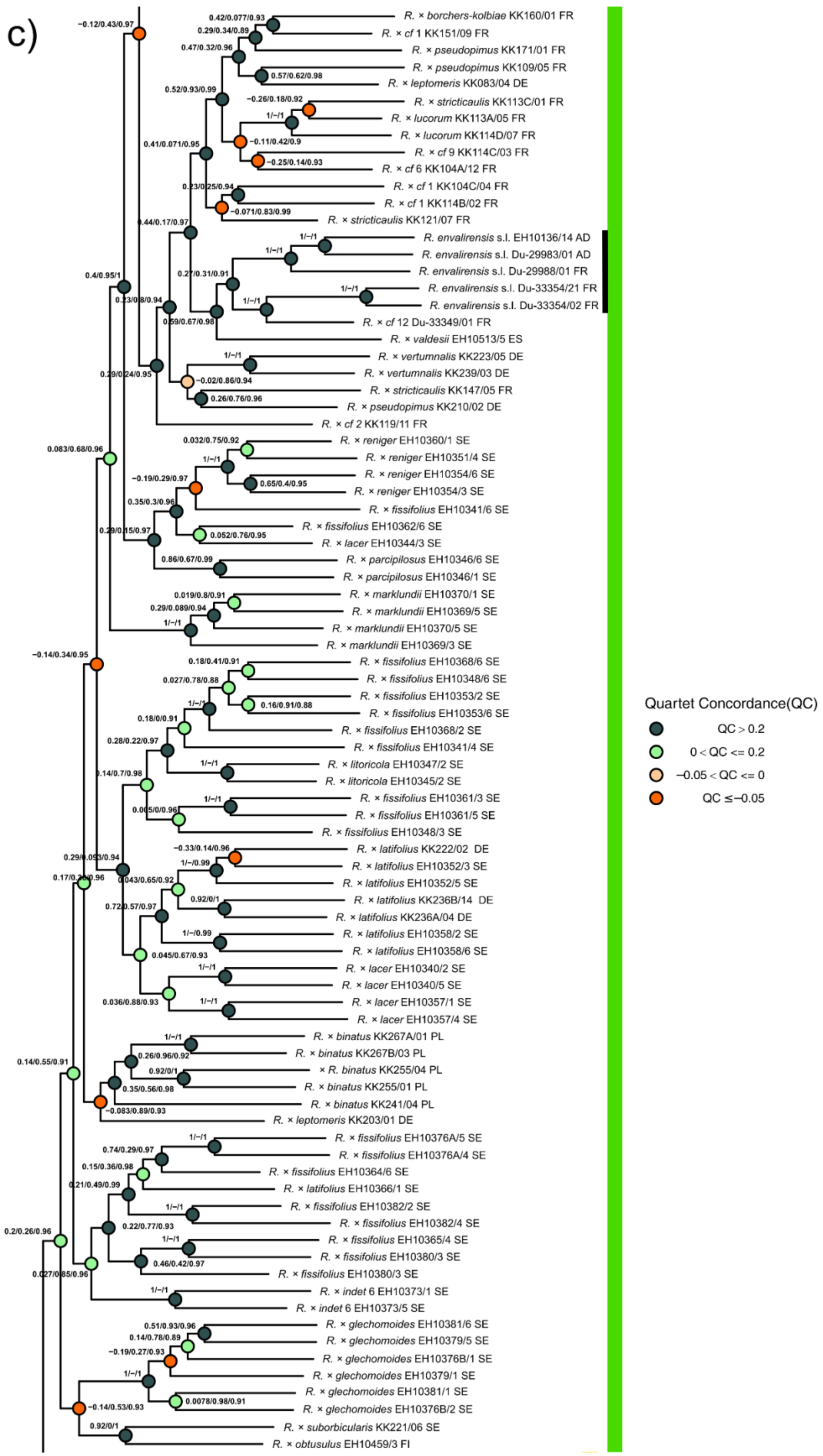
c)
R. × borchers-kolbiae KK160/01 FR
R. × cf 1 KK151/09 FR
R. × pseudopimus KK171/01 FR
R. × pseudopimus KK109/05 FR
R. × leptomeris KK083/04 DE
R. × stricticaulis KK113C/01 FR
R. × lucorum KK113A/05 FR
R. × lucorum KK113D/07 FR
R. × cf 9 KK114C/03 FR
R. × cf 6 KK104A/12 FR
R. × cf 1 KK104C/04 FR
R. × cf 1 KK114B/02 FR
R. × stricticaulis KK121/07 FR
R. envalirensis s.l. EH10136/14 AD
R. envalirensis s.l. Du-29983/01 AD
R. envalirensis s.l. Du-29988/01 FR
R. envalirensis s.l. Du-33354/21 FR
R. envalirensis s.l. Du-33354/02 FR
R. × cf 12 Du-33349/01 FR
R. × valdesii EH10513/5 ES
R. × vertumnalis KK223/05 DE
R. × vertumnalis KK239/03 DE
R. × stricticaulis KK147/05 FR
R. × pseudopimus KK210/02 DE
R. × cf 2 KK119/11 FR
R. × reniger EH10360/1 SE
R. × reniger EH10351/4 SE
R. × reniger EH10354/6 SE
R. × reniger EH10354/3 SE
R. × fissifolius EH10341/6 SE
R. × fissifolius EH10362/6 SE
R. × lacer EH10344/3 SE
R. × parcipilosus EH10346/6 SE
R. × parcipilosus EH10346/1 SE
R. × marklundii EH10370/1 SE
R. × marklundii EH10369/5 SE
R. × marklundii EH10370/5 SE
R. × marklundii EH10369/3 SE
R. × fissifolius EH10368/6 SE
R. × fissifolius EH10348/6 SE
R. × fissifolius EH10353/2 SE
R. × fissifolius EH10353/6 SE
R. × fissifolius EH10368/2 SE
R. × fissifolius EH10341/4 SE
R. × litoricola EH10347/2 SE
R. × litoricola EH10345/2 SE
R. × fissifolius EH10361/3 SE
R. × fissifolius EH10361/5 SE
R. × fissifolius EH10348/3 SE
R. × latifolius KK222/02 DE
R. × latifolius EH10352/3 SE
R. × latifolius EH10352/5 SE
R. × latifolius KK236B/14 DE
R. × latifolius KK236A/04 DE
R. × latifolius EH10358/2 SE
R. × latifolius EH10358/6 SE
R. × lacer EH10340/2 SE
R. × lacer EH10340/5 SE
R. × lacer EH10357/1 SE
R. × lacer EH10357/4 SE
R. × binatus KK267A/01 PL
R. × binatus KK267B/03 PL
× R. binatus KK255/04 PL
R. × binatus KK255/01 PL
R. × binatus KK241/04 PL
R. × leptomeris KK203/01 DE
R. × fissifolius EH10376A/5 SE
R. × fissifolius EH10376A/4 SE
R. × fissifolius EH10364/6 SE
R. × latifolius EH10366/1 SE
R. × fissifolius EH10382/2 SE
R. × fissifolius EH10382/4 SE
R. × fissifolius EH10365/4 SE
R. × fissifolius EH10380/3 SE
R. × fissifolius EH10380/3 SE
R. × indet 6 EH10373/1 SE
R. × indet 6 EH10373/5 SE
R. × glechomoides EH10381/6 SE
R. × glechomoides EH10379/5 SE
R. × glechomoides EH10376B/1 SE
R. × glechomoides EH10379/1 SE
R. × glechomoides EH10381/1 SE
R. × glechomoides EH10376B/2 SE
R. × suborbicularis KK221/06 SE
R. × obtusulus EH10459/3 FI
Quartet Concordance(QC)
QC > 0.2
0 < QC <= 0.2
−0.05 < QC <= 0
QC ≤ −0.05

d)

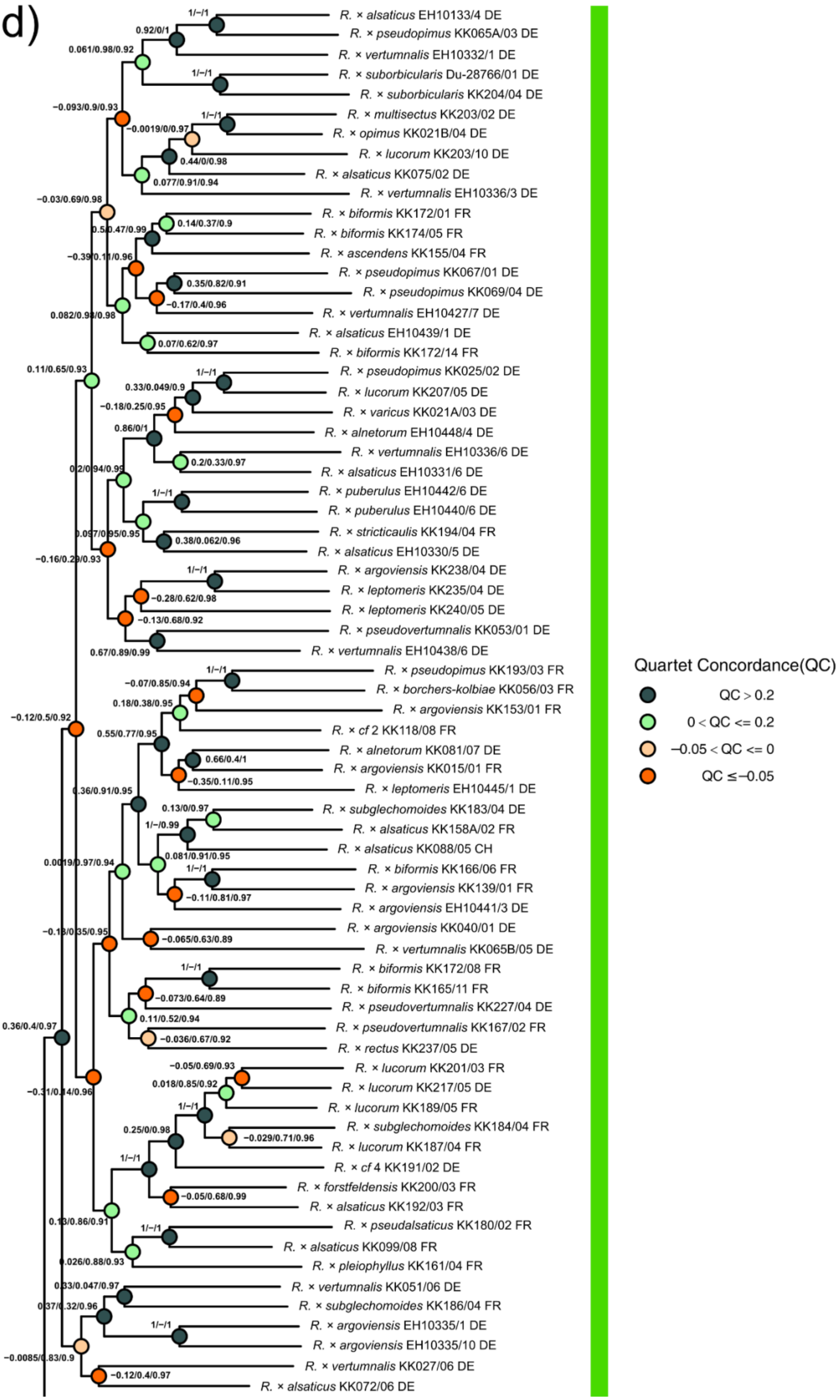

R. × alsaticus EH10133/4 DE
R. × pseudopimus KK065A/03 DE
R. × vertumnalis EH10332/1 DE
R. × suborbicularis Du-28766/01 DE
R. × suborbicularis KK204/04 DE
R. × multisectus KK203/02 DE
R. × opimus KK021B/04 DE
R. × lucorum KK203/10 DE
R. × alsaticus KK075/02 DE
R. × vertumnalis EH10336/3 DE
R. × biformis KK172/01 FR
R. × biformis KK174/05 FR
R. × ascendens KK155/04 FR
R. × pseudopimus KK067/01 DE
R. × pseudopimus KK069/04 DE
R. × vertumnalis EH10427/7 DE
R. × alsaticus EH10439/1 DE
R. × biformis KK172/14 FR
R. × pseudopimus KK025/02 DE
R. × lucorum KK207/05 DE
R. × varicus KK021A/03 DE
R. × alnetorum EH10448/4 DE
R. × vertumnalis EH10336/6 DE
R. × alsaticus EH10331/6 DE
R. × puberulus EH10442/6 DE
R. × puberulus EH10440/6 DE
R. × stricticaulis KK194/04 FR
R. × alsaticus EH10330/5 DE
R. × argoviensis KK238/04 DE
R. × leptomeris KK235/04 DE
R. × leptomeris KK240/05 DE
R. × pseudovertumnalis KK053/01 DE
R. × vertumnalis EH10438/6 DE
R. × pseudopimus KK193/03 FR
R. × borchers-kolbiae KK056/03 FR
R. × argoviensis KK153/01 FR
R. × cf 2 KK118/08 FR
R. × alnetorum KK081/07 DE
R. × argoviensis KK015/01 FR
R. × leptomeris EH10445/1 DE
R. × subglechomoides KK183/04 DE
R. × alsaticus KK158A/02 FR
R. × alsaticus KK088/05 CH
R. × biformis KK166/06 FR
R. × argoviensis KK139/01 FR
R. × argoviensis EH10441/3 DE
R. × argoviensis KK040/01 DE
R. × vertumnalis KK065B/05 DE
R. × biformis KK172/08 FR
R. × biformis KK165/11 FR
R. × pseudovertumnalis KK227/04 DE
R. × pseudovertumnalis KK167/02 FR
R. × rectus KK237/05 DE
R. × lucorum KK201/03 FR
R. × lucorum KK217/05 DE
R. × lucorum KK189/05 FR
R. × subglechomoides KK184/04 FR
R. × lucorum KK187/04 FR
R. × cf 4 KK191/02 DE
R. × forstfeldensis KK200/03 FR
R. × alsaticus KK192/03 FR
R. × pseudalsaticus KK180/02 FR
R. × alsaticus KK099/08 FR
R. × pleiophyllus KK161/04 FR
R. × vertumnalis KK051/06 DE
R. × subglechomoides KK186/04 FR
R. × argoviensis EH10335/1 DE
R. × argoviensis EH10335/10 DE
R. × vertumnalis KK027/06 DE
R. × alsaticus KK072/06 DE
Quartet Concordance(QC)
QC > 0.2
0 < QC <= 0.2
−0.05 < QC <= 0
QC ≤ −0.05

**Supplementary Fig. S3.** Concatenated ML tree with quartet sampling scores (QS). (a-d) Results are based on RAxML-NG results and the ‘min10’ RAD-Seq alignment (a-d, see also Fig. 4; 280 samples, 97,312 loci, 435,678 SNPs). The quartet concordance score (QC) is the ratio of concordant to both discordant quartets (1: all concordant, > 0: more concordant patterns, < 0: more discordant patterns), the quartet differential score (QD) describes the skewness of both discordant patterns (1: equal, 0.3: skewed, 0: all topologies 1 or 2), and the quartet informativeness score (QI) shows the proportion of informative replicates (1: all informative, 0: none informative; see also Pease et al. 2018). Incomplete lineage sorting (ILS) is indicated by QD values around 1 and means the presence of both discordant topologies (random pattern). QD values towards 0 hint at directional introgression, i.e., the presence of only one particular alternative topology (Pease et al. 2018; Karbstein et al. 2020). We used the R script plot_QC_ggtree.R (available at: https://github.com/-ShuiyinLIU/QS_visualization) and the R packages GGTREE vers. 2.2.4 (Yu et al. 2017), Treeio vers 1.12.0 (Wang et al. 2020), GGPLOT2 vers. 3.3.1 (Winston et al. 2020), and APE vers. 5.4 (Paradis and Schliep 2019; Paradis et al. 2020) for visualization. Colored bars highlight main clades of Fig. 4a. Black bars indicate sexual species. Names of polyploid apomictic taxa are tentative. Squared brackets: A part of the branch was cut for illustratative purposes.

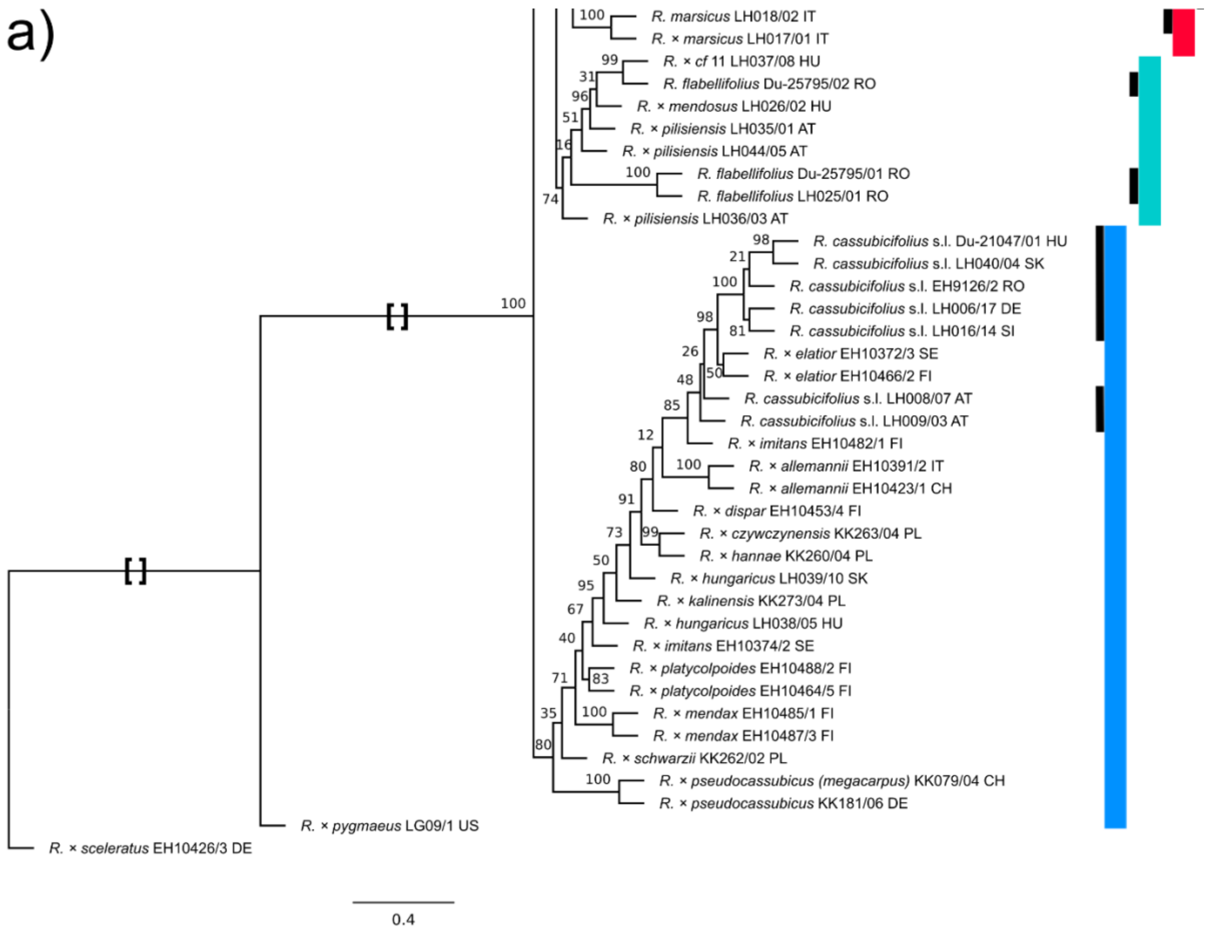

a)
R. marsicus LH018/02 IT
R. × marsicus LH017/01 IT
R. × cf 11 LH037/08 HU
R. flabellifolius Du-25795/02 RO
R. × mendosus LH026/02 HU
R. × pilisiensis LH035/01 AT
R. × pilisiensis LH044/05 AT
R. flabellifolius Du-25795/01 RO
R. flabellifolius LH025/01 RO
R. × pilisiensis LH036/03 AT
R. cassubicifolius s.l. Du-21047/01 HU
R. cassubicifolius s.l. LH040/04 SK
R. cassubicifolius s.l. EH9126/2 RO
R. cassubicifolius s.l. LH006/17 DE
R. cassubicifolius s.l. LH016/14 SI
R. × elatior EH10372/3 SE
R. × elatior EH10466/2 FI
R. cassubicifolius s.l. LH008/07 AT
R. cassubicifolius s.l. LH009/03 AT
R. × imitans EH10482/1 FI
R. × allemannii EH10391/2 IT
R. × allemannii EH10423/1 CH
R. × dispar EH10453/4 FI
R. × czywczynensis KK263/04 PL
R. × hannae KK260/04 PL
R. × hungaricus LH039/10 SK
R. × kalinensis KK273/04 PL
R. × hungaricus LH038/05 HU
R. × imitans EH10374/2 SE
R. × platycolpoides EH10488/2 FI
R. × platycolpoides EH10464/5 FI
R. × mendax EH10485/1 FI
R. × mendax EH10487/3 FI
R. × schwarzii KK262/02 PL
R. × pseudocassubicus (megacarpus) KK079/04 CH
R. × pseudocassubicus KK181/06 DE
R. × pygmaeus LG09/1 US
R. × sceleratus EH10426/3 DE
0.4

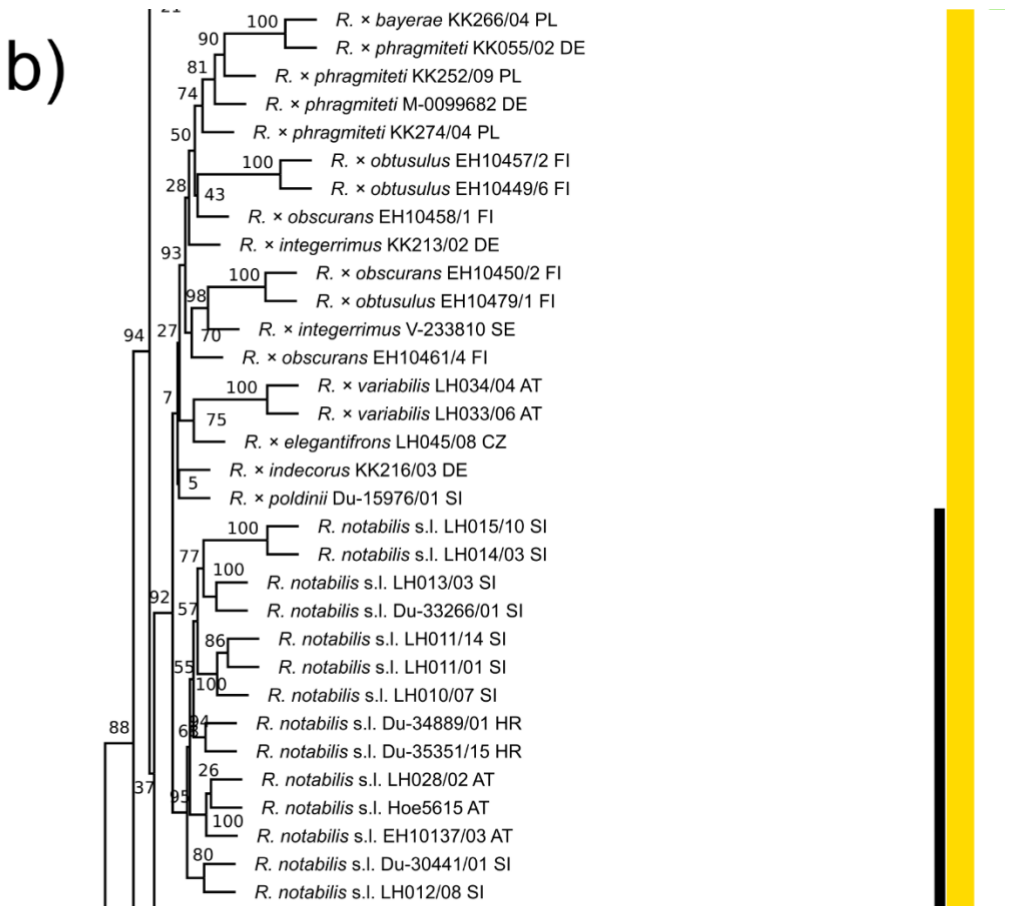
b)
R. × bayerae KK266/04 PL
R. × phragmiteti KK055/02 DE
R. × phragmiteti KK252/09 PL
R. × phragmiteti M-0099682 DE
R. × phragmiteti KK274/04 PL
R. × obtusulus EH10457/2 FI
R. × obtusulus EH10449/6 FI
R. × obscurans EH10458/1 FI
R. × integerrimus KK213/02 DE
R. × obscurans EH10450/2 FI
R. × obtusulus EH10479/1 FI
R. × integerrimus V-233810 SE
R. × obscurans EH10461/4 FI
R. × variabilis LH034/04 AT
R. × variabilis LH033/06 AT
R. × elegantifrons LH045/08 CZ
R. × indecorus KK216/03 DE
R. × poldinii Du-15976/01 SI
R. notabilis s.l. LH015/10 SI
R. notabilis s.l. LH014/03 SI
R. notabilis s.l. LH013/03 SI
R. notabilis s.l. Du-33266/01 SI
R. notabilis s.l. LH011/14 SI
R. notabilis s.l. LH011/01 SI
R. notabilis s.l. LH010/07 SI
R. notabilis s.l. Du-34889/01 HR
R. notabilis s.l. Du-35351/15 HR
R. notabilis s.l. LH028/02 AT
R. notabilis s.l. Hoe5615 AT
R. notabilis s.l. EH10137/03 AT
R. notabilis s.l. Du-30441/01 SI
R. notabilis s.l. LH012/08 SI

c)

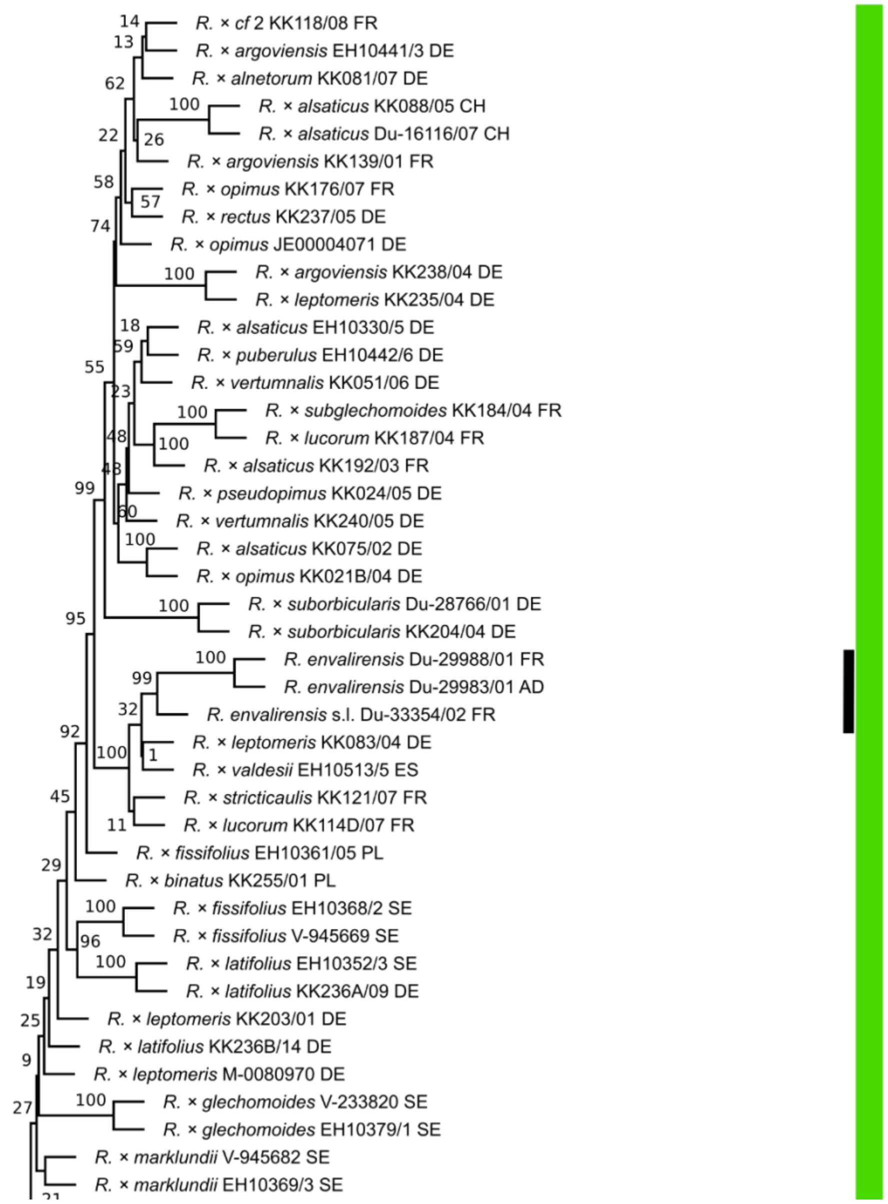


**Supplementary Fig. S4.** Coalescent-based tree ASTRAL tree. (a-c) Results are based on a TE alignment of 113 samples and 576 non-phased genes. Felsenstein Bootstrap Proportions (FBP) values are displayed on branches. Black bars indicate sexual species. Colored bars highlight main clades of Fig. 4b. Names of apomictic polyploid taxa are tentative. Squared brackets: A part of the branch was cut for illustratative purposes.

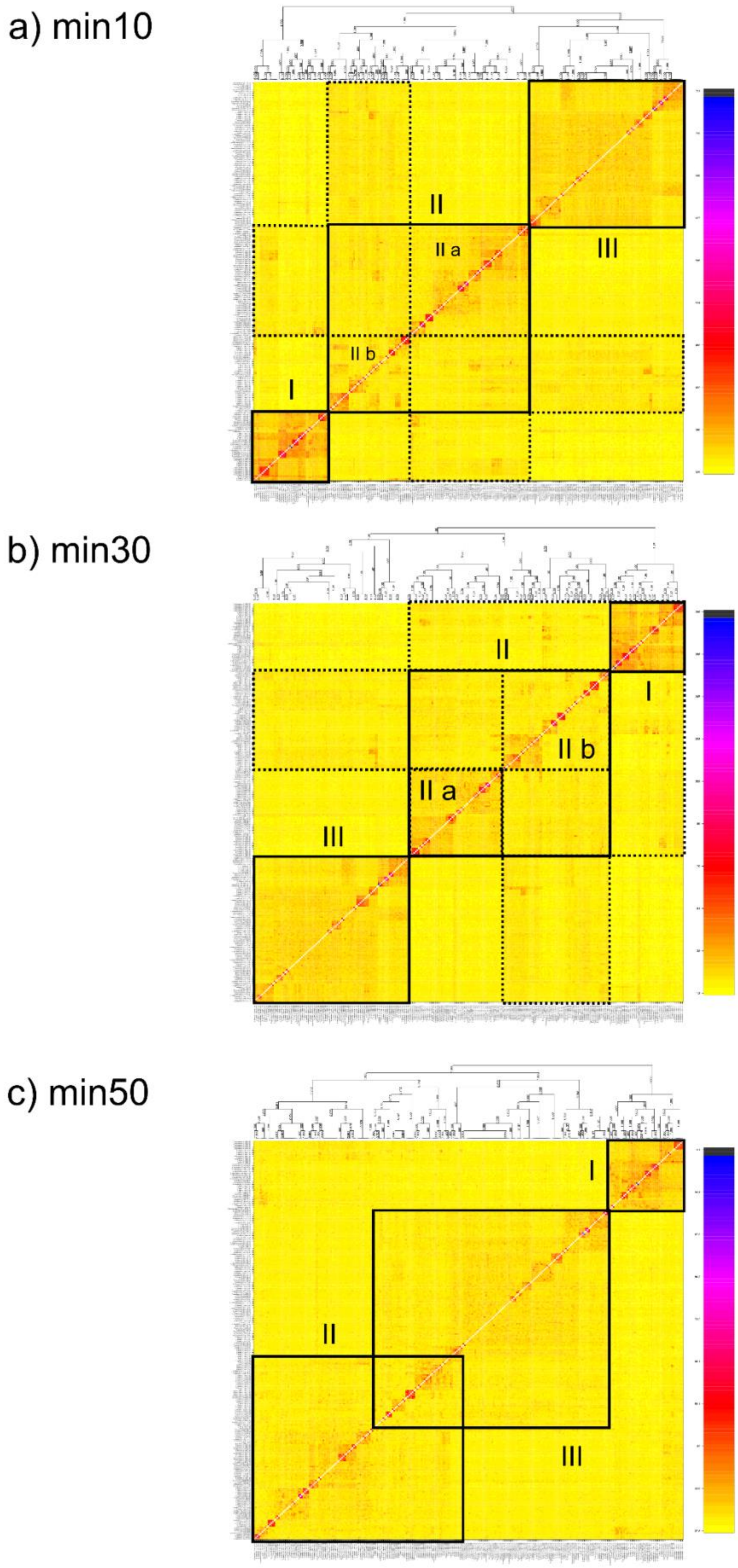

a) min10
II
II a
III
II b
I
b) min30
II
I
II b
II a
III
c) min50
I
II
III

**Supplementary Fig. S5.** Clustered fineRADstructure coancestry matrices. Results are based on 280 sexual and polyploid apomictic individuals of the *R. auricomus* complex and (a) the 'min10' (97,312 loci, 438,775 SNPs), (b) the 'min30' (33,165 loci, 194,083 SNPs), or (c) the 'min50' (11,196 loci, 64,554 SNPs) RAD-Seq alignments. The legend (on the right) shows the color-coding of pairwise genetic similarity: the darker the color, the higher the similarity between a pair of individuals. We highlighted supported genetic clusters with solid lines, shared similarity among (sub)clusters with dotted lines, and sexual species with broad dashed lines (subgenome C, F, M, N, and E). Above the coancestry matrix, the clustering structure is given (posterior probability group assignment probabilities). See Fig. 4 and Figshare data repository for different datasets and more detailed results.

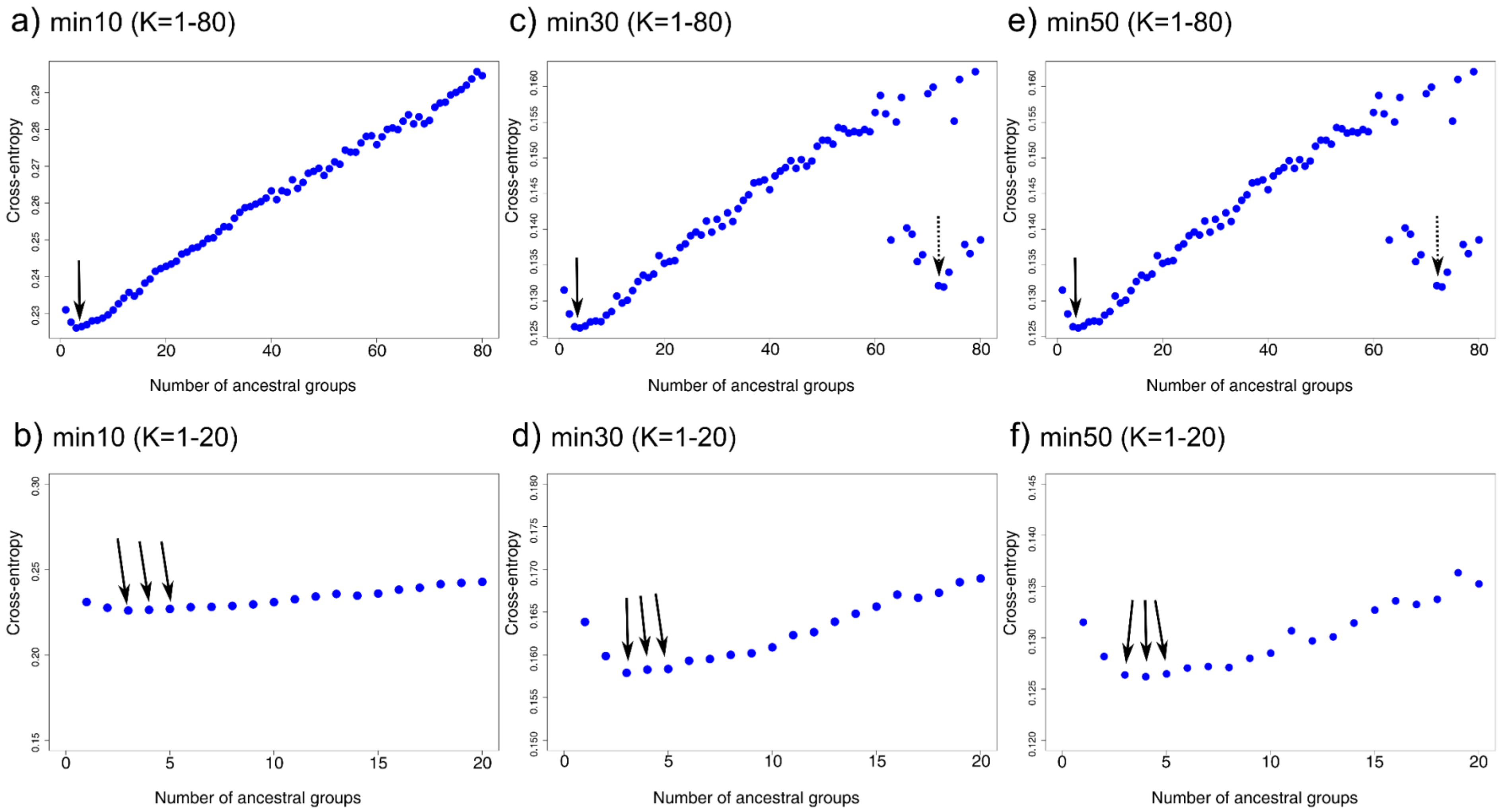


**Supplementary Fig. S6.** Cross-entropies based on sNMF results. Results are based on different unlinked-SNP-RAD-Seq datasets of K=80 assumed ancestral genetic groups incl. smaller sections (K=20), (a, b) 'min10' (97,312 loci), (c, d) 'min30' (33,165 loci), and (e, f) 'min50'

(11,196 loci). The cross-entropy criterion is based on the prediction of masked genotypes to evaluate the fit of a model with K populations (the lower, the better; Frichot et al. 2014). Solid arrows indicate global cross-entropy minima (K=3-5 across datasets), and dashed arrows show local minima.

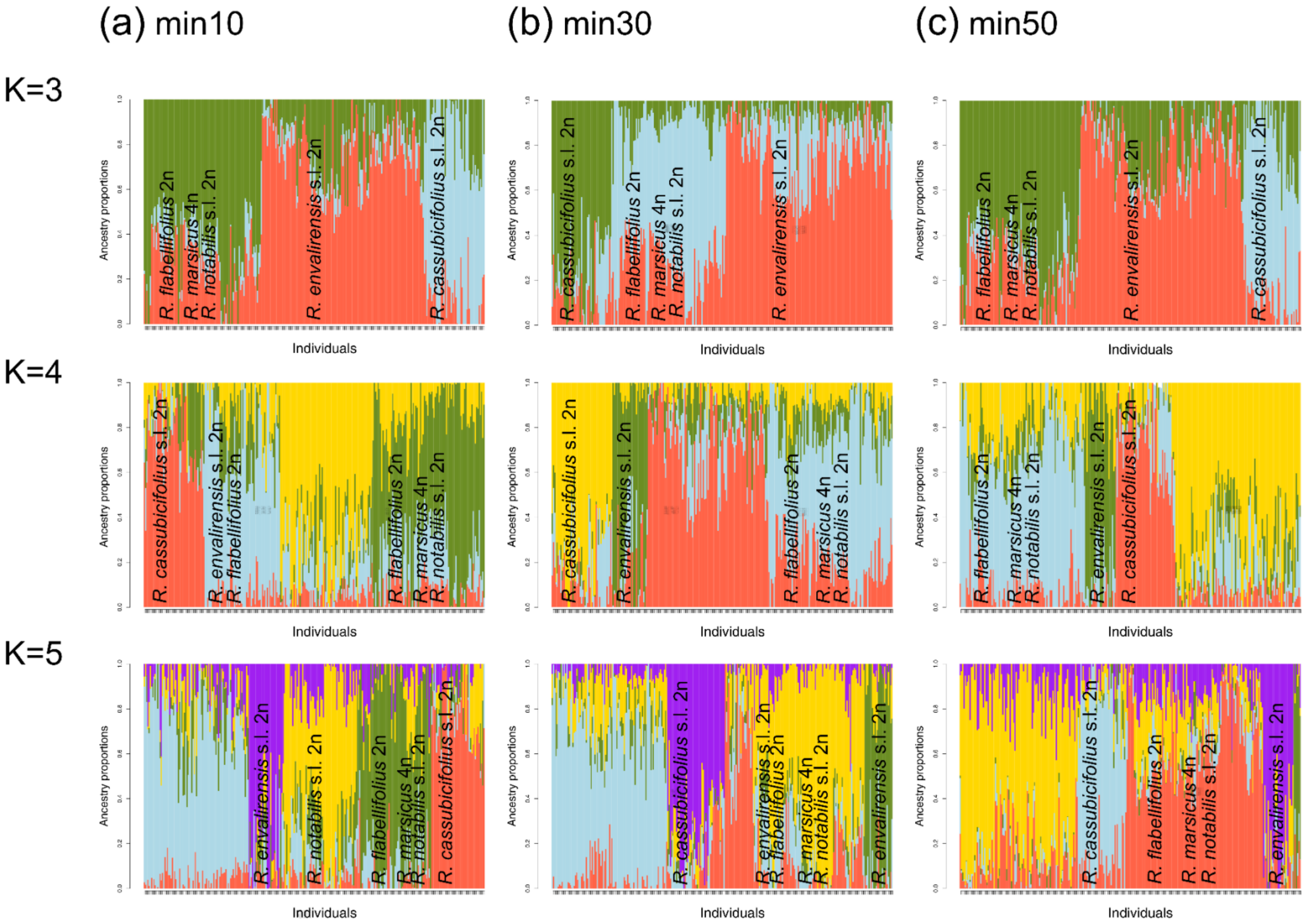

(a) min10
(b) min30
(c) min50
K=3
K=4
K=5
Ancestry proportions
Individuals
R. flabellifolius 2n
R. marsicus 4n
R. notabilis s.l. 2n
R. envalirensis s.l. 2n
R. cassubicifolius s.l. 2n

**Supplementary Fig. S7.** Ancestry proportions based on sNMF analyses. Results are based on 280 sexual and apomictic *R. auricomus* individuals, and different unlinked-SNP RAD-Seq datasets and numbers of ancestral genetic groups (K=3-5), i.e. (a) 'min10' (97,312 loci), (b) 'min30' (33,165 loci), and (c) 'min50' (11,196 loci). See Figshare data repository for detailed results (including sample names).

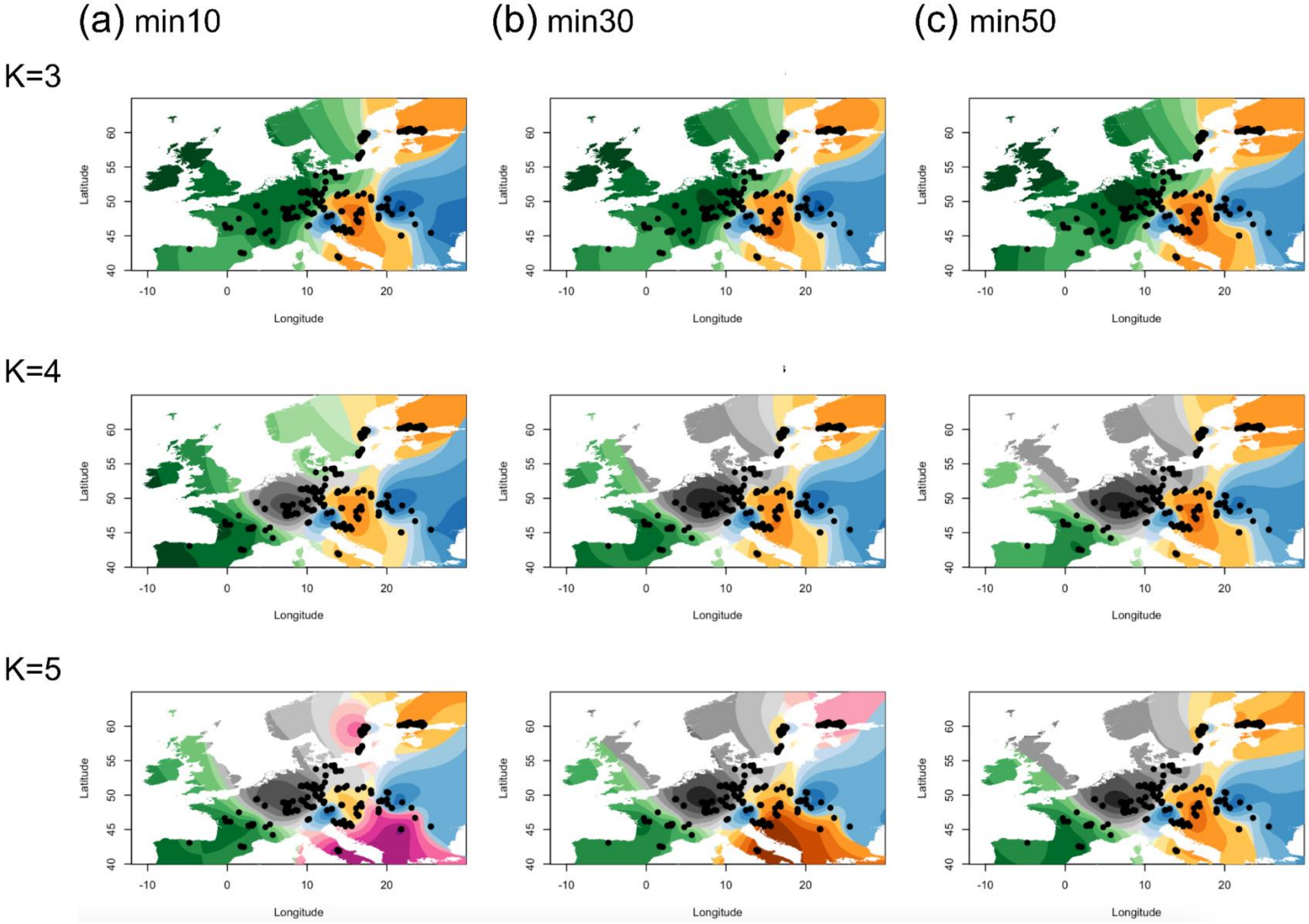
(a) min10
(b) min30
(c) min50
K=3
K=4
K=5
Latitude
Longitude

**Supplementary Fig. S8.** Geographic maps showing ancestry coefficients across Europe (K=3-5). Ancestry coefficients were illustrated with method ‘max’, i.e., at each point the cluster for which the ancestry coefficient is maximal. Results are based on sNMF analyses of 280 sexual and apomictic *R. auricomus* individuals, and different unlinked-SNP RAD-Seq datasets and numbers of ancestral genetic groups (K=3-5), (a) min10 (97,312 loci), (b) min30 (33,165 loci), and (c) min50 (11,196 loci). We colored maps according to Fig. 4 (except grey=Central European apomicts related to main clade E, pink=either (a) *R. flabellifolius*, *R. marsicus*, and Southern Scandinavian apomicts, (b) a separate apomictic cluster containing many Finnish taxa (related to the orange one (subgenome N, most reasonable according to other results), or (c) a separate apomictic cluster containing a few Finnish taxa). Results were similar among K=3 dataset. K=4 datasets slightly differed from each other (e.g., min10 unlikely because of disjunct grouping the subgenome ‘E’ group). K=5 datasets remarkably differed from each other (e.g., min10 unlikely because of disjunct distribution of *R. flabellifolius, R. marsicus*, and Southern Scandinavian apomicts or min50 unlikely because of a separate apomictic cluster containing a few Finnish taxa). Europe map source: https://maps.ngdc.noaa.gov.

a)

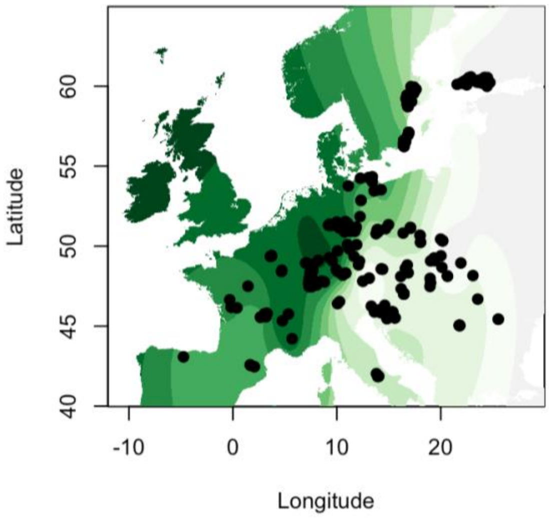

Latitude
40
45
50
55
60
-10
0
10
20
Longitude


b)

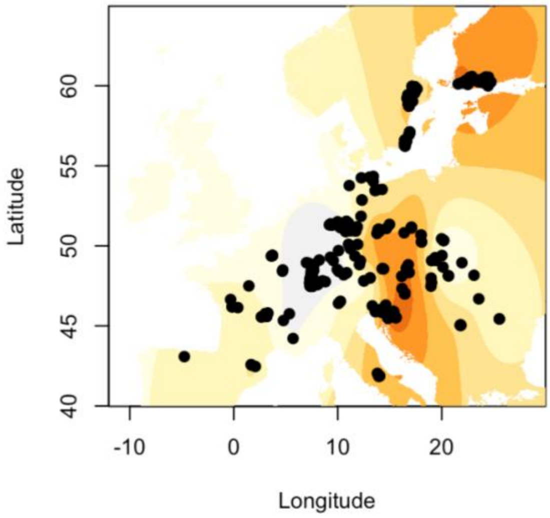

Latitude
40
45
50
55
60
-10
0
10
20
Longitude


c)

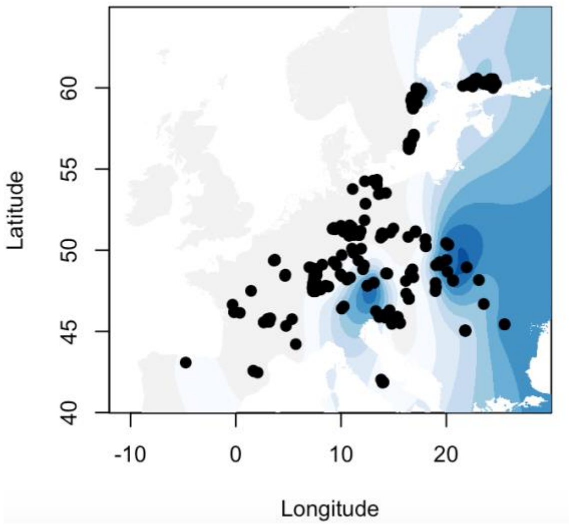


**Supplementary Fig. S9.** Geographic maps showing separate ancestry coefficients across Europe (K=3). Ancestry coefficients are illustrated. Results are based on sNMF analyses of 280 sexual and apomictic *R. auricomus* individuals and the ‘min30’ unlinked-SNP RAD-Seq dataset (33,165 loci) for K=3 genetic clusters (a-c; see Fig. 7, Supplementary Figs. S12, S13 and Results for details). Europe map source: https://maps.ngdc.noaa.gov.

a)

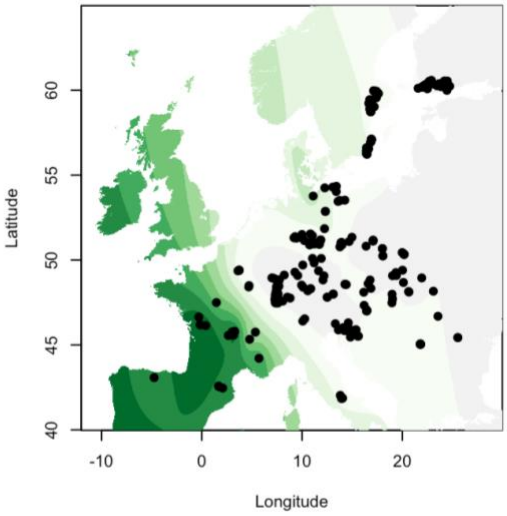

Latitude
60
55
50
45
40
-10
0
10
20
Longitude


b)

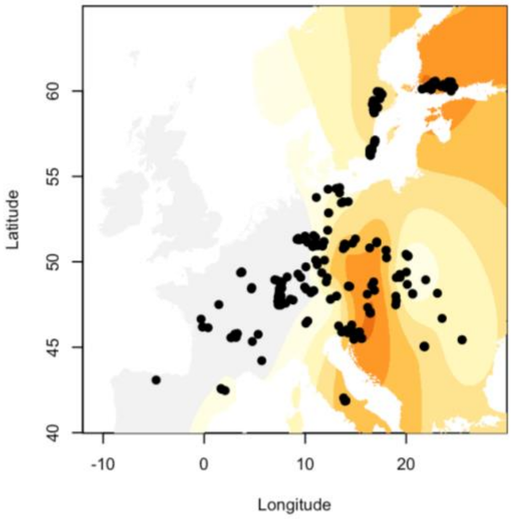

Latitude
60
55
50
45
40
-10
0
10
20
Longitude


c)

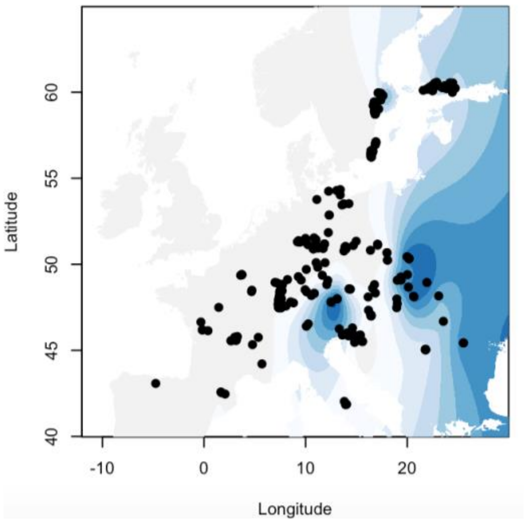


d)

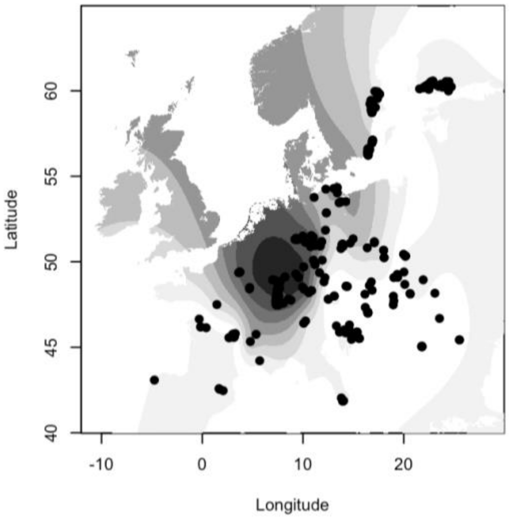


**Supplementary Fig. S10.** Geographic maps showing separate ancestry coefficients across Europe (K=4). Ancestry coefficients are illustrated. Results are based on sNMF analyses of 280 sexual and apomictic *R. auricomus* individuals and the ‘min30’ unlinked-SNP RAD-Seq dataset (33,165 loci) for K=4 genetic clusters (a-d; see Fig. 7, Supplementary Figs. S12, S13 and Results for details). Europe map source: https://maps.ngdc.noaa.gov.

a)

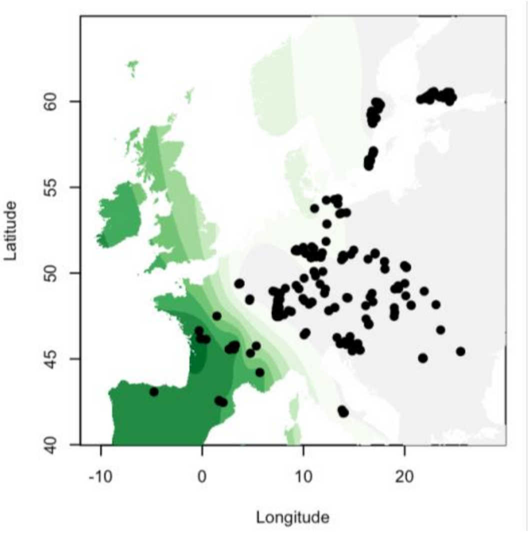
Latitude
40
45
50
55
60
-10
0
10
20
Longitude

b)

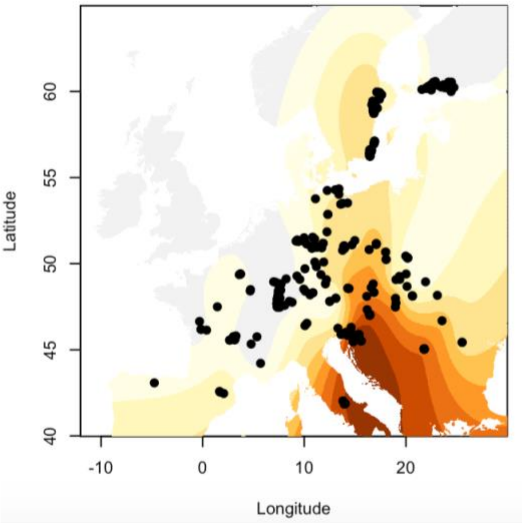
Latitude
40
45
50
55
60
-10
0
10
20
Longitude

c)

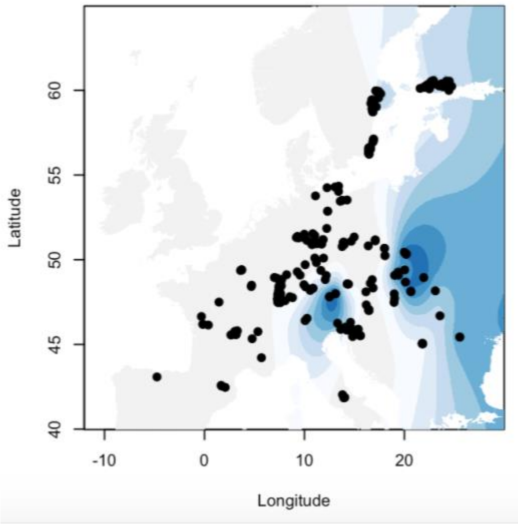
Latitude
Longitude
40
45
50
55
60
-10
0
10
20

d)

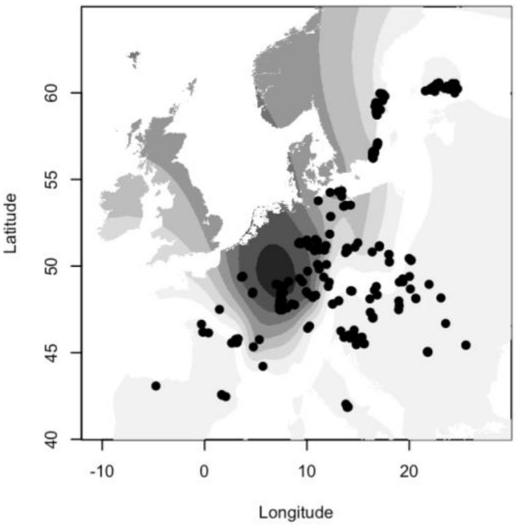
Latitude
Longitude
40
45
50
55
60
-10
0
10
20

e)

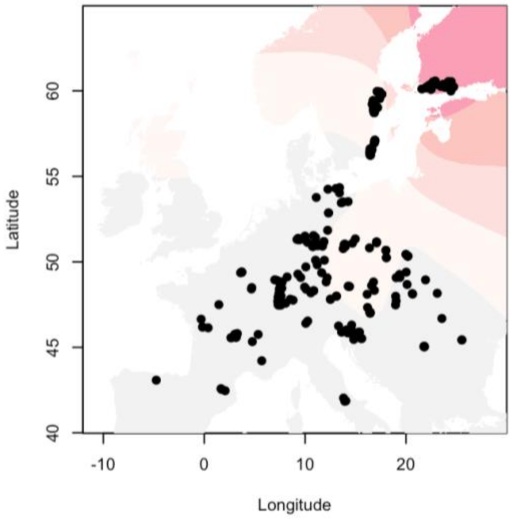


**Supplementary Fig. S11.** Geographic maps showing separate ancestry coefficients across Europe (K=5). Ancestry coefficients are illustrated. Results are based on sNMF analyses of 280 sexual and apomictic *R. auricomus* individuals and the 'min30' unlinked-SNP RAD-Seq dataset (33,165 loci) for K=5 genetic clusters (a-e; see Fig. 7, Supplementary Figs. S12, S13 and Results for details). Europe map source: https://maps.ngdc.noaa.gov.

a)

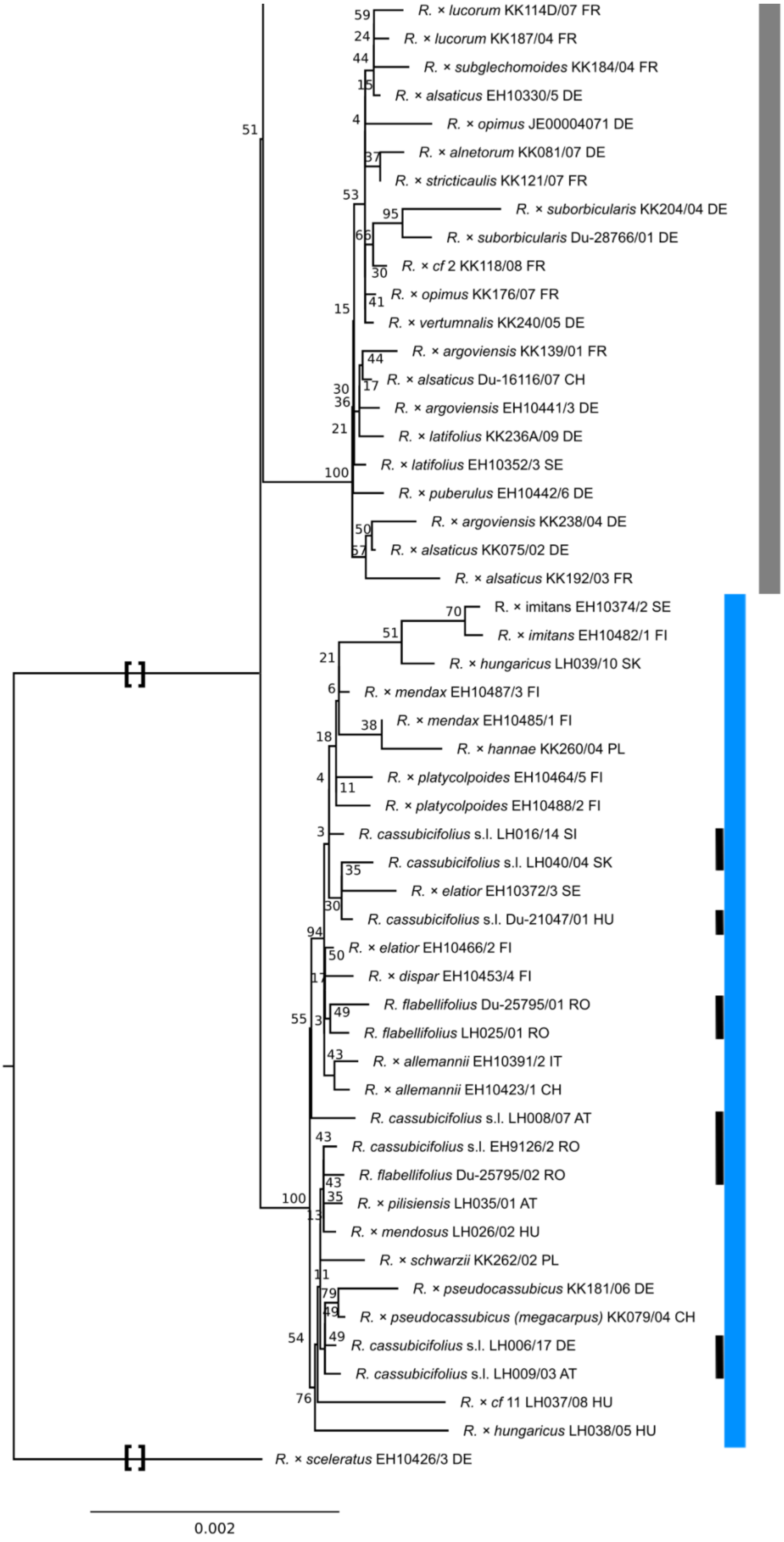
R. × lucorum KK114D/07 FR
R. × lucorum KK187/04 FR
R. × subglechomoides KK184/04 FR
R. × alsaticus EH10330/5 DE
R. × opimus JE00004071 DE
R. × alnetorum KK081/07 DE
R. × stricticaulis KK121/07 FR
R. × suborbicularis KK204/04 DE
R. × suborbicularis Du-28766/01 DE
R. × cf 2 KK118/08 FR
R. × opimus KK176/07 FR
R. × vertumnalis KK240/05 DE
R. × argoviensis KK139/01 FR
R. × alsaticus Du-16116/07 CH
R. × argoviensis EH10441/3 DE
R. × latifolius KK236A/09 DE
R. × latifolius EH10352/3 SE
R. × puberulus EH10442/6 DE
R. × argoviensis KK238/04 DE
R. × alsaticus KK075/02 DE
R. × alsaticus KK192/03 FR
R. × imitans EH10374/2 SE
R. × imitans EH10482/1 FI
R. × hungaricus LH039/10 SK
R. × mendax EH10487/3 FI
R. × mendax EH10485/1 FI
R. × hannae KK260/04 PL
R. × platycolpoides EH10464/5 FI
R. × platycolpoides EH10488/2 FI
R. cassubicifolius s.l. LH016/14 SI
R. cassubicifolius s.l. LH040/04 SK
R. × elatior EH10372/3 SE
R. cassubicifolius s.l. Du-21047/01 HU
R. × elatior EH10466/2 FI
R. × dispar EH10453/4 FI
R. flabellifolius Du-25795/01 RO
R. flabellifolius LH025/01 RO
R. × allemannii EH10391/2 IT
R. × allemannii EH10423/1 CH
R. cassubicifolius s.l. LH008/07 AT
R. cassubicifolius s.l. EH9126/2 RO
R. flabellifolius Du-25795/02 RO
R. × pilisiensis LH035/01 AT
R. × mendosus LH026/02 HU
R. × schwarzii KK262/02 PL
R. × pseudocassubicus KK181/06 DE
R. × pseudocassubicus (megacarpus) KK079/04 CH
R. cassubicifolius s.l. LH006/17 DE
R. cassubicifolius s.l. LH009/03 AT
R. × cf 11 LH037/08 HU
R. × hungaricus LH038/05 HU
R. × sceleratus EH10426/3 DE
0.002

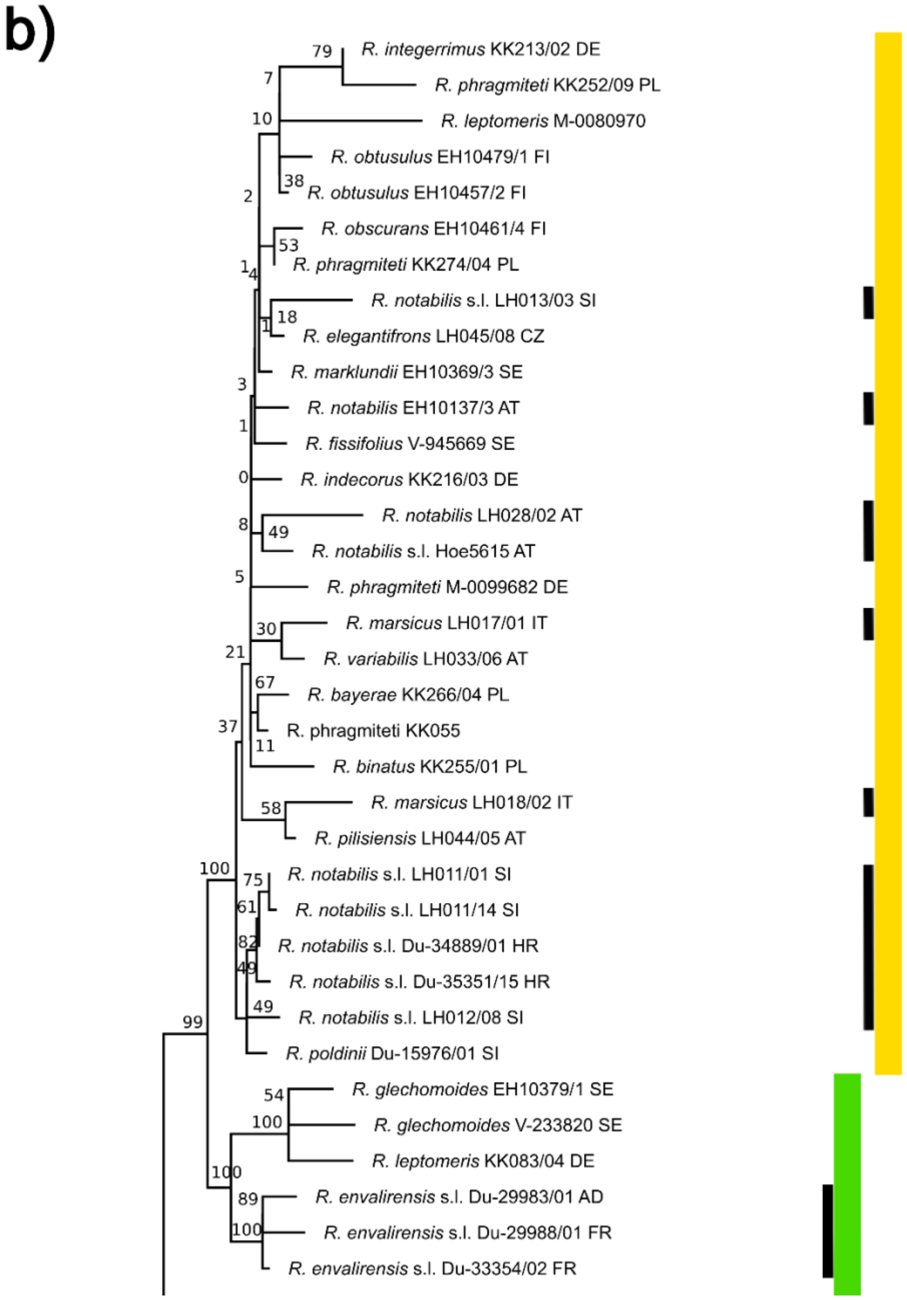


**Supplementary Fig. S12.** Concatenated ML tree inferred from plastid regions (CP). Results are based on a CP alignment of 87 samples and 71 plastid regions. Per branch, Felsenstein Bootstrap Proportions (FBP) values are displayed. Black bars indicate sexual species. Colored bars highlight main clades of Fig. 5a. Names of apomictic polyploid taxa are tentative. Squared brackets: A part of the branch was cut for illustrative purposes.

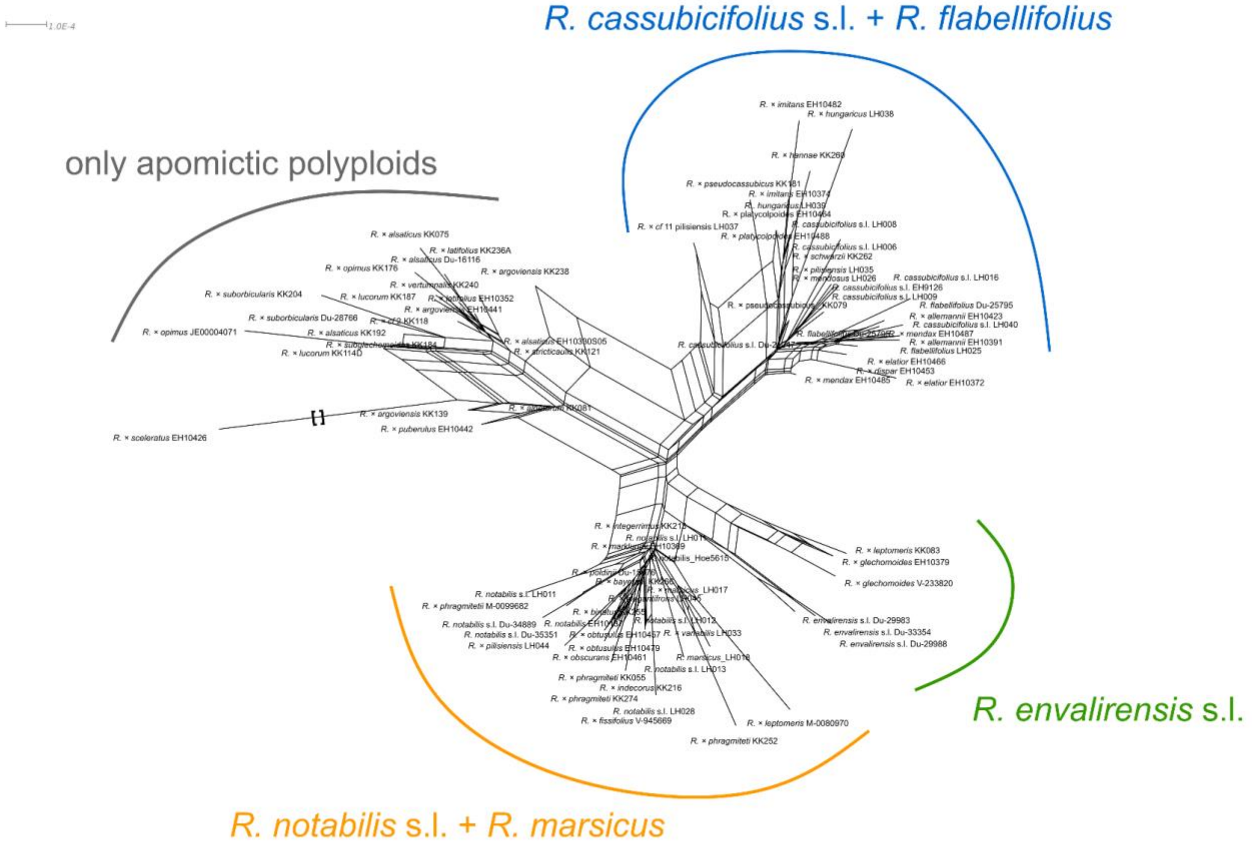


**Supplementary Fig. S13.** Neighbor-net analysis (SplitsTree) of CP data. Results are based on 87 samples and 71 plastid regions of the plastome dataset and genetic distances (general time reversible [GTR] model with estimated site frequencies and ML). We colored main splits according to Fig. 5a. No BT values are given for reasons of clarity and comprehensibility. Squared brackets: A part of the branch was cut for illustrative purposes.

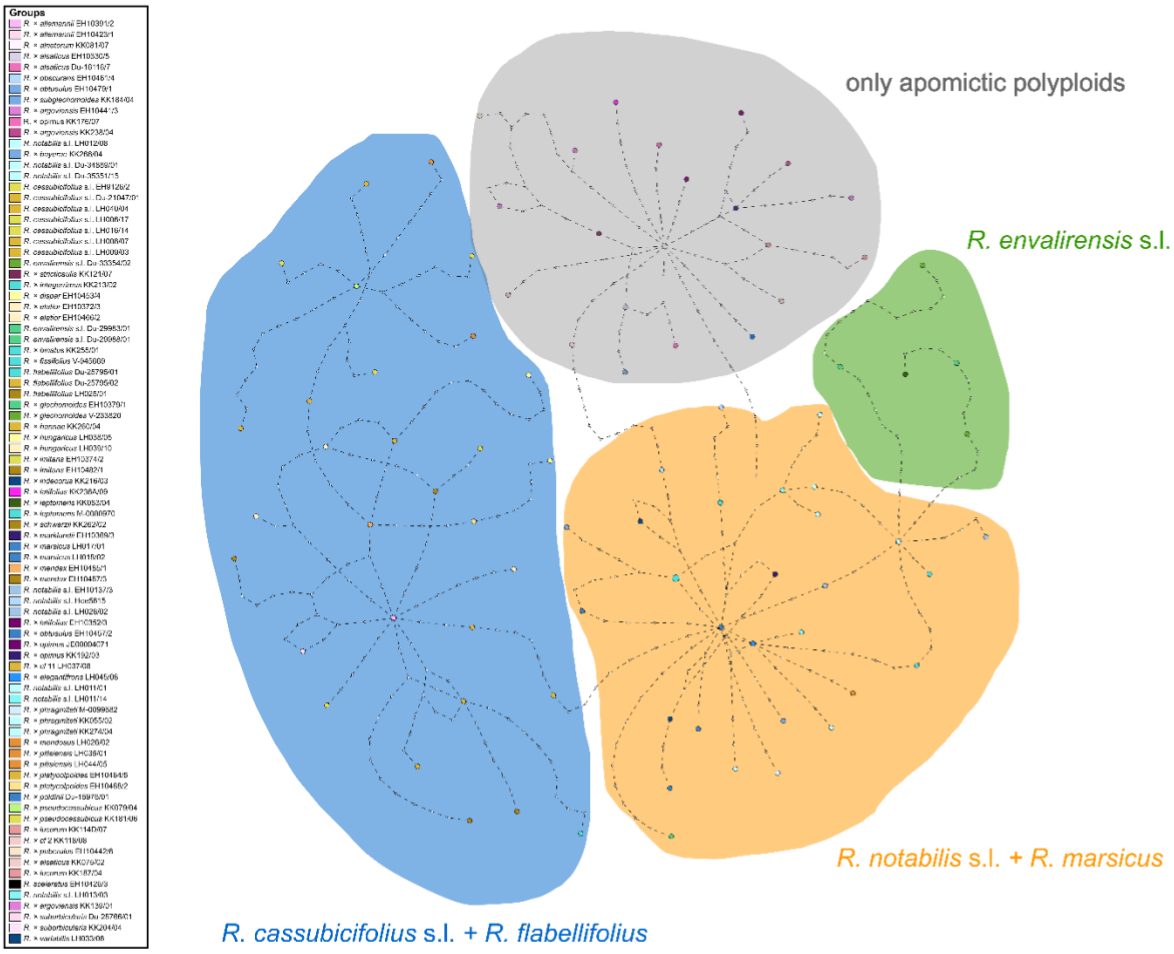


**Supplementary Fig. S14.** Haplotype network (TCS) of CP data. Results are based on 87 samples and 71 plastid regions. We colored main clusters according to Fig. 5a.

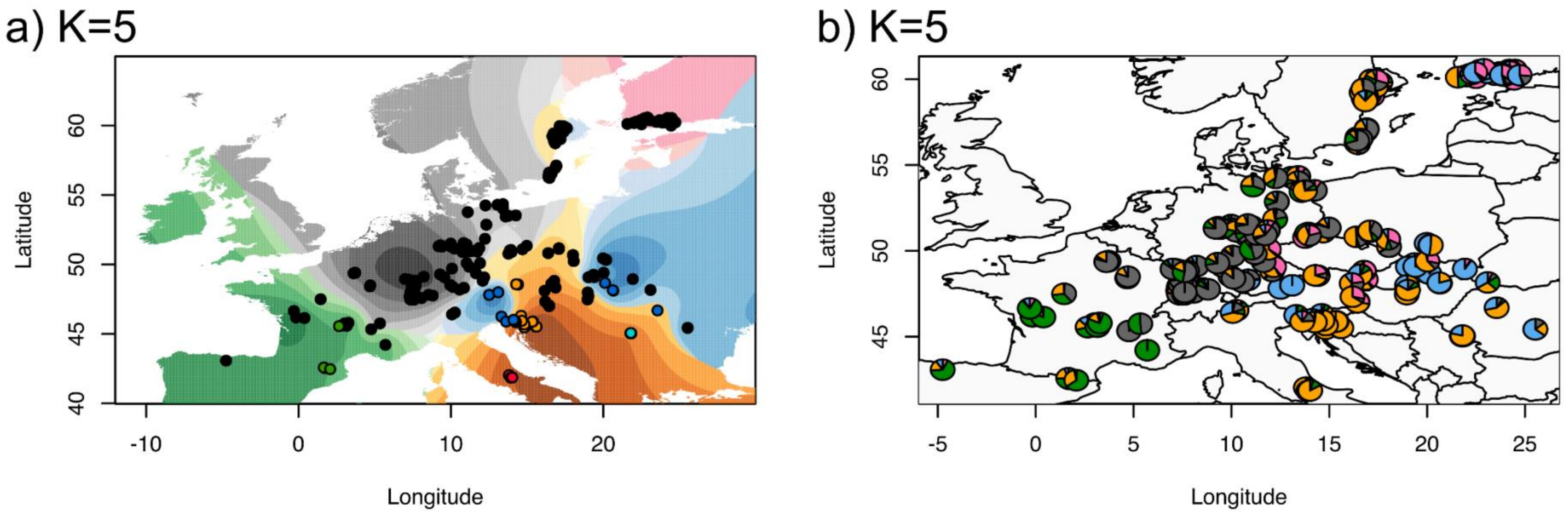


**Supplementary Fig. S15.** Geographic maps illustrating ancestry coefficients across Europe (K=5). sNMF results are based on K=5 genetic clusters, 280 sexual and apomictic *R. auricomus* individuals, and the 'min30' unlinked-SNP RAD-Seq alignment (33,165 loci). (a) Interpolated values of ancestry coefficients were illustrated with method 'max', i.e., at each point the cluster for which the ancestry coefficient is maximal and (b) location-wise admixture estimate pie charts were shown. See Figs. S11-16 and Figshare for more detailed sNMF results. In (a), colored circles represent sexual species (coloring according to Fig. 4): blue=*R. cassubicifolius* s.l. (C), turquoise=*R. flabellifolius* (F), red=*R. marsicus* (M), green=*R. envalirensis* s.l. (E), and orange=*R. notabilis* s.l (N). Europe map source: https://maps.ngdc.noaa.gov.

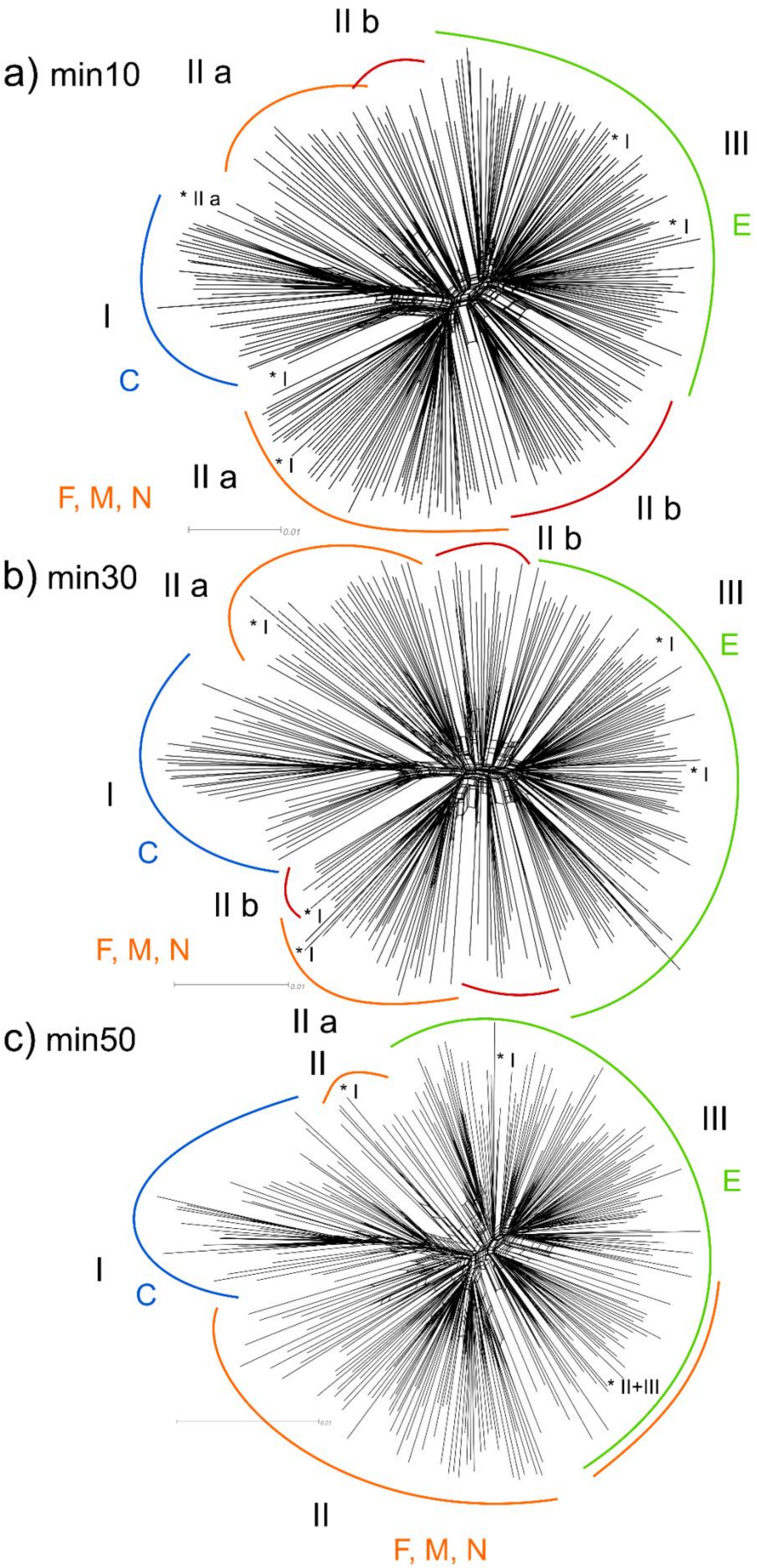

a) min10
II a
II b
III
E
* I
* I
* II a
I
C
* I
* I
II a
F, M, N
II b
0.01
b) min30
II a
II b
III
E
* I
* I
* I
I
C
II b
* I
* I
F, M, N
0.01
c) min50
II a
II
* I
* I
III
E
I
C
* II+III
0.01
II
F, M, N

**Supplementary Fig. S16.** Neighbor-net analysis (SplitsTree) of RAD-Seq data. We included 280 sexual and apomictic *R. auricomus* individuals. Results are based on genetic distances (GTR model with estimated site frequencies and ML) and the unlinked-SNP-RAD-Seq datasets, a) min10 (97,312 loci), b) min30 (33,165 loci), and c) min50 (11,196 loci). Major clusters were highlighted with colors according to Fig. 4 and designated with numbers, according to clusters found in the respective RADpainter analyses (min10, min30, and min50). Superscript asterisks ($^{*}$) show exemplary exceptions to RADpainter clusters. ‘min10’: cluster I contains *R. flabellifolius* samples that are situated within RADpainter cluster IIa; cluster III contains *R.* × *pseudocassubicus* and *R.* × *megacarpus* samples that are situated within RADpainter cluster I; cluster IIa contains *R.* × *platycolpoides* that is situated within RADpainter cluster I. ‘min30’: cluster III contains *R.* × *pseudocassubicus* and *R.* × *megacarpus* samples that are situated within RADpainter cluster I; cluster IIa contains *R.* × *platycolpoides*, *R.* × *czywczynensis*, and *R.* × *dispar* that are situated within RADpainter cluster I. ‘min50’: cluster III contains *R.* × *pseudocassubicus* and *R.* × *megacarpus* samples that are situated within RADpainter cluster I, a subcluster that contains a mixture of RADpainter clusters II + III; cluster IIa contains *R.* × *platycolpoides* that is situated within RADpainter cluster I. Colored letters indicate sexual species and also dominant subgenomes of each group, i.e., C=(subgenome of) *R. cassubicifolius* s.l., E=(subgenome of) *R. envalirensis* s.l., and F, M, N=(subgenomes of) *R. flabellifolius*, *R. marsicus*, and *R. notabilis* s.l.. See Fig. S10 (min30) in Karbstein et al. (2021) and data on Figshare for files in better resolution. Scale bar = no. of changes.

**Supplementary Table S7.** Genetic Structure and Phylogenetic Network results of tested tetraploid *R. auricomus* accessions ($H_1$-$H_{10}$). Each row ($H_1$-$H_{10}$) represents a separately analyzed individual. Results are based on RAD-Seq (RADpainter+fineRADstructure, PhyloNetworks) and TE (STACEY, PhyloNetworks, Phylonet) datasets. $P_1$ is always the parent with the largest/likeliest genomic contribution (either indicated by coancestry/posterior probability values of structure analyses or by inheritance probabilities of network analysis) followed by the other parental contributions. Genetic structure analyses (RADpainter+fineRADstructure, STACEY): Parents in brackets indicate genomic contributions below specific thresholds (i.e., only some sexual accessions of a species show significant subgenome contributions, and not the mean of all accessions, see Material and methods for details). For phylogenetic networks, percentages of subgenome contribution are given in brackets. Criteria for building consensus results of parental subgenome contribution(s) are: (i) take the most abundant parent within column; (ii) if there are two equal abundant parents (e.g., two-times 'C' and 'F') within a column, both parental subgenome contributions were taken for the consensus result ('C/F'); (iii) if there are two parental subgenome contributions for one parent, we included them with a value of '0.5' (instead of '1.0') in consensus calculations; (iv) parental subgenome contributions in brackets were ignored for consensus calculations.  The row 'final result' indicate the final subgenome contribution(s), i.e., consensus results corrected by the full likelihood approach followed by AIC calculations in Phylonet (likel+AIC, AIC = Akaike Information Criterion; different results if AIC network difference was less than 10 units) and plastome analysis results (CP type; C/F= plastid type shared by the diploid sexual species *R. cassubicifolius* and *R. flabellifolius*, * = not the same sample between CP analyses and phylogenetic network analyses, U# = haplotype from an unknown/extinct sexual progenitor species of Central Europe). Parental subgenomes not inferred by both analyses were removed from final results. Concerning the final results of $H_9$, we classified $P_1$ as

"E(U)" because of the *R. envalirensis*-like U plastid type (see above). According to final results, we classified the genome evolution of investigated and the number of involved subgenomes in polyploid formation. See also Figs. 5a,b, 8a-h, and data on Figshare for sample IDs, genetic structure, and phylogenetic network results. Letters indicate sexual progenitor subgenomes, i.e., C=*R. cassubicifolius* s.l., F=*R. flabellifolius*, N=*R. notabilis* s.l., E=*R. envalirensis* s.l., and U=probably extinct sexual progenitor.

| **Analysis** | **$H_1$ ('*R. × platycolpoides*')** | | | **$H_2$ ('*R. × elatior*')** | | | | **$H_3$ ('*R. × pseudocassubicus*')** | | | | **$H_4$ ('*R. × hungaricus*)** | | | | **$H_5$ ('*R. × fissifolius*')** | | |
|---|---|---|---|---|---|---|---|---|---|---|---|---|---|---|---|---|---|---|
| | **$P_1$** | **$P_2$** | **$P_3$** | **$P_1$** | **$P_2$** | **$P_3$** | **$P_4$** | **$P_1$** | **$P_2$** | **$P_3$** | **$P_4$** | **$P_1$** | **$P_2$** | **$P_3$** | **$P_4$** | **$P_1$** | **$P_2$** | **$P_3$** |
| **R+F** (RAD-Seq) | C | N | F | C | F | (N) | (E) | C | (N) | | | C | F | N | | E | N | (F) |
| **STACEY** (TE) | C | F | N | C | F | (N) | (E) | | E | F | N | F | C | N | (E) | E | F | N |
| **Phylo-Networks** (RAD-Seq) | C (63) | N (37) | | C (97) | E (3) | | | C (51) | E (49) | | | C (88) | F (12) | | | N (54) | E (46) | |
| **Phylo-Networks** (TE) | C (99) | E (1) | | C (68) | F (32) | | | C (63( | F (37) | | | C (51) | F (49) | | | F (61) | E (39) | |
| **PhyloNet** (TE) | E (64) | C (36) | | C (78) | C (22) | | | C (60) | C (40) | | | C (96) | C (4) | | | E (88) | F (12) | |
| **consensus** | C | N | F | C | F | | | C | E | F | N | C | F | N | | E | E<br>F | N |
| **likel+AIC** (PhyloNet) | reticulate (N, C) | | | reticulate (C, F) | | | | reticulate C=E) | | | | reticulate (F, C > C, C) | | | | reticulate (3x E, F) | | |
| **CP type** | C/F* | | | C/F* | | | | C/F* | | | | C/F* | | | | N* | | |
| **final results** | C | N | | C | F | | | C | E | | | C | F | | | E | F | N |
| **genome evolution** | allopolyploid | | | allopolyploid | | | | allopolyploid | | | | allopolyploid | | | | allopolyploid | | |
| **no. sub-genome/s** | 2 | | | 2 | | | | 2 | | | | 2 | | | | 3 | | |

| **Analysis** | **$H_6$ ('*R. × glechomoides*')** | | | **$H_7$ ('*R. × pilisiensis*')** | | | | **$H_8$ ('*R. × indecorus*')** | | | | **$H_9$ ('*R. × subglechomoides*')** | | | | **$H_{10}$ ('*R. × leptomeris*')** | | |
|---|---|---|---|---|---|---|---|---|---|---|---|---|---|---|---|---|---|---|
| | **$P_1$** | **$P_2$** | **$P_3$** | **$P_1$** | **$P_2$** | **$P_3$** | **$P_4$** | **$P_1$** | **$P_2$** | **$P_3$** | **$P_4$** | **$P_1$** | **$P_2$** | **$P_3$** | **$P_4$** | **$P_1$** | **$P_2$** | **$P_3$** |
| **R+F** (RAD-Seq) | N | F | (E) | F | N | C | (E) | N | F | (C) | | E | | | | E | (N) | (F) |
| **STACEY** (TE) | E | F | N | F | C | N | E | N | C | F | E | E | C | F | N | E | F | N |
| **Phylo-Networks** (RAD-Seq) | C (71) | F (29) | | N (67) | F (33) | | | N (92( | F (8) | | | E (82) | C (18) | | | N (60) E | E (40) | |
| **Phylo-Networks** (TE) | F (93) | E (7) | | F (91) | E (9) N | | | E (70) | C (30) | | | F (53) | E (47) | | | E (90) | F (10) | |
| **PhyloNet** (TE) | E (72) | F (28) | | E (96) N | F (4) | | | E (80) | C (20) | | | F (53) | E (47) | | | F (67) | E (33) | |
| **consensus** | E | F | N | F | F | C<br>N | E | N | C | F | E | E | C<br>F | F | N | E | E<br>F | N |
| **likel+AIC** (PhyloNet) | reticulate (E, F) | | | tree-like (F) | | | | reticulate (E, C > N, C) | | | | reticulate (E, F) > tree like (E) | | | | reticulate (E, F) | | |
| **CP type** | E | | | C*/F* | | | | N | | | | U# | | | | E | | |
| **final results** | E | F | | F | | | | N | C | E | | E(U) | (F) | | | E | F | |
| **genome evolution** | allopolyploid | | | autopolyploid | | | | allopolyploid | | | | allo- vs. autopolyploid | | | | allopolyploid | | |
| **no. sub-genome/s** | 2 | | | 1 | | | | 3 | | | | 1-2 | | | | 2 | | |

**Supplementary Table S8.** SNP discovery based on RAD-Seq SNiPLOID results. SNPs of different categories are shown in percents, and numbers of SNP positions are given (in total, without the category ‘others’). SNP percents of Cat1-5 were calculated based on all SNP positions without the category ‘others’. In brackets, we gave SNPs percent concerning all SNPs (see Material and Methods for evaluation of SNP category ‘others’). The category gives the percent of non-defined SNPs (i.e., heterozygous SNP calls for DIPLOID2) in relation to all SNP positions. Cat 1=SNPs identical to DIPLOID2, cat 2=SNPs identical to DIPLOID1/reference, cat 3/4=derived SNPs, cat 5=homeo-SNPs (hybrid heterozygous for homeologous alleles of both parental genomes; see also Materials and Wagner et al. (2020)). Concerning $H_6$ and $H_8$, we calculated two SNiPloid analyses because three parents have contributed to its origin. Coloring is according to Fig. 4. Subgenomes of C=*R. cassubicifolius* s.l., F=*R. flabellifolius*, E=*R. envalirensis* s.l., and N=*R. notabilis* s.l.

| **Analysis** | **$H_1$** (‘*R. platycolpoides*’) | | **$H_2$** (‘*R. elatior*’) | | **$H_3$** (‘*R. pseudocassubicus*’ ) | | **$H_4$** (‘*R. hungaricus*’) | | **$H_5$** (‘*R. fissifolius*’) | | | | **$H_6$** (‘*R. glechomoides*’) | | **$H_8$** (‘*R. indecorus*’) | | | | **$H_{10}$** (‘*R. leptomeris*’) | |
|---|---|---|---|---|---|---|---|---|---|---|---|---|---|---|---|---|---|---|---|---|
| | D1(REF)= C | D2= N | D1(REF)= C | D2= F | D1(REF)=C | D2=E | D1(REF)= C | D2= F | D1(REF)= E | D2= F | D1(REF)= C | D2= N | D1(REF)= E | D2= F | D1(REF)= N | D2= C | D1(REF)= N | D2= E | D1(REF)= E | D2= F |
| **Cat 1 & 2** (interspecific SNPs) | 73.65 (71.50) | | 84.51 (82.57) | | 64.35 (62.48) | | 65.34 (62.99) | | 91.46 (89.07) | | 92.53 (88.43) | | 91.6 (89.27) | | 86.94 (84.22) | | 85.08 (82.39) | | 87.47 (84.09) | |
| **Cat 3/4** (derived SNPs) | 3.08 (2.14) | | 3.86 (2.58) | | 2.81 (1.77) | | 2.63 (1.82) | | 2.78 (1.99) | | 4.67 (1.72) | | 2.72 (1.88) | | 4.84 (2.14) | | 4.41 (2.64) | | 3.08 (2.16) | |
| **Cat 5** (homeo-SNPs) | 23.26 (26.36) | | 11.64 (14.84) | | 32.84 (35.76) | | 32.04 (35.18) | | 5.76 (8.94) | | 2.80 (9.84) | | 5.69 (8.85) | | 8.22 (13.64) | | 10.03 (14.68) | | 9.45 (13.75) | |
| **others** to all SNP positions (not considered | 31.94% (2.31) | | 34.73 (2.62) | | 39.14 (2.72) | | 32.00 (2.05) | | 29.92 (1.97) | | 64.47 (3.75) | | 32.11 (1.96) | | 57.50 (4.02) | | 41.67 (2.60) | | 31.49 (2.18) | |

| | | | | | | | | | | |
|---|---|---|---|---|---|---|---|---|---|---|
| in calculations) | | | | | | | | | | |
| **SNP positions** (without others) | 5133 (7368) | 1633 (2438) | 8650 (13826) | 6208 (8947) | 3700 (5178) | 4214 (11431) | 2831 (4090) | 2665 (6019) | 2493 (4163) | 5872 (8384) |
| **SNP positions** (all) | 7542 (7542) | 2502 (2502) | 14212 (14212) | 9130 (9130) | 5280 (5280) | 11860 (11860) | 4170 (4170) | 6271 (6271) | 4274 (4274) | 8571 (8571) |
